# Phylogeny and biogeography of the algal DMS-releasing enzyme

**DOI:** 10.1101/2022.12.01.518734

**Authors:** Adva Shemi, Shifra Ben-Dor, Ron Rotkopf, Orly Dym, Assaf Vardi

**Author notes:** These authors contributed equally to this work (Adva Shemi, Shifra Ben-Dor). **Author Contributions:** A.S. and A.V. conceptualized this study and wrote the manuscript. S.B.D. identified DLHs, defined sequence motifs and analyzed the DL phylogeny. A.S. analyzed the geographic distribution, taxonomy and environmental drivers of DLH expression in the Tara Oceans dataset and created the figures. S.B.D, A.S. and R.R. analyzed the dinoflagellate DLH expression from the METZYME dataset. R.R. conducted statistical analysis and normalization of expression data. O.D. created the structural predictions for DL orthologs.

## Abstract

Phytoplankton produce the volatile dimethyl sulfide (DMS), an important infochemical, which is emitted to the atmosphere and affecting the global climate. Albeit the enzymatic source for DMS in eukaryotes was elucidated, namely a DMSP lyase (DL) called Alma1, we still lack basic knowledge regarding its taxonomy and biogeographic distribution. We defined unique sequence motifs which enable the identification of DL homologs (DLHs) in model systems and environmental populations. We used these motifs to predict DLHs in diverse algae by analyzing hundreds of genomic and transcriptomic sequences from model systems under stress conditions and from environmental samples. Our findings show that the DL enzyme is more taxonomically widespread than previously thought, as it is encoded by known algal taxa as haptophytes and dinoflagellates, but also by chlorophytes, pelagophytes and diatoms, which were conventionally considered to lack the DL enzyme. By exploring the *Tara* Oceans database, we showed that DLHs are widespread across the oceans and are predominantly expressed by dinoflagellates. Certain dinoflagellate DLHs were differentially expressed between the euphotic and mesopelagic zones, suggesting a functional specialization and an involvement in the metabolic plasticity of mixotrophic dinoflagellates. In specific regions as the Southern Ocean, DLH expression by haptophytes and diatoms was correlated with environmental drivers such as nutrient availability. The expanded repertoire of putative DL enzymes from diverse microbial origins and geographic niches suggests new potential players in the marine sulfur cycle and provides a foundation to study the cellular function in marine microbes.

## Introduction

Phytoplankton and marine bacteria produce the climatic gas dimethyl sulfide (DMS), a volatile organosulfur commonly known as the smell of the ocean(1). The DMS flux from the ocean to the atmosphere is estimated as ∼ 27 Tg S per year(2), making the oceans the main source of atmospheric DMS. Once emitted, DMS is being oxidized to sulfate aerosols(3), thus promoting cloud formation and increasing the albedo of Earth(4). DMS also serves as a ubiquitous infochemical (a chemical cue that conveys information), mediating diverse trophic interactions including predator-prey interaction(5, 6), pathogenicity(7) and habitat selection(8).

The cellular precursor of DMS is dimethylsulfoniopropionate (DMSP), an osmolyte and a cryoprotectant which is found in high cellular concentration in dinoflagellates and haptophytes, and in lower cellular concentration in diatoms(9, 10). DMSP is catabolized to DMS by distinct enzymes in bacteria and eukaryotes. The bacterial *ddd* (DMSP-dependent DMS) genes such as *dddL, dddD* and *dddP*, are diverse and are highly different from their eukaryotic counterparts(11). In eukaryotes, a DMSP lyase (DL) enzyme called Alma1, isolated from *Emiliania huxleyi* (haptophyta), cleaves DMSP to produce DMS and acrylate. Alma1 is a redox-sensitive enzyme and a member of the Asp/Glu/hydantoin racemase superfamily(1). This superfamily consists of mainly bacterial or archaeal enzymes, which catalyze the conversion of enantiomers of amino acids and play important roles in the biosynthesis of peptidoglycans and other bacterial polymers. Asp/Glu/hydantoin racemase genes are commonly transferred to eukaryotes by horizontal gene transfer (HGT), but their biological role in the receiving organisms is currently unknown(12).

The current study provides a new perspective on the ecological significance of the eukaryotic DL enzyme and its metabolic product, DMS, in marine ecosystems. As DMS is a climatic volatile, and in light of recent elevated emissions which were linked to the current global warming(13, 14), it is crucial to deeply understand the molecular basis for DMS production. Although the ecological role of the DMS-generating enzyme in the marine sulfur cycle is widely recognized, we still lack basic knowledge regarding its cellular function(1). Likewise, the taxonomy and geographic distribution of the DL enzyme are understudied, which hinders our ability to understand which species are the main players in the sulfur cycle and to predict how environmental conditions will control their DMS production. Following the recent discovery of the algal DMSP lyase enzyme(1), we are now able to address these key questions. We present an extensive mapping of the phylogeny and biogeographic prevalence of algal DL homologs (DLHs) by analyzing hundreds of genomic and transcriptomic sequences from model systems under diverse stress conditions in the lab and from global environmental samples. Our findings show that the DL enzyme is more taxonomically widespread than previously thought, as it is encoded by known algal taxa as haptophytes and dinoflagellates, but also by chlorophytes, pelagophyte and diatoms, which were conventionally considered to lack DL. By exploring the *Tara* Oceans gene dataset(15), we demonstrate that DLH mRNA expression is occurring in diverse oceanic provinces, predominantly by dinoflagellates. Haptophytes and diatoms DLHs are enriched in specific ‘hotspots’, with correlation to nutrient availability. The updated taxonomy and environmental distribution of the DL enzyme opens a new avenue to investigate its yet unknown biological role(s) in phytoplankton.

## Results

### Domain architecture of DLHs and their identification in diverse algal species

In order to investigate the phylogeny of the DL enzyme, we utilized the amino acid sequences from two orthologues which have high DL enzymatic activity (∼600-800 µM DMS hour^-1^): *E. huxleyi* (herein Alma1) and *Symbiodinium* A1 (herein Sym-Alma)(1). Alma1 and Sym-Alma have similar modeled 3D structures, except Sym-Alma has three cysteine residues located at the active site of the enzyme, as compared with only two cysteines in Alma1(1) (Fig. S1). We used the Alma1 and Sym-Alma amino acid sequences, which are quite different (26% identity), as query in several databases and published genomes (see Methods), and identified a total of 153 eukaryotic DLHs (Datasets S1, S2). DLHs were defined according to their Asp/Glu/hydantoin racemase domain (cl00518 in the CDD database, herein ‘racemase domain’(16)), and two canonical cysteine residues (C108 and C265 in Alma1), which are critical for DL activity(1) (Fig. S1). These cysteines are located in the active site and catalyze epimerisation by proton abstraction and addition. Accordingly, mutations in these residues compromised activity(1). Racemase domain duplication occurs in 33% of identified DLHs, and a DEP domain (Dishevelled, Egl-10 and Pleckstrin domain, cl02442) is conserved in several homologs (Fig. S2). Protein alignment of 151 racemase domains from DLHs and 34 bacterial sequences belonging to the closest racemase subfamily showed a clear difference between eukaryotic and prokaryotic homologs (which do not possess DL activity, PRK07475 in the CDD), which suggests different functions (Fig. S3). We also defined ten conserved and discriminatory sequence motifs by comparative analysis(17) between the racemase domain of all DLHs with all the racemase superfamily available in the CDD database (which are bacterial sequences, see Fig. 1 and Table S1). Following confirmation of the racemase domain, the canonical cysteines, and the unique motifs, we defined 110 DLHs in 41 photosynthetic and one heterotrophic protist (*Crypthecodinium cohnii*, dataset S1, S2 and Fig. S4). Lastly, corals are the only animals suggested so far to encode DLHs, which expanded to gene families in some species. The coral DLHs that were defined here were not studied in depth, as they were recently described elsewhere(18, 19).

**Fig. 1.**
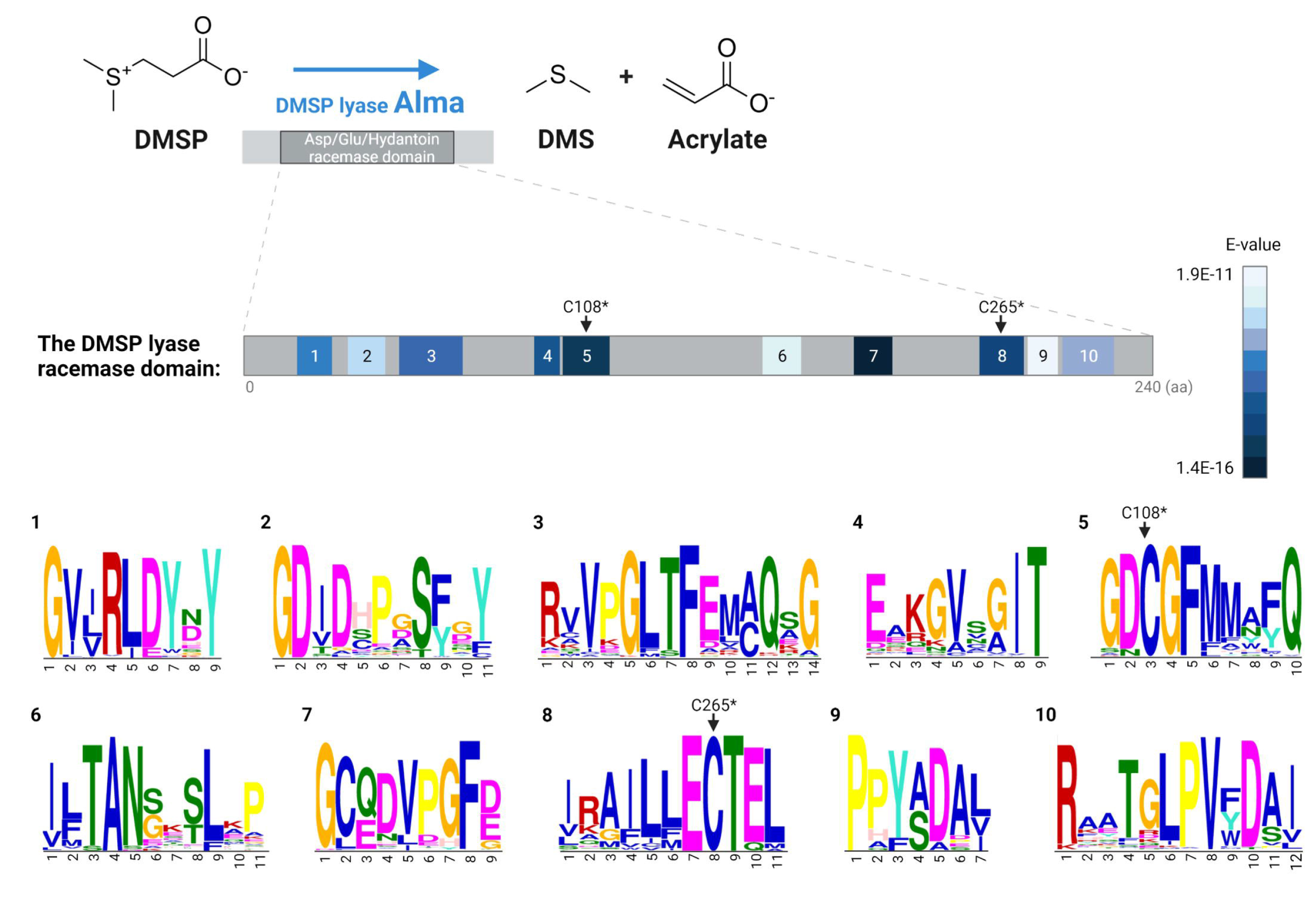
Conserved domain and sequence motifs in the DMSP lyase (DL) protein. **A** The DL enzyme Alma1 cleaves DMSP to produce DMS and acrylate. It has an Asp/Glu/hydantoin conserved domain, which contains ten unique amino acid motifs. The motifs are numbered according to their order in the sequence and are colored according to their E-value score. The E-value represents how the sequence is significantly shared between 151 DLHs and also discriminant from 917 bacterial racemase domain. **B** The sequence logos of the ten identified conserved motifs, numbered as in (**A**). *According to the *E. huxleyi* Alma1 sequence(1).

### Phylogeny of DMSP lyase homologs (DLHs)

Marine phytoplankton evolved through multiple endosymbiosis events. Primary endosymbiosis gave rise to glaucophytes, rhodophytes (‘red algae’) and chlorophytes (‘green algae’), and secondary endosymbiosis gave rise to chromalveolates comprising dinoflagellates, haptophytes and bacillariophyceae (herein, diatoms)(20). Photoautotrophs originated from primary endosymbiosis (including land plants, which evolved from a chlorophyte ancestor) are generally considered to lack DL enzymes, with the exception of green macroalga *Ulva mutabilis* (sea lettuce), which has two predicted DLHs(21). Other *Ulva* species were shown to produce DMS, but their putative DLHs remain to be identified, since they are still missing sequenced genomes(22). Here, six DLHs were predicted in *Ulva prolifera*, and new DLHs were defined in green microalgae including: *Cymbomonas tetramitiformis, Tetraselmis striata, Tetraselmis suecica*, and *Platymonas subcordiformis*, with the latter previously shown to synthesize DMSP(23) (Dataset S1). Targeted searches for DLHs in additional genomes derived from the green lineage (including plants), as well as in all available glaucophytes and rhodophytes genomes did not yield any significant putative homologs, according to the above-mentioned criteria (Dataset S3).

Photoautotrophs originated from secondary endosymbiosis include major DMS/P producers such as haptophytes and dinoflagellates(1). We previously identified DLHs in bloom-forming haptophytes as *Prymnesium polylepis* (formerly called *Chrysochromulina polylepis*), *Prymnesium parvum, Isochrysis* sp., *Phaeocystis antarctica* and *E. huxleyi*(1) (Table S2). Here, new homologs were identified for *Isochrysis* sp. and *P. antarctica* (A total of 4 and 8 homologs, respectively). In addition, two DLHs were identified in a new haptophyte genome, *Ochrosphaera neopolitana* (Dataset S1). In contrast to the apparent gene families predicted in those haptophytes, no DLHs were detected in other haptophyte genomes, suggesting that DL conservation is species-specific and not a general haptophyte trait (as *Isochrysis galbana*, Dataset S3). Therefore, reports for DMS-production in some haptophytes lacking DL may be derived from an alternative enzymatic source, which is yet to be discovered, or their associated bacteria(24).

In dinoflagellates, more than nine species were previously described to encode for DLHs, including *Symbiodinium* and the toxic, bloom-forming *Karenia brevis* and *Alexandrium temarense*(1, 19). We expand DL gene families in most species by identifying new members, with up to 11 DLHs in *G. foliaceum* (Dataset S1). DLHs were also identified in nine new dinoflagellates: *Cladocopium goreaui, Amoebophrya ceratii, Heterocaspa triquetra*, and six *Symbiodinium* species. The various *Symbiodinium* species also have multiple DL family members. Note that *S. microadriaticum* strain CCMP2467 has sequences identical to the ones previously identified (*Symbiodinium* sp. A1)(1). Like haptophytes, the DL gene is not common to all dinoflagellates but encoded by specific dinoflagellate genomes (Dataset S1, S3).

Unlike haptophytes and dinoflagellates, diatoms are considered to have a low cellular DMSP pool(25), and were not reported so far to have DL enzymatic activity. We identified putative DLHs in eight ecologically important diatoms, including four *Skeletonema* species, a cosmopolitan bloom-forming diatoms(26); *Seminavis robusta*(27); *Attheya*; and the bloom-forming *Pseudo-nitzschia multiseries* and *Pseudo-nitzschia fradulenta* (Fig. 2, Dataset S1). In addition, three putative DLHs were defined in a related group named pelagophyceae (pelagophyceae sp. CCMP2097). A phylogenetic tree of the DL protein was constructed by aligning and comparing the racemase domain of 173 DL sequences from 34 eukaryotic and 20 bacterial species (Dataset S2). The bacterial Alma1-like homologs were clearly different form their eukaryotic counterparts and appeared in the tree as an out-group (Fig. 2 and Fig. S5). The general distribution of the species in the tree differs from their taxonomy, which may be the result of HGT from eukaryotic or bacterial sources(12).

**Fig. 2.**
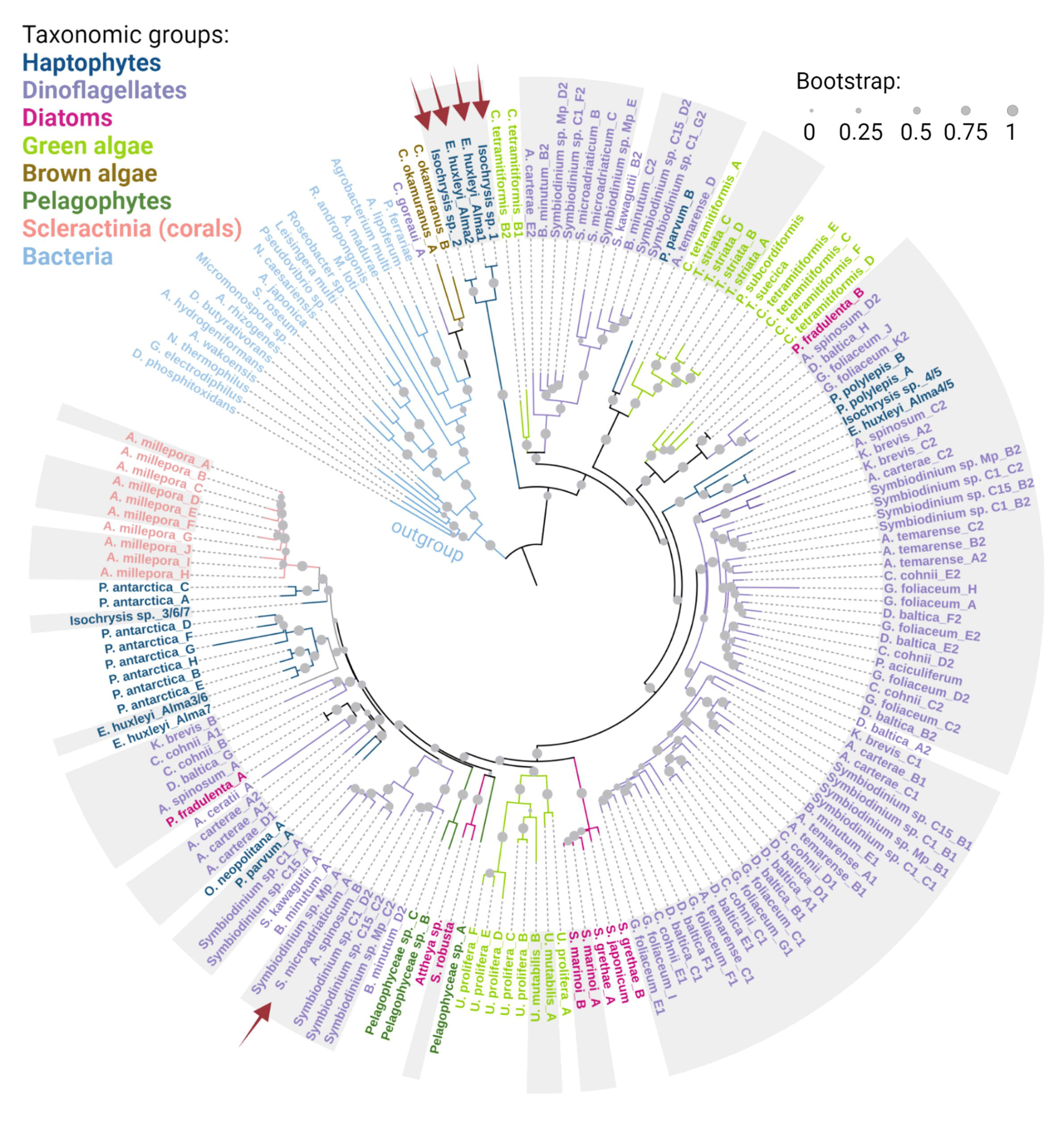
Phylogenetic distribution of the DMSP lyase enzyme. A phylogenetic tree of identified DLHs, colored according to their taxonomy. Grey background designates experimental evidence for mRNA expression (see Tables S3, S4). Red arrows indicate biochemical confirmation of DL activity in the marked homologs. The Isochrysis sp. orthologs are ≥99% identical to alma1 from *E. huxleyi* and are therefore confirmed by identity(1).

In order to validate the expression of predicted DLs under environmental conditions and to gain insights into their possible physiological function, we searched available algal transcriptomes for DLH expression in datasets generated from cells exposed to diverse environmental stress conditions)28(. Transcription of most DLHs was confirmed, including differential expression in some species in response to environmental stressors such as high light or nutrient limitation (Tables S3, S4). For example, *S. dohrnii* and *S. robusta* DLHs expression was ∼8-9 fold less upon nitrogen depletion. In addition, in *Isochrysis* sp., expression of one out of four identified DLHs (DLH C) was increased ∼300-fold in response to high light (Table S4). Interestingly, DLH expression was species-specific under diverse environmental conditions. Future biochemical studies are required to determine DL enzymatic activity of these predicted homologs and to validate their physiological role under these stress conditions.

### Global biogeographic distribution of DMSP lyase homologs in the ocean

To date, atmospheric and seawater DMS concentrations are measured on a global scale(2), yet investigation of the algal origin of DMS formation is restricted to specific ecological niches or bloom events(29, 30). Here, the global biogeography of DLHs was examined using the *Tara* Oceans dataset, a collection of pan-oceanic plankton metagenomes and metatranscriptomes(15). When using the DL sequence from *Symbiodinium* (Sym-Alma) as query (which is more similar to other DLHs, as compared to Alma1, see Dataset S1) high abundance of eukaryotic DLHs was detected in both metagenomes and metatranscriptomes databases (Marine Atlas of Tara Ocean Unigenes, MATOU(31); Fig. 3 and Fig. S6). Transcripts homologous to DL were detected in all 581 samples collected from surface, the deep chlorophyl maxima and, unexpectedly, in mesophotic depths. DLHs transcripts were widely distributed across the oceans, with highest abundances in the Norwegian Sea (station *Tara*_163), Barents Sea (*Tara*_168), South Atlantic Ocean near Argentina (*Tara*_82) and Antarctica (*Tara*_85), and the North Pacific Ocean near Hawaii (*Tara*_131 and 132) (Fig. 3A). These locations include nutrient-enriched areas with seasonal blooms of DMS-producers such as *Phaeocystis(32)*. To assess whether DL transcription level is a proxy for DL production in the field, 17 stations for which DMS concentration was previously reported by independent scientific cruises were selected (Table S5). Indeed, DL homolog expression was positively correlated with the DMS levels reported in the same locations, suggesting that DLH transcript abundance is a good proxy for DL activity in the environment (*r* = 0.7386, *p* = 0.0010, Fig. 3B and Table S5). To characterize the taxonomy of DLHs, we focused on stations with high transcript abundance (Fig. 3C). Oceanic DLHs were mainly from dinoflagellates such as gonyaulacales (for example: *Azadinium* and *Alexandrium*), suessiales (*Symbiodinium*), peridiniales, gymnodiniales and others. Stations *Tara*_82 and 163 were enriched with haptophyte DLHs belong to noelaerhabdaceae (*E. huxleyi*), phaeocystaceae (*P. antarctica*), prymnesiaceae (*Prymnesium*), isochrysidaceae (*Isochrysis*) and *Haptolina*. Furthermore, stations *Tara*_173, 175, 188 and 189 were enriched with DLHs from diatoms, mainly fragilariophycidae (*Thalassiothrix antarctica*), thalassiosirophycidae (*Thalassiosira antarctica*) and chaetocerotophycidae (*Chaetoceros*) (Fig. 3C and Fig. S7). Interestingly, heterotrophs such as radiolarians, ciliates and metazoans were also associated with DLH expression, which can be derived from their potential algal symbionts (Fig. 3C, “other”)(33, 34).

**Fig. 3.**
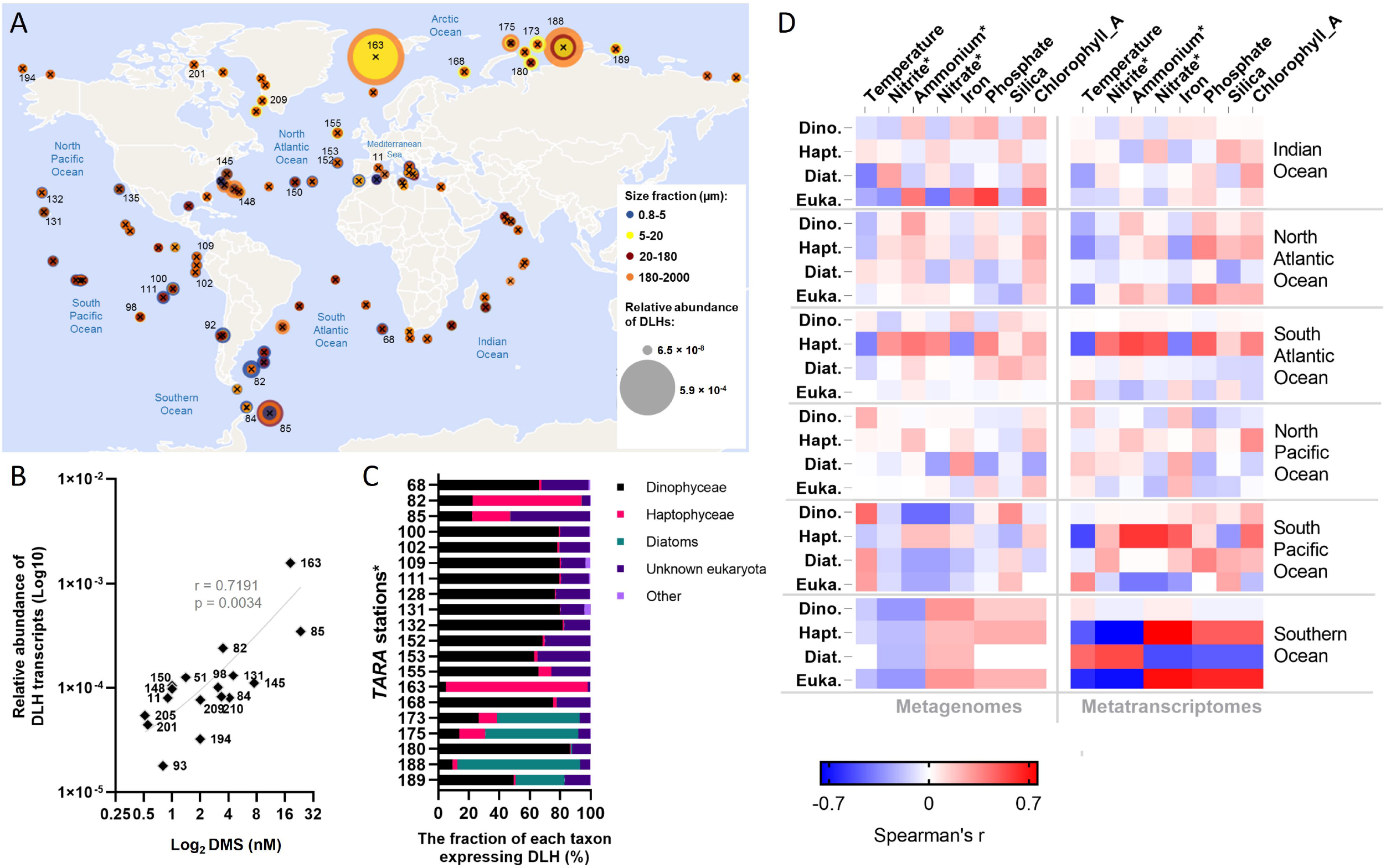
Biogeographic distribution of DMSP lyase homologs (DLHs) in the oceans. **A** The relative abundance of DLHs in metatranscriptomes from 86 locations worldwide, in samples collected from surface water (5 m). The circles are proportional to transcript relative abundance (normalized as percent of mapped reads) and colored according to the different size fractions collected for each sample. The numbers designate *Tara* stations which were further analyzed in (**B-C**). **B** Correlation between DLHs mRNA expression level in 17 *Tara* station (labels correspond to station no.) and the DMS concentration in the surface water (as reported by independent studies summarized in Table S5). Spearman’s r = 0.7191, *p* = 0.0034, n = 17. **C** Taxonomic distribution of DLHs in the *top 20 stations with the highest DLHs expression, as described in (**A**). **D** Correlation between environmental conditions and total DLH abundance in 316 DNA samples (left panel) and 401 RNA samples (right panel). Pairwise comparisons are shown with a color gradient denoting Spearman’s correlation coefficient. Comparisons were done for per taxa and per each oceanic region. For metagenomes, n = 60 for Indian; 83 for North Atlantic; 65 for South Atlantic; 64 for North Pacific; 32 for the South Pacific; and 12 for the Southern Ocean. For metatranscriptomes, n = 66 for Indian and North Pacific; 95 for North Atlantic; 71 for South Atlantic; 90 for South Pacific; and 14 for the Southern Ocean. The DLHs abundance and taxonomic distribution were based on the 0.8-2000 µm size fraction from all depths available in the *Tara* Oceans dataset. Dino.=Dinophyceae, Hapt.=Haptophyceae, Diat.=Diatoms, Euka.=Other eukaryotes. *Estimated values based on oceanographic models(15, 60).

To further investigate the environmental drivers of DL mRNA expression, which may imply on the physiological role for DL, we analyzed the correlation between abiotic factors measured in each station (temperature and nutrients) and the relative abundance of DLH sequences in all metatranscriptomes versus metagenomes (Fig. 3D). Dinoflagellate DLHs mRNA were generally highly expressed across the oceans, regardless of the environmental parameters tested. In the South Pacific and Southern Oceans, haptophyte DLHs mRNA expression (mainly *Phaeocystis*) was positively correlated with nutrient availability (nitrogen, phosphate, or iron). This contrasts with diatom-derived DLHs expression in the Southern Ocean, which correlated with low nutrients, suggesting that the DL cellular role under stress conditions may be taxon-specific.

Dinoflagellates are unique phytoplankton since they inhabit the sunlit as well as dark deep ocean, owing to their versatile metabolism which alternates between photoautotrophy and heterotrophy (also known as mixotrophy(35)). As dinoflagellate DLHs expression was not affected by the environmental parameters measured during the *Tara* campaign, we asked whether they differentially express DL in surface samples versus deep water samples. Indeed, a remarkable abundance of dinoflagellate DLH transcripts was detected in the deep dark ocean (200-740 m) in various locations (Fig. S8). We further explore this phenomenon in a metatranscriptomic study presenting a detailed vertical profile of dinoflagellate populations from the surface to the depth (800 m) of the central Pacific Ocean (the METZYME transect(36)). In addition to the light gradient, deep water was rich in nitrogen, which was gradually depleted toward the surface. Dinoflagellate transcripts were prevalent in the euphotic (≤200 m depth) as well as the mesopelagic (>200 m) zones. Among 25 distinct dinoflagellate DLHs identified, 16 homologs were significantly expressed in the sunlit, nitrogen-limited zone. Interestingly, four homologs were significantly upregulated in the dark, nitrogen-replete zone (two from *H. triquetra*, one from *Gambierdiscus polynesiensis* and one from *D. baltica*, similarity >83%, Fig. S9). Taken together, the differential expression of dinoflagellate DLHs between the euphotic and mesopelagic zones suggests functional specialization of DLHs even in the same taxa, and may imply its role in the metabolic plasticity in mixotrophy in dinoflagellates(36). In summary, DLH transcript abundance in natural populations was correlated to nutrient and/or light availability in a taxon-specific manner, thus suggesting a function for the DL enzyme as part of nutrient and carbon metabolism and the recycling machinery in phytoplankton.

## Discussion

Oceanic DMS is an important component of the marine sulfur cycle, with ecological implications ranging from chemical signaling across trophic levels to the global climate. Here, we investigate the phylogeny and geographic distribution of the eukaryotic DMS-releasing enzyme, the DL Alma1(1). DLHs were detected in the genomes of dinoflagellates, haptophytes, diatoms, chlorophytes, pelagophytes, and scleractinians. The expanded repertoire of putative DL enzymes from diverse microbial origins suggests new potential players in the marine sulfur cycle beyond the known representatives from dinoflagellates and haptophytes. We revealed that DLHs are species-specific and are encoded in the genomes of distantly related taxa (Dataset S1, Fig. 2). We could not detect DLH in red algal genomes, including the seaweed *Polysiphonia* from which DMS was historically detected for the first time(37), which may be attributed to alternative enzymatic sources or to strain variability (according to available genomes). Therefore, DL enzymes in chromalveolates may have originated from a green ancestor, or from the heterotrophic host or red alga (prior to gene loss). In addition, the distribution of DLHs in the phylogenetic tree suggests that HGTs from prokaryote and eukaryote genomes may played a role during the evolution of the DL enzyme(12) (Fig. 2 and 4A).

**Fig. 4.**
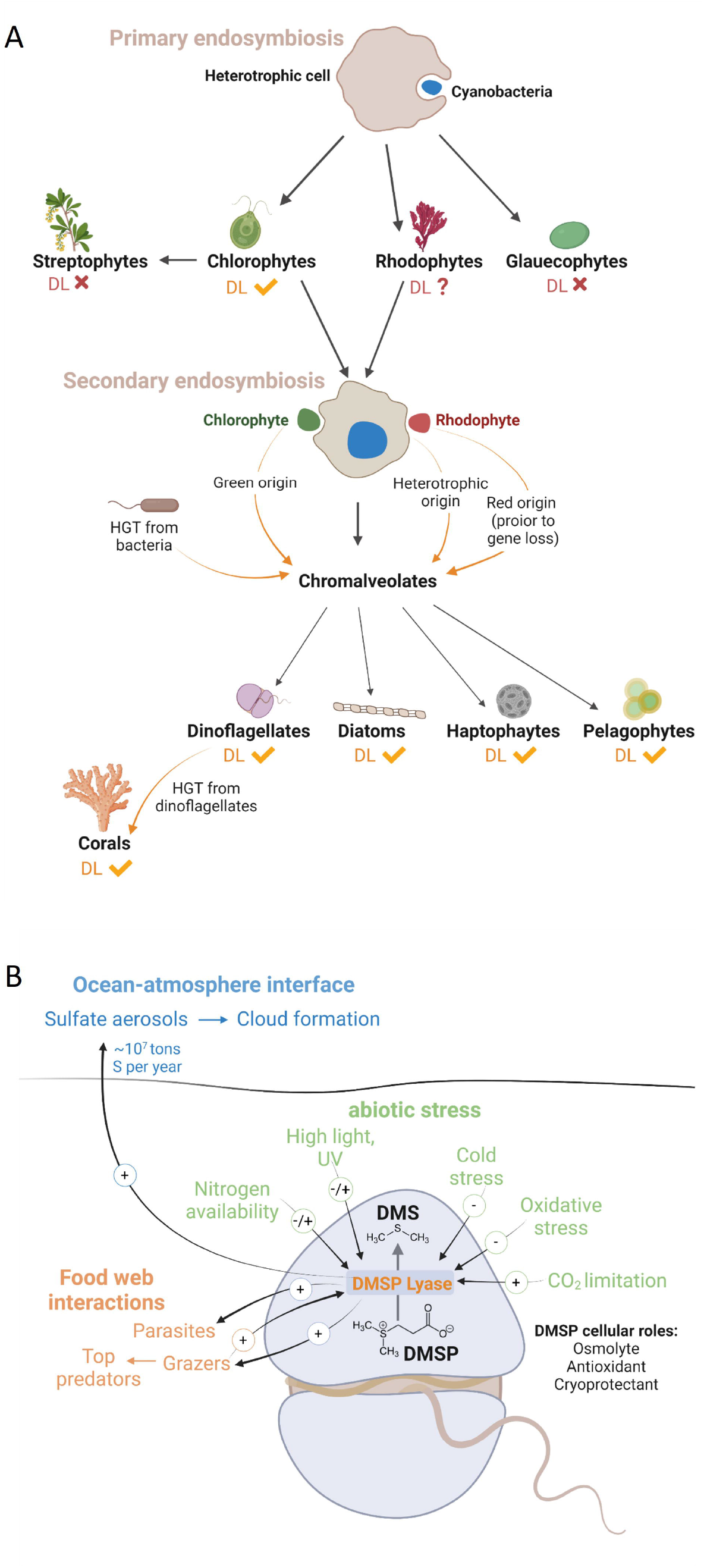
The evolution, cellular and ecological roles for the DMSP lyase enzyme in phytoplankton. **A** Possible acquisition pathways for the DL enzyme in chromalveolate algae. Primary endosymbiosis gave rise to chlorophytes, including species which possess DLHs, and to rhodophytes and glaucophytes, which their DLH are either missing or remained to be identified. Secondary endosymbiosis gave rise to chromalveolates as haptophytes, dinoflagellates, diatoms and pelagophytes, which all have representatives encoding for DLHs. The DL evolutionary origin in chromalveolates may be attributed to the heterotrophic host that engulfed a rhodophyte or chlorophyte, to a rhodophyte (considering unidentified DL, or that DL gene loss in red genomes occurred after secondary endosymbiosis), or to chlorophytes or bacteria via horizontal gene transfer (HGT). Corals DLHs were probably horizontally transferred from their chromalveolates symbionts. **B** Proposed cellular and ecological roles for the DL enzyme under abiotic stress. DL enzymatic activity was linked (up- or down-regulated, depending on tested species) to nitrogen levels (this study, and Ref. (27, 43)); high light and ultraviolet (UV) radiation (this study, and Ref. (39, 41, 42)); cold stress(21); carbon dioxide(39), and oxidate stress(1). During microbial interactions, DMS act as an activation cue for parasites(7) and plays a pro-grazing role for protistian grazers(5) and copepods(46). DMS is also accumulated in the water during grazing, serving as chemoattractant for diverse top-predators(61). In addition to the mentioned cellular and ecological roles, DMS plays a climatic role; the ocean-atmosphere DMS flux is estimated as 10 tons S per year(2), and atmospheric DMS is rapidly oxidized to form sulfate aerosols, which enhanced cloud formation and precipitation.

The prevalence of DLHs in diverse, but highly selective, algal genomes propose important cellular role(s) for this enzyme. Unlike DMSP, which is a known osmolyte(38), cryoprotectant, antioxidant(39), and energy dissipation metabolite(40), we know very little regarding the biological function of the DL enzyme. Does the DL activity act merely in controlling the cellular DMSP levels? Does DMS play a cellular role as well? Previous studies had shown that temperature, high light(41, 42), carbon dioxide(39) and nitrogen(43) modulate DMS production in different phytoplankton, but the DMS role is still not fully understood (Fig. 4B). Such environmental stress conditions can induce cellular reactive oxygen species (ROS) levels, which can be a mechanism for DL activation or inhibition, as it is a redox-sensitive enzyme(1). Furthermore, DMS can scavenge ROS by being oxidized to dimethyl sulfoxide, thus acting as a potent antioxidant and promoting cellular resistance to stress(39, 44).

DLH expression by most taxa was detected across the oceans, with dinoflagellates emerging as the dominant taxa expressing DLHs. Dinoflagellate DLHs mRNAs were even detected in mesopelagic depths, implying a possible unexpected metabolic role for DLH in mixotrophy(36). Another possibility is that DMS generation by those mixotrophs is a strategy to attract algal or bacterial prey(45). Haptophytes, diatoms and others expressed DLHs in correlation with some nutrient availability in specific oceanic provinces, suggesting again a link between DMS/P metabolism and nutrient metabolism which is rarely studied(43) (Fig. 4A). This potential vital role for the DL enzyme under ecologically-relevant conditions is especially interesting in light of its wide evolutionary distribution comes with a cost, since DL activity in phytoplankton can strongly enhance predation by zooplankton(5, 46) (Fig. 4B). Consequently, there should be high negative selection to retain this DMS-generating enzyme in algal genomes. It is plausible that tradeoff between abiotic stress conditions (such as nutrient limitation) and grazing pressure promotes the selective retention of the DL enzyme in phytoplankton, including ecologically successful species such as *E. huxleyi* and *Symbiodinium* (Fig. 4B).

Elucidating the DL function in its physiological context and during microbial interactions poses great methodological challenges. Future research should confirm DL enzymatic activity in cultured species, which will extend the repertoire of model organisms available to study the DL physiological role in phytoplankton. To date, specific *in vitro* DMS production was demonstrated by expression and purification of recombinant DL enzymes only for *E. huxleyi* and *Symbiodinium* A1(1). In *Phaeocystis, Ulva* and others, DMS was detected in cell extract and not from a specific gene product(1, 47, 48). Interestingly, the reef-building coral *Acropora* encodes a set of genes similar to Alma1 and Sym-Alma (∼43% and 60% similarity in the amino acid sequence, respectively. Dataset S1 and(19)). DLH genes in corals were predicted to be horizontally transferred to the coral genome from its dinoflagellate symbiont(18), and their enzymatic activity remains to be fully validated(49). However, the absence of genetic tools to interrogate gene function in dominant DMS-producing species is a major bottleneck to investigating the ecophysiological role of the DL enzyme. The current study provides the basis for developing model systems and experimental setups needed to interrogate the cellular role of the DL enzyme in ecologically relevant phytoplankton. In the current study, we identified DLHs in several model algal species that are amenable to genetic transformation as *P. multiseries, T. striata, C. cohnii, A. carterae*(50) and *Ulva*(51). In addition, applying a selective DL inhibitor(49) during growth experiments under stress conditions and in the course of biological interactions(5) could shed light on DL function.

With the apparent impact of the global climate change and DMS emissions(13, 14), there is a critical need to understand the molecular, biochemical and cellular origin of DMS in the ocean. Our findings have expanded the taxonomy, biogeographic distribution and potential cellular roles attributed to the DL enzyme and provide new genomic resources and environmental insights that will help to unravel the ecological importance of this unique enzyme in the oceanic ecosystem.

## Materials and Methods

### Predicted protein structures of DL orthologs

In order to predict the structure of the DL enzyme, a BLAT(52) search was conducted against the Protein Data Bank (PDB) using the DL sequences from *Symbiodinium* (Sym-Alma) and *E. huxleyi* (Alma1) as queries. Since no significant similarity to any known protein structure was found, theoretical structural models were created using the AlphaFold server(53, 54). Ribbon representation (Fig. S1) was created using PyMOL (The PyMOL Molecular Graphics System, v. 1.2r3pre, Schrödinger, LLC).

### Identification of DLHs

In order to identify DLHs, database searching was performed at various websites using BLAST(52), either BLASTP or TBLASTN, with three sequences as initial input: *E. huxleyi* Alma 1 (AKO62592), *Symbiodinium* (P0DN22) and *Acropora millepora* (XP_029211597). Data sources used for searching DLHs include the Marine Microbial Eukaryotic Transcriptome Sequencing Project (MMETSP)(28); the Joint Genome Institution (JGI); Supplemental data from Nelson *et al*., 2020(55); and various genomes and TSA in NCBI. For red algal genomes, the ‘Red Algal Resources to Promote Integrative Research in Algal Genomics’ Blast site(56) was used. For *A. millepora*, the draft genome (v. 2.01) published by the Przeworski lab(57) was used. When a sequences from species close to the target had already been defined, those sequences were also utilized as queries in the searches. For downloaded genomes and datasets, local BLAST(52) was performed (v. 2.11.0). Genes were constructed manually as described in Feldmesser *et al*., 2014(58). Sequences found in the initial search were then studied further, and considered as DLHs only if they contained both canonical cysteines at the active site (C108 and C265 from *E. huxleyi* Alma1)(1) and the surrounding sequences (DCGF or NCGF and ECTE or ECTQ, respectively). Domain analysis was performed on the sequences using the Batch CD-Search (CDD v3.19) at NCBI(16). Only sequences with a hit to Asp/Glu/Hydantoin racemase superfamily domain (cl00518) were retained.

### DL sequence motifs and logos

To define unique sequence motifs for the DL enzyme, differential motifs in the amino acid sequence were analyzed with Streme (v. 5.4.1), a part of the MEME suite, using 151 DLH sequences without bacterial or putative bacterial sequences from the alignment. A total of 917 bacterial sequences from the racemase domain family in the CDD were used as control. Default parameters were applied except for logo length, where 4-15 positions were allowed. Logos of the alignment were constructed with WebLogo (v. 3.0).

### DLHs sequence alignments and phylogenetic tree

In order to align the identified DLHs for phylogenetic analysis, sequences were cut to the domain (according to the CDD search), and alignments were performed using Muscle (v. 3.8.31) and ClustalW (v. 2.1)(59). The final alignment (Muscle) for further analysis was cleaned of extensions by removal of all columns with less than 10 sequences. The alignment was used as input for similarity/identity calculation using MacVector (v. 18.0). Phylogenetic tree was constructed using Neighbor Joining in ClustalW with 1000 bootstraps (v. 2.1) and Maximum Likelihood with the default parameters in PhyML (v. 3.0). The PhyML trees were visualized with iTol (v. 6).

### Transcription of DLHs in cultures phytoplankton

To validate the transcription of predicted DLHs, mRNA expression data was taken from published datasets. The MMETSP database(28) was used for expression levels in cultured species, in response to abiotic stress conditions (Tables S3, S4). Additional transcriptomic studies are referenced in Table S4. Read counts were normalized with Deseq2.

### Biogeography of DLHs in the Tara Ocean dataset

In order to map the bio-geographic distribution of DLHs in the marine environment, the Ocean Gene Atlas platform was used(15, 60). DLHs search was performed with tblastn, utilizing *Sym*-Alma (from *Symbiodinium* A1, (1)) as query. The databases queried were the Tara Oceans Eukaryotic Genomes (SMAGs) and the Marine Atlas of Tara Oceans Unigenes (MATOU_v1_metaG, eukaryotes) for metagenomes, and the MATOU_v2_metaT for metatranscriptomes. The expected threshold was 1E-10. The abundance was calculated as present of mapped reads.

### Dinoflagellate DLHs expression from the METZYME transect

To compare dinoflagellates DLHs expression between euphotic and mesophotic depths, we searched for DLHs in the metatranscriptomic dataset published by Cohen et al.(36), using the Blast criteria listed above (TBlastN, conserved cysteines and surrounding sequences). The putative DLH expression was then extracted from the dataset.

Illustrations were created with BioRender.com.

Graphs were created with GraphPad Prism.

## Acknowledgments

We thank Dr. Gust Bilcke for providing the DLH mRNA expression data for *S. robust*a (Table S4). We thank Dr. Uria Alcolombri for his contribution to scientific discussions and his constructive feedback on the manuscript. This work was funded by the research grant from the Israeli Science Foundation (ISF No. 1972/20) and by a research grant from the Takiff Family Foundation both were awarded to A.V.

## Competing Interests Statement

The authors declare no conflict of interest.

## Data Availability Statement

The data used in this study is available in datasets S1-S3.

## Notes

### Competing Interest Statement

The authors have declared no competing interest.

