## Supplemental dataset 2 for "Phylogeny and biogeography of the algal DMS-releasing enzyme"

### Dataset S2. Amino acid sequences of predicted DL homologs.

<u>Prymnesiophyceae</u>

>P.polylepis_A HBOG01061503.1
MNAGSKQYLEKFGSKIEGAICDALAATIRAEPDDPMKFLQSWFMDNAQGGSSVLQERQFLEHQLARSKHRLDELSTL
MPTIGVLRTERAGYSSAWLELSGEGDNSYMYKIVEQEVAGLSYEHVRAGKPLSAEQADALKIAIKQLEAQGAVAICGD
CGSFVHYQQLAVKLTTLPVVLSPLLQAPLLAAMYPVSEKFLVITNDSRSIDQATMRANLLKAGLAEANADRFVLVGLEH
IDGFVENALTHGKSLRQLDRLEQQLIETVERKQKADPALRAVLLESTLMPEFSDSLRYRKRLPVFDSITIADFIHAATSP
NPRFGTADPRFPEQYKSEALKTRWDSLDKEKMPTIGVLRIDYSYPPAPGDIDHPSSYYYRTTQRTSKGLTFEAAQEGE
PLTAEQRREMQAAIAELETHNVVGITGDCGFLMNYQIDARRIAKVPCFISSMLQCPLLANFFDGEEHILVCTANGKSLA
PRIGTMLTACGVPAAQQSRFIVVGCEDVDGFDAVEKGEKVDVARVQPGIIDLVQREMAAAGRKVRAIMLECTELPPYA
DALRAATGLPVLDAITLVDFFQSAISDNPYFGIDFQRSSQRLSTAVEERRRESRAYSMAKIVE

>P.polylepis_B HBOG01030655.1
MGCCGSKTNDKPFEPIVTPDLDGEKIAVAGTVTAGAAIGAAIAGPAAPIGALVGAAVGAAVAKSTAGTVGKVIVAPIGLL
KDVWRGLLNRPEILDQLPDIGMLRSERKVYPPSMSQLTETNSYLYQIHPATVTGLKFEEALEGKPLSDTQRVALADAIK
VLEAKNCKAITGDCSAMLHYQEEVLRMTKLPVLLSALLQAPLLAASYSKDEKILVLTSDATVTSPAKMAAALSKCGLAA
EDLDRFITVGLETIDGFKASSLASGAATIDPNETAQQLLKIVQDAKASTPGLRAVLFESTMLPMFSDLLRKELKLPVFDV
VTLADLVHNAQTDNPRFGVAFGDKTSSAAPTLRLDKGQMPAIGIMRIDYTYPPAMGDAAHPNSYYYRTPHATVTGLTF
EAAQKGDPLTPAQATAMEAAVKTLEADPGLMGIAGDCGFLVNYQAHAVKLATRVPLFISSVLQCSLLTNIFGPDSQVL
VLTANGPELQRVLPNMLSLVNVPAKDHGRFVVAGCESLPGFEAVALAQKVDVDQVQPHCVALVQAKLAEHPMIRAVL
LECTELPPYADAIREATGLVVMDVITLVDFFHGAVSEDPYFGIDWDKIASTPVGRGAGSSSS

>P.parvum_A HBJC01021092.1
MSHVASTAKTMTEYVTSQALNATKAKEAPVRAKDGTLGIVRLDYNYPPAPGDIDHPASFAYDVFYKVVPGLTFEMCQ
SGKLTPEVEERFLNTIKYFEAKGVSGITGDCGFMMYFQALARQATNKPVFMSALAQLPAVTAAFGNEELIAIFTANGST
LKPMEPMICDECGVNVEEQRYVIVGCEDVPHFEAVARGEKVDVDKVTPGMIAKARDVLATYPNIRAILLECTELPPYAD
ALRKETGLPVYDAITACDFFMMGVQDNERFGLQDWQVVWDGQQEEYTFAQNLTEEEKKHLVNKVS

>P.parvum_B HBJC01045452.1
MQRSVEQGKVARSQLLLRHTIQRRLAMEHHGDLGDDLYELLYAKPKPAEERRAEPRHAVALAGDRHDAKAVRPLGVI
RLDYGYPPIPGDIDHPSSFDYPVVYRKVPGLLFEVAQEGVLTPEIRSGIEQAVRELEAHQVFGITGDCGFMANYQHLV
RSLASCPVFLSSLVLLPAISPLIGSRDRILCCTANGNSLAKLLPALWGAPPPPAAAYGVTHTFTHKHLDLHLLRPEQLVV
CGFERVPGFDVVAEASSLMSHAVDRELVEQGITSMVLTILSQDPTIKMILLECTELPHYTVTLRQATGLPVYDSLSICNF
VHAAYDVSPSEASHRSRCLGHHCQIKYPDTPTTG

>P.antarctica_A
MLPALRRVAPRLTRTANVGVRGLSKGTQATNPKLGIIRLDYDYVAAPGDIDHPGSFGYEVVYRVVPGLSFEMAQEGT
MSPDVKTRFIEAIKYLEAEGVNGYTGDCGFMMNFQQLARTNTKLPVYMSSLCQLPSVTCAYGQNEKIAIFTANGVSLA
PMRDLIRDECGVDTEQERYIIVGCENVDHFGDEVAKGLKVNTDLATPGIVKKALETQAANPEIRAFLFECTELPQFSDA
VRAATGLPVYDSITMSDSFMSGVTDNKRFGLNDWQGEFDGKQADYKFGDNLTDEQKENIEYKDMVKVPPLARQGPS
SPPAPPRGAPGGFGQLETPMKRPAHSAPSHSLGYLS

>P.antarctica_B
MNLHRHASLGVVRLDYDYPPAPGDIDHPGSFGYDVFYRVVPGLTFAMCQSGEMPPEIEERFTNAVKWLDAKGVSCIT
GDCGFFMNFQHIARRVTHKPVIMSSLCALPSIVCAYAKHERIAIFTANGVSLQPMRQLLRDECGVDPEDQRFVIVGCQ
DVPGFEAVALGERVDVEKCTPHIVKLAQDIVAKHAGTSTPIRAIVFECTELPPYSDAVRAATKLPVCDAITTCNSFVALM
RDNPRFGLDNWHVSWDGKQEEYRFGQNLTSDMKELLVNDEHAENTPNDEHAEDKPFLPYVARPGRGPHRNAGVA
ENMR

>P.antarctica_C

GKMTDEVEARYKDAIKYLEAEGVSGITGDCGFMMYFQPLARTLTKLPVYKSSLCQLPSVTCCYGQTEKIAIFTANGKSL
APMRDLIRDECGVDTEQERYIIVGCEDIPHFGEEVAAGDKVDTDKATPGMVQKALEVQKANPDIRAFLFECTELPQFS
DAVRAATGLPVYDSMTMSDSFMSGVMNNKRFGKQDWQAEFDGKQVDYKFGDDLTDAERKEVEYTKLDDAN

>P.antarctica_D
MLRSLRLTRPLAAKHLRACSLSTKPKKAAATLGVVRLDYEYPPALGDIDHPGSYSYDVFYRVVPGLTFAMCQNAVPKT
GTPLPDDVMKSFVEAIKYLDQDKNVSGITADCGFFFNLQDEARKHTKKPVFMSSLCAAPTVAAAYAKDEQIAIFTANKA
HLDPMSAEINEACGIALEDKRFVIVDCLTVPGFGHAVDVGDRVDLAAATPNIVKLALDTVKANPKLRAIIFECTELPPYS
DAVREATGLPVFDSITTSNAFMASMQDNPLFGANDWQEKWDGKQDSYHFGDNLTAEQKAKITWQTKK

>P.antarctica_E
MLLKRPVGGAKEALEKAGFCVAKRRGPNNGGPNDDKPEPWFSPKQKPPASPAVRKARATREYIAANVPGASASWV
AGSVANQLIPDEEGLQLHRHASLGVVRLDFDYPPAPGDIDHPGSFGYDVFYRVVPGLTFAMCESGNMPPEIEERFTN
AVKWLDAKGVSTITGDCGFFMNFQHIARRVTHKPVIMSTLCALPSITCAYAKTERIAIFTASKETLHPMRQLLWDECGV
DPEDKRFVIVGCQDVPGFETVAQGERVDVQECMSHIVKLAQDVVAEHAETDSPIRAIVFECTELPPYSDAVRAATKLP
VFDAITTCNSFVASMRNNPRFGVDNWHVSWDSKHDDYRFGEFLSSEMKEKLVNDERPGAPGRSQGVHVSM

>P.antarctica_F
MLRSLRLTRPLAAKHLRACSLSSKPKRPAASLGVVRLDYNYPSAAGDIDDPRSYAYPVFFTGDCGYFFNLQHEARKH
TKKPVFMSSLCAAPTVAAAYAKDELIAIFTANKAKLEPMRDLIKEECGIDPEDKRFIIVDCRYVEDFGDAVDKGDKVDTV
KATPNIVKLAEKTLQENPKLRAIIMECTELPPYSDAVRAATGLPVFDSITTSNAFMASMQDNPLFGANDWQEKWDGVQ
EKYDLGDNLSAEEKAKLGNAK

>P.antarctica_G
TGDCGFMLSLQEEARKHTKKPIFMSSLCQAPSVTNLFDRDELIAIFTSDAAALTRMLPLIKEQCRIDVGEERFIIVDCSKV
DGFGIEIEKGLQVNVEEVSPLMVALAKKIVQEDHDRQADHKLRAIIFECTELPPYSDAVRHATGLPVFDSITASNAIVES
MVDNPRFGKTGWQEAWDQSQEDYEFGQKLSKEEKERLLN

>P.antarctica_H
MLRSLRLTRPLAAKHLRACSLSTKGKKAAASLGVVRLDYDYPAAPGDIDSPESYGYDVFYRAVPGLSFEMCQHAVPK
TGTPLPDDVMKGFVEAIKYLDQEKNVSGITADCGFFFNLQHEARKHTKKPVFMSSLCMAPTVAAAYAKDELIAIFTANS
ASLEPMRDLIKEECGIDPNDKRFVVVGCQDVPIFGKAVAEGTKVDWEAVTPHMVKLAQDTVKANPALRAIILECTELPP
YSDALRAATGLPVFDSITTSNAFIASMQDNPRFGANDWQENWDGEQDEYKLGQNLSAEDKAKLQSK

>Alma1 AKO62592.1 Alma1 DMSP lyase [Emiliania huxleyi]
MGNCTSHPHHEPQVHFDTVDGMATVKAFAGIRQVTMGVLRIDYEYQTNLGDILDPRSFDFRIISATAEGLTFAKAKAG
DKLDATGKELLERAVRQLIDNGADFIVGDCGFLVYWQVMVRDYAQDYAQKKYGRKCPVMMSSLVLALPLLATIPSGG
QIGILTASEKSLQAVQKKLPIVIEDHQKEDGGQRSRSVPAAEDPSGIMINFSDPRFKVVGLDEVKDFKHALDAQGADAV
NDRRDIAVQIAAYCQKVQEKNPQIAAWLIECTEAGGFAWAIKVGTGLPVWDPITLGRFLSLGFTANVPNVALTLGQHG
ENPLDPSATTAGKGRCTGEEPGQIHALGAEFQAIREGTL

>Alma2 AKO62593.1 Alma2 DMSP lyase [Emiliania huxleyi]
MGSASSKTRKDKAKSSTTAACPTDTAAAKACPTRLDDGLAQVAMFAGLRQVTMGVLRIDYDYQTNLGDILDPRSFDF
RLVSATVEGLTFKRAQEGEPLPCYVMSNLDGAVKKLIDAGADFIVGDCGFLVYWQVYVRDFAQQYARGRACPVMLS
SLVLSLPLLATIPVGGKIGILTASKGSLMKMQKKLASVIELQKEEARTRAVPAAVQPSGIEINFSDPRFKVVGLDTVNSFK
TALADDSGVDDRRSIAIEIAKYCKQVACEDPAICAWLIECTEAGGFSWAIKLGTGLPVWDPVTIGRFLSLGFTSSLPSVA
LTLGETGQVALDPNETDVSKGRPTKAEHRFGPEFEEMLQ

>Alma3 sp|R1G6M4.1|ALMA3_EMIHU RecName: Full=Dimethylsulfonioproprionate lyase 3; Short=DMSP lyase 3;
AltName: Full=Dimethylpropiothetin dethiomethylase 3
MGCAGSTLRSGASFEDSRLAAIEDSRFHEVGHHAQFEDSRLAAIEDSRFHEVGHHAQFDEGGRFKQLPPANDDAKLL
VADHPSLGVIRLDYDYPPALGDVDHPGSFYYDVFYRVVPGLTFELCQSGELPDDVKQRFIDAITWLDEQGVAGITGDC
GFFMYFQALARSVTSKPVFMSSLCQLPAVVCAYAADEQIALFTANGESLKPMREIIKKECGVDPDDTRFVIVGCEDVP

GFEAVANGDRVDVDSVVPHLVRLAEDTVAKHAGTAKPIRAILFECTELPPYSDAVRAATRLPVFDSITCCNSMLASLM
DNPRFGVNNWHLSWDGAHTAHRFGDNVPPHLKGKLVNREHPENIARWNASLAERSSFSSAQQESIGRGSRKL

>alma4 sp|R1ERP2.1|ALMA4_EMIHU RecName: Full=Dimethylsulfonioproprionate lyase 4; Short=DMSP lyase 4; AltName: Full=Dimethylpropiothetin dethiomethylase 4
MGCTHSKEHSTANKPPAVGILRCHGARWMPAQAVVGTNSYLFRTVHAEVKGLRYLDVRAGKELSVSQKKALKEAVK
ELDAEGVVAITGDCGSFVHYQTAVRRMTKTPAVLSPLLQAPLLATMYMKEETILVLTNDSSDYDQAALESNLVEIGLSQ
EDAERFVIQGLQHIEGFSTSEVADMSDERTWAMVDTSERLRLEIMSTIEAAKKANPSLRAILLESTLLPSFSDSMRQTR
GVPVFDAITLADYLAAASTDNPRFGSNIDASVWSSAMERANVMDELSQRATPAIGILRIDYSYPPAPGDVDYPGSYYY
RTVQEVAAGLTFEAAQEGRPLTAQQREAMEGAIRRLEAAKGVVGITGDCGFLMNYQVDARRMSHLPCFISAMMQCH
MLAASFAADEEFLVLTANGKSLAPKFGEMLSLAHVTRPEDQARFHILGCEDVDGFDAVAKGEAVDVARVTPGIVALAK
AAAARRPKVRAVLLECTELPPYADALRHALRIPVLDAITLVDFVHSASTDNPAFGVDFQKSSKVFV

>Alma5 sp|R1ENF4.1|ALMA5_EMIHU RecName: Full=Dimethylsulfonioproprionate lyase 5; Short=DMSP lyase 5; AltName: Full=Dimethylpropiothetin dethiomethylase 5
MPAQAVVGTNSYLFRTVHAEVKGLRYLDVRAGKELSVSQKKALKEAVKELDAEGVVAITGDCGSFVHYQTAVRRMTK
TPAVLSPLLQAPLLATMYMKEETILVLTNDSSDYDQAALESNLVEIGLSQEDAARFVIQGLQHIEGFSTSEVADMSDER
TWAMVDTSERLRLEIMSTIEAAKKANPSLRAILLESTLLPSFSDSMRQTRGVPVFDAITLADYLAAASTDNPRFGSNIDA
SVWSSAMERANVMDELSQRATPAIGILRIDYSYPPAPGDVDYPGSYYYRTVQEVAAGLTFEAAQEGRPLTAQQREAM
EGAIRRLEAAKGVVGITGDCGFLMNYQVDARRMSHLPCFISAMMQCHMLAASFAADEEFLVLTANGKSLAPKFGEML
SLAHVTRPEDQARFHILGCEDVDGFDAVAKGEAVDVARVTPGIVALAKAAAARRPKVRAVLLECTELPPYADALRHAL
RIPVLDAITLVDFVHSASTDNPAFGVDFQKSSKVFV

>Alma6 sp|R1F493.1|ALMA6_EMIHU RecName: Full=Dimethylsulfonioproprionate lyase 6; Short=DMSP lyase 6; AltName: Full=Dimethylpropiothetin dethiomethylase 6
MGCAGSTLRSGASFEDSRLAAIEDSRFHEVGHHAQFEDSRLAAIEDSRFHEVGHHAQFDEGGRFKQLPPANDDAKLL
VADHPSLGVIRLDYDYPPALGDVDHPGSFYYDVFYRVVPGLTFELCQSGELPDDVKQRFIDAITWLDEQGVAGITGDC
GFFMYFQALARSVTSKPVFMSSLCQLPAVVCAYAADEHIALFTANGESLKPMRELIKKECGVDPDDTRFVIVGCEDVP
GFEAVANGDRVDVDSVLPHLVRLAEDTVAKHAGTAKPIRAILFECTELPPYSDAVRAATRLPVFDSITCCNSMLASLMD
NPRFGVNNWHLSWDGAHTAHRFGDNVPPHLKGKLVNREHPENVARWNASLAERSSFSSAQQESIGRGSREL

>Alma7 sp|R1CW23.1|ALMA7_EMIHU RecName: Full=Dimethylsulfonioproprionate lyase 7; Short=DMSP lyase 7; AltName: Full=Dimethylpropiothetin dethiomethylase 7
MAGKDRKTIEKNYPGAEVDEGGRFKPLPPADDDAKLLVCEYPSLGVIRLDYDYPPALGDIDHPGSFYYDVFYRVVPGL
TFGMCQKGEMPDEIKQRFIDAIKWLDAQGVAGITSDCGFFMNFQDLARTVTDKPVFMSSLCQLPAVVCAYAAHEHIAL
FTANGESLKPMRDLIKKECGVDPEESRFIIVGCQDVPGFEAVANGDRVDVDSVMPHIVRLAKETVAKYADTAKPIRAIL
FECTELPPYSDAVRAATRLPVFDAITSCNSFLAALMDNPRFGVNNWHLSWDGSQTDYRYGDNLSADLKAKLVNAEHA
ENVAAAERKLAKDRQKPKPATGTGTAFDA

>O.neopolitana_A
MRRKLASTLGIIRLDYNYPPAPGDIDSPDSYEYDVFYRVVPGLTFEMCQSGHMTEDVKAEFQEAIRYFEQKGVCGITG
DCGFMMHFQPMARQLTKKPVFLSALAQLPAVTCSFGPKEKIAIMTANGDTLTPMQPLINSECGVDVVGEERFIIVGCQ
DVPGFEAVALGKKVDVKAVTPGMVAKAKQVLAENPDVRAFLMECTELPPYSDALRKETGLPVYDAITCSNFFMTGFQ
DNPRFGLNDWQECWDCEQHSYTFGEELNEAQRAKLVNKTNA

>O.neopolitana_B
MPSCKTVCAIAAQSFTLAPAHGQALRPVIRTTPRSALLDLRGSEGEDPKVLQGLDPDFIADLDKVNRNKAEEQKLREP
PLVGVLTLDGSNYCRDDIDETSKVKNWTLGDADNRNSYWQLGYRCLFKRVEGFTFETCKRGLSPQGPYDDINEALF
QISGTVTYPDDGKSQDFIDSHAKLGGKMRPRNGEKTEGKAEDGKTDTTYTYGNETLRANLQLALDELKARNVSVIVA
DCGFMGEIQRLCVETSKQLCLMSSVSLLPTIKTMIPQDQLIFILTANGTSFENSYEQMIRPAWGIEQSSIQLIGLQGVEG
FGLAVKDGTSVDTSLAQANIVEEVKKAMKTSAKVGCILCECTELVGFSNALRREFKDVPVFDVMSCAAFALASVSQHN
AFPADATPDEKQGRPL

>Isochrysis_sp.1 CAMPEP_0188750792 /NCGR_PEP_ID=Isochrysis-sp-CCMP1244-20130912|8451_1
/TAXON_ID=37098 /ORGANISM="Isochrysis-sp-CCMP1244" /LENGTH=328 /DNA_ID=CAMNT_0032210317
/DNA_START=1 /DNA_END=986 /DNA_ORIENTATION=-
XVKAFAGIRQVTMGVLRIDYEYQTNLGDILDPRSFDFRIISATAEGLTFAKAKAGDKLDATGKELLERAVRQLIDNGADF
IVGDCGFLVYWQVMVRDYAQDYAQKKYGRKCPVMMSSLVLALPLLATIPSGGQIGILTASEKSLQAVQKKLPIVIEDQQ
KEDGGQRSRSVPAAEDPSGIMINFSDPRFKVVGLDEVKDFKHALDAQGADAVNDRRDIAVQIAAYCQKVQEKNPQIA
AWLIECTEAGGFAWAIKVGTGLPVWDPITLGRFLSLGFTANVPNVALTLGQHGENPLDPSATTAGKGRCTGEEPGQIH
ALGAEFQAIREGTL

>Isochrysis_sp.2 CAMPEP_0188739558 /NCGR_PEP_ID=Isochrysis-sp-CCMP1244-20130912|1392_1
/TAXON_ID=37098 /ORGANISM="Isochrysis-sp-CCMP1244" /LENGTH=386 /DNA_ID=CAMNT_0032198223
/DNA_START=1 /DNA_END=1159 /DNA_ORIENTATION=+
XEQEPMGSASSKTRKDKAKSSTIAACPADTAAATACPTRLGDGLAQVAMFAGLRQVTMGVLRIDYDYQTNLGDILDP
RSFDFRLVSATVEGLTFKRAQEGEPLPCYVMSNLDGAVKKLVDAGADFIVGDCGFLVYWQVYVRDFAQQYARGRAC
PVMLSSLVLSLPLLATIPVGGKIGILTASKGSLMKMQKKLASVIELQKEEARTRAVPAAVQPSGIEINFSDPRFKVVGLDT
VNSFKTALADDSGVDDRRSIAIEIAKYCKQVACDDPAICAWLIECTEAGGFSWAIKLGTGLPVWDPVTIGRFLSLGFTS
SLPSVALTLGETGQVALDPNETDVSKGRPTKAEHRFGPEFEXDAAVRSEGVVYAMPHDPDRLVEGRPLVANQP

>Isochrysis_sp.3_6_7 CAMPEP_0188740626 /NCGR_PEP_ID=Isochrysis-sp-CCMP1244-20130912|2246_1
/TAXON_ID=37098 /ORGANISM="Isochrysis-sp-CCMP1244" /LENGTH=301 /DNA_ID=CAMNT_0032199431
/DNA_START=53 /DNA_END=958 /DNA_ORIENTATION=+
MGCGASKVAAATGEQGKKKKLKACSLGVIRLDYDYPPAPGDIDNPESFAYDVFYRVVPGLTFDMCQSGKMTPEVEE
RFIGKVKELDELGVNGITGDCGFMMFFQELARCHTKKPVFMSALCCLPALTCGYAANEKIAIFTANGETLAPMRDLIRD
ECGVDTQGERFIIVGCQDVPHFEAVALGEKVDVEKVTPGMVEKAKTVIKDNPNVRAILMECTELPPYSDAVRATMGLP
VYDAITACDFFMMGVQDNKRFGLQDWTEDWDGKQDSYSFGQELTQLEREKMVNKATVPPTPTVHEALR

>Isochrysis_sp.4_5 CAMPEP_0188770424 /NCGR_PEP_ID=Isochrysis-sp-CCMP1244-20130912|23305_1
/TAXON_ID=37098 /ORGANISM="Isochrysis-sp-CCMP1244" /LENGTH=737 /DNA_ID=CAMNT_0032231643
/DNA_START=1 /DNA_END=2210 /DNA_ORIENTATION=-
XGCTHSKEHSTANKPPAVGILRCHGARWMPAQAVVGTNSYLFRTVHAEVKGLRYLDVRAGKELSVSQKKALKEAVK
ELDAEGVVAITGDCGSFVHYQTAVRRMTKTPAVLSPLLQAPLLATMYMKEETILVLTNDSSDYDQAALESNLVEIGLSQ
EDAERFVIQGLQHIEGFSTSEVADMSDERTWAMVDTSERLRLEIMSTIESAKKANPSLRAILLESTLLPSFSDSMRQTR
GVPVFDAITLADYLAAASTDNPRFGSNIDASVWSSAMERANVMDELSQRATPAIGILRIDYSYPPAPGDVDYPGSYYY
RTVQEVAAGLTFEAAQEGRPLTAQQREAMEGAIRRLEAAKGVVGITGDCGFLMNYQVDARRMSHLPCFISAMMQCH
MLAASFAADEEFLVLTANSKSLAPKFGEMLSLAHVTRPEDQARFHILGCEDVDGFDAVAKGEAVDVARVTPGIVALAK
AAAARRPKVRAVLLECTELPPYADALRHALRIPVLDAITLVDFVHSASTDNPAFGVDXPEELKGLCLRREAAKTDGSIA
XIAASPSVKRERAXLKYKTYSCCAYMHTRFLQYNLQILRSGELKTLGVHTRGTCRPLQPLQIAESADVIVTKLQIGSSRV
ARIFVRSRRVSAGKLLDDRAAARVVTEVGGRVKYAASDDDPASKLRGVLADNAPAGLVVIGCTAAAAARGLHSSLAG
RPPPLASAVGGLEPLVLRRGSQNRRGSMLLFLQRT

Dinoflagellata

>A.carterae_A CAMPEP_0186439658 /NCGR_PEP_ID=Amphidinium-carterae-CCMP1314-20130924|22079_1
/TAXON_ID=2961 /ORGANISM="Amphidinium-carterae-CCMP1314" /LENGTH=669
/DNA_ID=CAMNT_0029243069 /DNA_START=1 /DNA_END=2010 /DNA_ORIENTATION=-
AQVTATLALKLKSELVDKHKVLPAFQKGNSDNMGCGASSTKNTEAEQKPAATSNEPPKAEEKKAEPAPAAKPAEPAK
TEEKKPEAKKEEVKKFEVDEKAEDASLGVIRLDYDYPPAKGDIDHPGSFCYDVFYRVVPGLTFEMCQSGKMTPEVEK
EFVEAVKYLEGRGVNAITGDCGFMMYFQALARQNTKLPVFMSSLAHLPAITCGYSFKEKIAIFTANSETLKPMKPLIKD
ECGIDSDESRYVIVGCQDVPGFEAVALGEKVDVAKVTPGMVKKCKEVLAEHPTIRAILMECTELPPYSDALRHVTGLP
VYDAITCCDFFINGMRDNPRFGLNDWHLEWDGVQETYKFGKHLSKEDKEKLVNKPKTEENNPKKSTLAEPAVKKQKK
AMQLMAKVRAERAHASLGVVRLDYDYPPAPGDIDSPQSYAYDVYYRVVPGLTFSMCQAGKMTPEVEKEFIEAINYLK
NEKKVSAITGDCGFMMYFQALARQHCNLPILMSSLAQLPAITCAFAKEELIAIFTANSETLTPMRALIKDECGVEPEEKR

FIIVGCQDVPGFEAVAAGEKVDVDKVTPGMVAKAKEVIAKHPTLRGILLECTELPPYADALRFSTGLPVYDSITGCDLF
MQGLMDNPRFGLNDWHEEWDGEQEAYEFGKYLDEEDRQKLVNKVA

>A.carterae_B CAMPEP_0186460378 /NCGR_PEP_ID=Amphidinium-carterae-CCMP1314-20130924|63059_1
/TAXON_ID=2961 /ORGANISM="Amphidinium-carterae-CCMP1314" /LENGTH=787
/DNA_ID=CAMNT_0029267009 /DNA_START=1 /DNA_END=2361 /DNA_ORIENTATION=+
HPAAVQYHFFGDSQGSNVMDMVPARGSSVHERLDRLEQELSKVCMVQQAPQRHPFDMPSGGALRGVGTLMMSQS
TVPPRTSGGAHSARGGDPHERLQRLEQEITSLKQGSIVAPAQTTAISTPLVLATGSTMAPKAQGMETCGNLNSLAMS
SYSPNTGYSREHLELMSAHKELLQMMKEQSKTHTELMNAHKELLKSNAKPASGRGTQDVGEKKKSKHPTLGVIRLDY
NYPPAEGDIDCPASFGYDVIFRVVPKMTFEMAQKGKFTEEVEREFAEGIKYLELQGCDAITGDCGFMMAFQVLARKIA
TKPIFMSSMVQCPVLAAAFDKSEKILILTANSTSLGPQKEVLLSSCGFDVEEDRFHIVGCENVPGFEAVAKGEKVPLDL
VTPGIVKLVTEELRKDPTITGILLECTELPPYADALRAATELPVWDAITAADFYISSRRDNPRFGINDWQAQWDGQHDE
YVFGQNLTEMDKEALVNKPTGQAAPKAMPKKAPKKQDADKLKKKLQKKQCPTLGILRLDYNYPPAKGDIDCPGSYDY
DCLFRMVPGLTFEMAQAGKMTFAVEQEFRKAIKWLEAKGASGITGDCGFMMAFQPLARQIAAVPVFMSSMMQCPLIS
VAFDKYDKILILTANSDTLLPQKENLLSNCGFDVDDDRFIIKGCQDVPGFDAVAKAQKVDVDSVTPGIVALVKDVLAKEI
STRAILLECTELPPYADALRKETGLSVFDAITNADFFISSVRDNPRFGLNQWQMDWDGVVEDYKLGENLSAQERAKAS
LFDITDKMARG

>A.carterae_C CAMPEP_0186456096 /NCGR_PEP_ID=Amphidinium-carterae-CCMP1314-20130924|57390_1
/TAXON_ID=2961 /ORGANISM="Amphidinium-carterae-CCMP1314" /LENGTH=755
/DNA_ID=CAMNT_0029262017 /DNA_START=1 /DNA_END=2267 /DNA_ORIENTATION=-
XLEPGSRSPQRAQVPQTVVTRAMYSAGGCGSPGMMQSLSMASLPPYGYDQSSLRGAGAQMMSPAGNHQHVERL
NRLEYEIMKLRDGPAMGMHGQPPPLSMSGGQQIMPVSNHYAGNMPLQLGGMMQGDALSQLMSAHTQLIQVMTKQ
GETHTELMQAHKELMNTHRQMLASGNFSGGAAKGGAAVKNKSKHPSLGVIRLDYDYPPAEGDTDCPASFGYDVFYR
VIPGLSFEMAQRGKFSEEVERRFADGIKFLEQRGVSAITGDCGFMMAFQILARKIAGKPVFMSSMVQCPVMSCAFDT
SEKMMIMTANGHSLKPQKEVLLSSCGFDVEEDRFVIVGCQDVPGFDAVAKGLEVPLDVVQPGMVKLCMETLRRHPQI
KGILLECTELPPYADAIRAATDLPVWDAITAADFYISGFRDNPRFGVNDWQQEFDDIQEEYHFGSNLVRSDQKLLENQ
GGSKGADLRKKKAIMSQNNPQVQKLKKQMKKKQFPSLGVVRLDYSYPPAAGDIDCPGSYDYDVVFRAVPGLSFEMA
QAGKMTPEVDAEFRKAIQFLEAKGVSGITGDCGFMMAFQPIARKIAKVPVFMSSMVQCPLISVAFDKYDKILILTANDLT
LKPQKETLLSHCGFDVDDDRFIIKGCQAVPGFDAVAKAGKVDVPFVTPGIVKLTKEMIQKMPSIRAILLECTELPPYADA
LRKETGLAVFDAITCADFFISARKDNPRFGLNAWQNPWDQTVDEYKLGQNLSQAERSQMQSLPS

>A.carterae_D CAMPEP_0186442448 /NCGR_PEP_ID=Amphidinium-carterae-CCMP1314-20130924|23864_1
/TAXON_ID=2961 /ORGANISM="Amphidinium-carterae-CCMP1314" /LENGTH=567
/DNA_ID=CAMNT_0029246317 /DNA_START=96 /DNA_END=1799 /DNA_ORIENTATION=-
MGCSQSVPAAKPAGALEEKALGVIRLDYDYSPAKGDIDHPGSFCYPVYYRVVPGLTFSMCQTGQMTPEVETKFSEAI
KYLESRNVCAITGDCGFMMYFQEKARQLTKKPVFMSSLAHLPSVTSSFASEEKIAIFTANSSSLAPMKPLIKSECGVEV
DDTRYVIVGCEDVAGFEAVEKGEKVDLAKVTPGIVKKCRTTLAKDPSIKAICFECTQLPPFSDALRHVTKLPVFDAITCC
DFFMSGLRDNPRFGVDDWHKQWDGLQEEYKFGKNLSAEDKAALQNKAAEDEGKTMSQLAEPLVKKQKQAMHLFSK
VRADRCNAKVGILRMDGERTAQPGDIISPQSFAYGAVYKAVPGLTREMCESGEYLADDVKSKVLEAIGDLVDQNGAK
SIVGDSAAMVHLQSLVRQRCSMPVSMSPLTQLPALLCAYNKTEKIAILTDASGDCAKAMASLKEADGSALDDSRLVIVS
CTDVGEVGLVSKAQQAMKDASIRAFLLENPELPRHSDAIRAATGLPVYDAITGSDMLMSAFLDNPRFSNSTAWHLNW
DGVQETLDWMKYLSPEDKQLFGK

>A.carterae_E CAMPEP_0186432918 /NCGR_PEP_ID=Amphidinium-carterae-CCMP1314-20130924|16687_1
/TAXON_ID=2961 /ORGANISM="Amphidinium-carterae-CCMP1314" /LENGTH=625
/DNA_ID=CAMNT_0029234477 /DNA_START=1 /DNA_END=1878 /DNA_ORIENTATION=-
WGMCTCFSRSALQEHQTARGERPEPLVPAAVQDGLDASRNDGDASPTSEPASPLSPVSPGFGLPKGSSRSRILNKA
KEAKEKSMHMAIGVLCVDEGSWPEPVHIEESIDSGTKIMARCKGVTLAILQEETLSEATLGAIKAAYKMILKKAEFRRLV
GVTADVGYLWLKHSWRVRLAIGADVPVCLSPLIQFPMISACFKNFESTCLVTWGSTEDWESRKAEFMESCRLKTASP
DQLIVCGIPSDEERWKKYLDWEACETDLEAYGEELVSLVKARLEQHRREHEGGADRIVKAILVTARLSRYTELLRTKLG
LPCFDEGTMLRFFKNAQGVGHYDDVNVLARLSDRLRGKDVLHIFGKKEVNTQKIGLIRLEYEYPPALGDPDHPGTFGF
NIVSKIVDGLTFEAAQEGSMEERILENMANAIQWLQEQGVIGITGNCGFMMHYQCFARFVAGVPCFMSALLQAATLEA

AVLPHERVLILTANSSSLEPGKDKLLRQSGISINESETFLIRGLQDLPGFEAVANAEKVDTIKVMNGISERVKSLMESQE
ADGNPIRLILLECTELPHYSSKLREVTQLPVFDLVTCVNFFFEATREVDWNVQTFSPHNEGFWKKNFAKFDRSQGAA

>S.triacnidorum 1123948 2555 10991
MFACALDVCARAAKKTCLRLTLAHKPPPRSPTTEKHIQGKDPNVKHASLGVVRLDWAYHPLPGDVGSPDSFEYPVYY
RAVPGLTFEICQSGKMTPEIQQNFKDAIEFLDKERGVSVITSDCGFFMWFQKEARLYTSKPVVMSSLALLPSLHASLGT
DGKVAIFSANSSSLGPMHDTVAKECGVDWNEASYVLVGCQDVEGFDAVASGEPLDLEKVTPGIVRKAKQVIAEHPDI
HAILMECTQLPPFSDDVRAATGLPVYDAIVSADFFIRGFVDNPRFGLNDWHQAWDGQQEKYKLGDNVQDKGKLLYYK
P

>Symbiodinium_A1 (identical to S.microadriaticum_A)
MPLILGGLAGAAAGYGAAAWLERRKPPPGKDPNIKHASLGVVRLDWHYHPLPGDVGSPDSFEYPVYYRAVPGLTFEI
CQSGKMSPEIQQNFKDAIEFLDKERGVSVITSDCGFFMWFQKEARLYTSKPVVMSSLALLPSLHAAIGTDGKIAIFSAN
STSLGPMHDTVAKECGVDWDEACYVLVGCQDVEGFEAVASGEPLDLEKVTPGIVRKAKQVIAEHPDIHAILMECTQLP
PFSDDVRAATGLPVYDAIVSADFFIRGFVDNPRFGLNDWHRPWDGHQEKYKLGDNVQDKEKLLHYKP

>S.microadriaticum_A
MYGCHTNAQAAVPCPSTEKHIQGKDPNIKHASLGVVRLDWHYHPLPGDVGSPDSFEYPVYYRAVPGLTFEICQSGK
MSPEIQQNFKDAIEFLDKERGVSVITSDCGFFMWFQKEARLYTSKPVVMSSLALLPSLHAAIGTDGKIAIFSANSTSLGP
MHDTVAKECGVDWDEACYVLVGCQDVEGFEAVASGEPLDLEKVTPGIVRKAKQVIAEHPDIHAILMECTQLPPFSDD
VRAATGLPVYDAIVSADFFIRGFVDNPRFGLNDWHRPWDGHQEKYKLGDNVQDKEKLLHYKP

>S.microadriaticum_B
MLDLFASASALSHYNDAKVLCRLEGKLGTKRSPSPNPSSPTSSHENCKLGLLQLEYEYPPAFGDIDHPGTFGFKTCPR
VVKGLTFQKAQEGSFDQDILENMTSEIRRLEQEGVVGITGNCGFMMHYQCFARYVSPVPVFMSALIQAATMAAAMEP
AERVLILTANAESLQPGKDKLLLESGIQVTNSEKFVIRGCENLPGFEAVANAERVDTIRVQNAISAYVLEIIKEESAREGY
GAIKMILLECTELPHYADELRRITGLPVFDAVTCVNYFFYATAAQNWNTATFTPHNLSFWTSNFDEDGRVRTGGKCSS
DGCAAPGV

>S.microadriaticum_C  OLP88173.1 hypothetical protein AK812_SmicGene30557 [Symbiodinium microadriaticum]
MAKSCLNTGSIPDWIATKARILSEMMRNMLSDEPLLLIRLPAAPSTGPLMAACKAPFNFARYEDRMVQWASAATSALS
LWCTDLGEGLLPELVEWMGWSDTGLEVQTNRKMILTPPGEVFGVSKMMHGKPATKPSGPCATLWAEEDTLEGQHQ
SEINNVAQWRSDTNVIQEPQKKCTFRKRIFEAARRTREVSMKKVFGVLCVDDGLTQEDLVEDGADLQRCHVVCRGIT
RERLRSDHLDEEALASIRASLFRLCQGHRPLAGLAVDLGYLWMKHQGALRRLVPRWLLLTTPLVQLPLIRTCFRTRDA
TLLVVWEDAAVSELSWLEASSNIRIDSQVEVIVVRASQPEWSNFLSKQTDQQEQALLLDLVDRVRIRVAELSRKDVAV
SSILVDGALSRFSKGLRKATQLPIFDEVSMMGLFSAASSLSEFSEASVVSRLEDRLQRSGNSQNRERDRPTGQLGIVQ
LDSYEYVRAVGDADHESTFSFQTCPRMAEGLTFEAAQKAAQEPVMLESLRQLAKEMETAHCFGIAGNCGFMQFYQD
LVREAVSVPVFLSALVQVPTMAAALDPRDRILILTANETSFREAQELLLRAECCNVALDQIVVRGCENVPGFEAVAKME
QVDTQKVENSLSQFVEEILAESDGAFIKMILLECTELPHYAARLRRVSGLPDLDLVTCANFFAKVLVGSF

>Symbiodinium_sp.C15_A CAMPEP_0192409804 /NCGR_PEP_ID=Symbiodinium-sp-C15-20130923|6074_1
/TAXON_ID=226949 /ORGANISM="Symbiodinium-sp-C15" /LENGTH=320 /DNA_ID=CAMNT_0036574075
/DNA_START=1 /DNA_END=962 /DNA_ORIENTATION=+
XKLQRRRRGTFRISAMPMLMLGGLGVCAGYACALWMKEQEPPKGKDPNIKHASLGVIRLDWSYSPLPGDVGSSDSF
EYPVYYRAVEGLTFEVCQKGEMTDEIKARFKEAIRYLDQDKQVSVIISDCGFFMWFQKLARDYTNKPVVMSSLALLPAI
HAAVGKTGKRTGKIAIFTANRDSLKTIHETVAEEMPIDWTKKCYLLVDCRDVPGFEAVATGEPVDFDKVSPGIVKKAVE
LTKNHRIDAFLFECTQLPPFSDDVRAATGLPVYDAVVCADFFVRGFVDNPRFGLNDWQQSWDGKQEAYQLGDVEDK
EKLVHYNQTH

>Symbiodinium_sp.C15_B CAMPEP_0192460220 /NCGR_PEP_ID=Symbiodinium-sp-C15-20130923|119450_1
/TAXON_ID=226949 /ORGANISM="Symbiodinium-sp-C15" /LENGTH=683 /DNA_ID=CAMNT_0036646103
/DNA_START=1 /DNA_END=2050 /DNA_ORIENTATION=+
XLDQRLQLMEMELQNLRQRNGGAMQMASANPMGLNHFAPQFMGPPSLQMTPRQPEETVDPQEVDRRRKAHQEM
LALHHDLLQSHTSLQKQHADLMKAHKELLHAFKNASFNGAAASAAVAKKQVKHPALGVVRLDYDYPPAPGDSDHPAS

FGYDVYFRCVPGLSFEMCQTGQFTEEVERRFADAIKHLEARGVSAITGDCGFMMAFQVLARKIAAKPIFMSSMVQCPI
IAAALDPDDQIMVLTANEHSLKPQKEILMTSCGFDVEEHRFVIHGCQDVPGFDAVAKGEKVPLDIVQPGIIKMALDLIQK
NPMIRAILLECTELPPYADALRYHTGLPVWDAITAADFYVSAFQDNPRFGISDWQQEWDEEQEHYELGTHLTKDELLL
LQNKAVHKKEKKQRALLHKKKLDKVQRKVRKQQAPILGVIRLDYNYPPAPGDIDFPGSYDYDVLFRVVPGFTFAMAQS
GKLSEAVEQEFIDAVRWLERKGVSGITGDCGFMMAFQPLASAIASVPVFMSAMMQCPMISIAFDKYDKILILTANGDSL
KPQKETLLKQCGFNVDDRRFMIVGCQDVPGFDAVAKGEKVDVDYVTPGIVYMVKEILKQHASIRAICLECTELPPYAD
ALRSETKLPVFDAITNADFFISARRDNPRFGFNQWQLDWDGLQDNYEFGSNLTAEQAEKLLN

>Symbiodinium_sp.C15_C CAMPEP_0192423434 /NCGR_PEP_ID=Symbiodinium-sp-C15-20130923|18175_1
/TAXON_ID=226949 /ORGANISM="Symbiodinium-sp-C15" /LENGTH=556 /DNA_ID=CAMNT_0036593715
/DNA_START=39 /DNA_END=1703 /DNA_ORIENTATION=+
MGCTTSSAATNLASLPQSLVETFGSYDVFVKENKAKLPVLGMLMLDAKSGKRDMGGFDHPGSFSYQIKQQVVKGLTL
EVCHSGKLTPDLEKSFVASISALEAAGASVICGDTGFMMWFQSLARQSTKLPVILSSLAVLPSITCSYGAHEQIAVLTAD
SQLLKPMATLIHDECSVDINDQRFVFVGCEDIPGFQQISAGQHDDAAKMEAGLVAKATKVLEEHPQVRAFLLESTELP
RYAKALRKATFRPVFDALNLCDFYIASHLDNPRFGVKDWDDAKQGKRPEFNFGQLVSNATEIVKPVLDTLKVTDIAMQI
YETSKQAAVSMLVRSKRAGITLGVIRLDYHYPPAPGDVDHPGSYPYEVHYHMVKGLTFEMCQAGKLTPEVETAFLQG
IKDLEAKGVSVITGDCGFMMWLQELARRNTKLPVIMSSLATLPAITASYTRNEQVAIFTANGQSLLGMEQLIKAECGVD
VKDKRYIFVGCEEVPGFEAVALGQKVDVRKVTPGIVKKAQEILTKHPQVRAILMECTELPPYSDAVRHVCGVPVFDXA
LGQKVDVX

>Symbiodinium_sp.C15_D CAMPEP_0192423268 /NCGR_PEP_ID=Symbiodinium-sp-C15-20130923|18047_1
/TAXON_ID=226949 /ORGANISM="Symbiodinium-sp-C15" /LENGTH=647 /DNA_ID=CAMNT_0036593495
/DNA_START=1 /DNA_END=1942 /DNA_ORIENTATION=-
XFDIALGRSGLAGADKGENSGXEMVQMTMVTEVQKMKDMTDMKDMKASKLEAVDNLKGVDSVDVISTEVQTACGN
QQQYSNEAFVAQPVITTNAEDDTSAPSPQPLAPVAPVAPTLSQRLFKAAKKTKAAQTKECFAVLCLDKGDGDEEVIEV
KEAGCKRLLSVCKGVTKEKLMSDRLDSSAVASIKACVDRIIGKAGKESIAGISCDVGYLWLKHQRTLRQALPQFPVLTT
PMMQLPFIWTCFGNEKTLLVTWKDEVLCEDSESHHLDFLDTAGIRLDTDRVEILTLSTSDLEWSAFLSGMTTHTTASE
HERLLNQLVNMVHQKVIQFSDKDGVGHGVNSVVLDSMLSPFSKELSAATDLPVFDEVSMLRLFSSASSLSHFSDANV
LCRLDESQSQSQRSKIDTIDGTRMGLVRLEHEYPYGVGDIDHGSTFPFQSCPGVVQGLTFEQAQRGSKDPVILENLK
RVVDQMEAKECFGIAGNCGFMHFYQEFVKNYATVPVFMSALVQVPTMAAALEPDERILILTANESSFMESRDALLSAE
GRPFCDFNRVLVRGCEHVPGFEAVANATLVDNIKVQENMSKYVEEILEQEKDSAKGPIKSILLECTQMPHYAAAIRQST
GLPVFDVVTCVNFFASSLCPRHERRG

>Symbiodinium_sp.Mp_A CAMPEP_0192603402 /NCGR_PEP_ID=Symbiodinium-sp-Mp-20130822|28546_1
/TAXON_ID=230996 /ORGANISM="Symbiodinium-sp-Mp" /LENGTH=301 /DNA_ID=CAMNT_0036815439
/DNA_START=65 /DNA_END=970 /DNA_ORIENTATION=-
MPLIIGGLCGAALGFGAAAWVENHKPPKGKDPNIKHASLGVVRLDWSYHPVAGDVGSSDSFEYPVVYRAVPGLTFEV
CQSGKMSPEIKQSFKEAIEYLDKERGVSVITSDCGFFMWFQKEARLYTAKPVVMSSLALLPALHAALGTDGKIAIFSAN
SVSLAPMHDTVAAECGVDWNEGCYVLVGCQDVEGFDAVAHGDPVDLAKVTPGIVRKAKAVLAEHPDIHAILMECTQL
PPFSDDVRAATGLPVYDAIVSADFFIRGFVDNPRFGLNDWHQAWDGQQEEYKFGDHVKDKSNLVHYKG

>Symbiodinium_sp.Mp_B CAMPEP_0192627018 /NCGR_PEP_ID=Symbiodinium-sp-Mp-20130822|90026_1
/TAXON_ID=230996 /ORGANISM="Symbiodinium-sp-Mp" /LENGTH=708 /DNA_ID=CAMNT_0036843329
/DNA_START=61 /DNA_END=2187 /DNA_ORIENTATION=-
MMPGFSPIDSRLDMLEMELQRLKGVRPPPFGATGLPHMTFTQPQQALAPMVPPGSAFAPNYAMPHEPTGPAPALAN
VEGKTTSEIMTAYKDIALALKDAQAKHNDLLLRHTELSKHHADLMKAHHDLLEQLKKGGIAKAGAGTNGGHRAMHPSV
GVVRLDYDYPPAPGDSDHPGSFGYDVFFRCCPGLTFEMCQAGKFSEEVERRFADAIKHLEARGVSAITGDCGFMMA
FQVLARKIASKPVFMSSMVQCPIIAASLEPADQILILTANDLSLKPQKQVLMKSCGFDVDEDRFIIKGCQHVPGFEAVAL
GQKVPLEKVQPGIVQLTLDSMKEHPRISAILLECTELPPYADALRYHTGLPVWDAITCADFYIAACKDNPRFGQNDWQ
LEWDGEQEEYELGKNLTKDELDCVVCKKKELAEKKARALHKKKKLDKIQRKVRKQQAPMLGVLRLDYNYPPAAGDID
CPGSYDYDVLFRVVPGFTFEMAQSGKLSPDVEKEFIGAVKWLENKGCGGITGDCGFMMAFQPLARKIASVPVFMSA
MMQCPMISIAFDKYDKVLILTANDATLKPQKETLLSQCGFDVDDDRFLIRGCQNVPGFDAVAKGEKVDVEYVTPGIVY
MVKQLLKKMPTIRAICLECTELPPYADALRAETKLPVFDAITNADFFLSARQDNPRFGFNQWQLDWDGEQDEYEFGQ
NLTAMQQAKLIN

>Symbiodinium_sp.Mp_C CAMPEP_0192576804 /NCGR_PEP_ID=Symbiodinium-sp-Mp-20130822|7847_1 /TAXON_ID=230996 /ORGANISM="Symbiodinium-sp-Mp" /LENGTH=564 /DNA_ID=CAMNT_0036782987 /DNA_START=54 /DNA_END=1748 /DNA_ORIENTATION=-
MGCASSTASSLAALPQSLAQSFASYEIFVKESKAKLPVLGMLMLDGKGQQDVGSFDHPGSFSYLVKQRVVKGLTMEI
CQSGSLTEDLEKAFVESIAALEAAGASVICGDTGFMMWFQPLARRSTNLPVILSSLAVLPSVTCSYGAHEKIAVLTANS
ESLKPMASLIHDECGVDVSDSRFVFVGCEDVPGFEAIAAGQHGDFAQMEAGLVKKAIAVLEEHPQVRAFLLESTELPR
YAKALRKATFRPIFDAVNLCDFYISSHLDNPRFGVADWDEAKQGKKPQYNFGQLMSDTAGIVKPVLDTLKVTDVAMQI
YESSKQVAVSMLARSRRAGVKLGVIRLDYHYPPAPGDVDHAGSYAYEVHYHMVKGLTFEMCQAGKLTPEVETAFLQ
GIKDLEAKGVSVITGDCGFMMWLQEKARRNTKLPVIMSSLAILPAMTASYHKDEQIAIFTANGKSLLGMEQLIKEECSV
DVKDQRFIFVGCEEVPGFEAVALGQKVDVKKVTPGIVQKAQKILTQHPKVRAILMECTELPPYSDAVRHVCGIPVFDAI
TACDFYIASHVEHPMMKA

>Symbiodinium_sp.Mp_D CAMPEP_0192623584 /NCGR_PEP_ID=Symbiodinium-sp-Mp-20130822|75336_1 /TAXON_ID=230996 /ORGANISM="Symbiodinium-sp-Mp" /LENGTH=628 /DNA_ID=CAMNT_0036838139 /DNA_START=1 /DNA_END=1886 /DNA_ORIENTATION=-
XFGSRFGPNLPLSMITSPTSSAKCRSMLLSLAKVAEKKRSEKTFAVLCVDEGNWPHEVMRVDTGAGIKVMSKCKNVT
RACLMTDTLDKDSLKSIRECFERLSSVAEEITGVTADLGNLWLKHHSAVRSVIPERPVYLTPIIQLPMISASFKSSEQTLL
VTWGSQDHVDSMKPKFVSDCNIVLDMDRLETLGVDPLQDSRWARYLNFDTDAAEDEALGSVLVCMVLEKIAAINVEG
KKDKLIKSIIVTTRLSMFSGKLREETGLPVFNEITMLNLFSSASSLSHFHDAKVLLRLDEKLAKAPSPRSPTSPACAAGP
KPRLGLLQLEYEYPPAFGDVDHPGTFGFQMCPRTVKGLTFERAQEGSFDPSILEHMAKEIQALEEEGVVGITGNCGF
MMHYQCFARYVAHVPVFMSALIQAATMAAATEPLERVLILTANASTLQPGKDKLLLESGIQVTASDKFVIRGCESLPGF
EAVANAERVDTIRVQAAISDYVMRTLKEENEKDGAGAIKMILLECTELPHYADELRRTTGLPVFDAVTCVNYFFHATAA
SNWNTGTFTPHNPSFWSSNFDADGRKLKDAGAPARSESDESSAKLGEVSAVFSKSQLEIVQSIIGVHAYLHSPEWLQ
L

>Symbiodinium_sp.Mp_E CAMPEP_0192649158 /NCGR_PEP_ID=Symbiodinium-sp-Mp-20130822|189624_1 /TAXON_ID=230996 /ORGANISM="Symbiodinium-sp-Mp" /LENGTH=730 /DNA_ID=CAMNT_0036871389 /DNA_START=1 /DNA_END=2192 /DNA_ORIENTATION=+
XADLRNMERKENVLPHSPWFYKGAQAQAEQQLPQNPQHIRRPARDEPQKLWLTAEQFGEEADASTEAGTEGSTTS
EEEDDEEPEKKVLPKAASVVLSNPKVCPALPEAQPRSSERGLLSQRLFTAARRTQQALTRETFAVLCLDEAIAVHSQE
VEEVSEDGVTRLLSVCHGVTRAHLMPDQMSTKAIESIRACLDRITCRLAGGLLAGVTVDVGYLWLKHQRTLRQLFPSL
PVIITPLLQLPLILTCFGTTAKTLLVTWEDADAPDPHLSILEAAGMRLDQTRVETLTICATSAKWPDFLFRTTTAAEEREL
LLEVVNMVQQRLMTLLDEDVRINSIVLDHGLAKFGSYLSDMVRMPVFDEASALSLFKSASSLSHFSDANVLCRLEAML
TQREPEGHGLRHTQETNGLGIVQLEYEYVAAVGDVDHHSTFSFPTCQQTVEGLTFDKAQCGVQEASILDSLMSMVEL
MESEGCFGITGNCGFMHHYQWLARDAASVPVFLSALVQLPVMAMALEPEDRILVLTANESSFMKFRDELLSADGIQA
RVFDRVVVRGCEDVPGFEAVANAEQVDTIKVQENLSKYVEEIMAEDAGQIKMILLECTELPHYAAEIRRKSQLPVFDIV
TCANFFAKVLLRSNHGRAPEFDACAFTVSSPLAPWQSNSHSNSHQLPDELSSCIKDMREIDMLKTEKTSAASASVGE
TIRARTSTVTAVLPSWQPASTTLAFRESAG

>Symbiodinium_sp.C1_A CAMPEP_0199584958 /NCGR_PEP_ID=Symbiodinium-sp-C1-20140214|19746_1 /ASSEMBLY_ACC=CAM_ASM_001145 /TAXON_ID=226962 /ORGANISM="Symbiodinium sp., Strain C1" /LENGTH=301 /DNA_ID=CAMNT_0045427509 /DNA_START=65 /DNA_END=970 /DNA_ORIENTATION=-
MPVLMLGGLGVCAGYAFAVWVKGQEPPKGKDPSIKHASLGVIRLDWHYHPLAGDVGSTDSFEYPVFYRAVPGLTFE
VCQSGVMTPEIKEHFKEAIRYLDQDKQVSVITSDCGFFMWFQKEARQYTAKPVVMSSLALLPAIHAALGTDGKIAIFSA
NSESLQPMHDTVAREMGVDWNADCYVLVGCQDVEGFEAVATGEPVDLGKVTPGIVKKAVEVTKTKNIHAILMECTQL
PPFSDDVRAATGLPVYDAIVCADFFVRGFVDNPRFGLNGWHQRWDGKQETYKLGDVEEKEKLVYYSPTH

>Symbiodinium_sp.C1_B CAMPEP_0199597350 /NCGR_PEP_ID=Symbiodinium-sp-C1-20140214|27762_1 /ASSEMBLY_ACC=CAM_ASM_001145 /TAXON_ID=226962 /ORGANISM="Symbiodinium sp., Strain C1" /LENGTH=685 /DNA_ID=CAMNT_0045441819 /DNA_START=1 /DNA_END=2056 /DNA_ORIENTATION=-
XGLDQRLQLMEMELQNLRQRNGGAMQMASAPMGFNHFAPQFMGPPSVQMTPRQEENVDPQEVDRRKKAHQEML
ALHHDLLQSHTSLQKQHADLMKAHKELLQAFKNASFNAAGGAAGAAAAAKKMVKHPALGVVRLDYDYPPAPGDSDH
PASFGYDVYFRCVPGLSFEMCQTGQFTEEVERRFADAIKHLEARGVSAITGDCGFMMAFQVLARKIAAKPIFMSSMV

QCPIIAAALDPDDQIMVLTANEHSLKPQKEILMTSCGFDVEEHRFVIHGCQDVPGFEAVAKGEKVPLDIVQPGIIKMAM
DLIQQNPMIRAILLECTELPPYADALRYHTGLPVWDAITAADFYVSAFQDNPRFGISDWQQEWDGEQEHYELGTHLTK
DEMLLLQNKATHKKEKKQAALLHKKKLDKVQRKVRKQQAPILGVIRLDYNYPPAPGDIDFPGSYDYDVLFRVVPGFTF
AMAQSGKLSEAVEQEFIDAVRWLERKGVSGITGDCGFMMAFQPLASAIASVPVFMSAMMQCPMISIAFDKYDKILILTA
NGDSLKPQKETLLKQCGFNVDDRRFMIVGCQDVPGFDAVAKGEKVDVEYVTPGIVYMVKQILKQHASIRAICLECTEL
PPYADALRSETKLPVFDAITNADFFISARRDNPRFGFNQWQLDWDGLQDNYEFGSNLTAEQAEKLLN

>Symbiodinium_sp.C1_C CAMPEP_0199555188 /NCGR_PEP_ID=Symbiodinium-sp-C1-20140214|794_1
/ASSEMBLY_ACC=CAM_ASM_001145 /TAXON_ID=226962 /ORGANISM="Symbiodinium sp., Strain C1"
/LENGTH=678 /DNA_ID=CAMNT_0045394265 /DNA_START=1 /DNA_END=2032 /DNA_ORIENTATION=+
XPRWVPPLSPLPSPLPPMGPMGLTGMQGMTPRLLPMPPQHLNKPSFVHPPPQDGFEMEEEDFKELQTLHQELVDS
HNELRNEHAHLLKAHEDLIIAMRQNATNGAEKNLKRHPALGVVRLDYEYPPAPGDSDHPGSFGYDVYYRCVPGLTFE
MCQSGQFTELVERRFADAIKHLEARGASAITGDCDFMMAFQVLARKIASKPIFMSSMVQCPIIAASLDPDDDILILTANS
ESLKPQKEILLTSCGFDVSKERFHIHGCEDVPGFEAVSLGQKVDVEKVQPGILQLVKGIIKEKPEIRAILLECTELPPYAD
ALRYHTGLPVWDAITAADFYVTAFRDEPRFGQQDWQREWDGQQEAYELGDHLTKDEQALLVNKAQKNLGKKVEAL
SKGQQIEQIQKKVRRQQAPILGVIRLDYNYPPAPGDVDFPGTYNYDVLFRVVPGFTFAMAQSGQLSEDVEQEFIDAVK
WLEMRGVAGITGDCGFMMAFQPLASSIASVPVFMSSMMQSPMISVAFDKYDKVLILTANDESLKPQKETLLRQCGFN
VDDQCFMIVGCQNVPGFNAVAEGRKVDVEYVTPGIVYLVKQHLKKHPSVRGILLECTELPPYADALRDETGLPVFDAI
TNADFFISARMDNPRFGFNQWQLGWDGIQDTYEFGSNLSFXKGVETLELTVELTANG

>Symbiodinium_sp.C1_D CAMPEP_0199592752 /NCGR_PEP_ID=Symbiodinium-sp-C1-20140214|25073_1
/ASSEMBLY_ACC=CAM_ASM_001145 /TAXON_ID=226962 /ORGANISM="Symbiodinium sp., Strain C1"
/LENGTH=584 /DNA_ID=CAMNT_0045436773 /DNA_START=1 /DNA_END=1753 /DNA_ORIENTATION=+
XKRSTPSETDAMGCTTSSAATNLASLPQGLAETFASYEIFVKENKAKLPVLGMLMLDQKSGKREMGGFDHPGSFSYQ
IKQQVVKGLTLEVCHAGKLTPDLEKSFVASISALEAAGASVICGDTGFMMWFQSLARKSTKLPVILSSLAVLPSITCSYG
AHEKIAVLTADSQLLKPMATLIHDECSVDINDQRFVFVGCEDIPGFHQISSGQHDDAAKMEAGLVAKATKVLEEHPQV
RAFLLESTELPRYAKALRKATFRPVFDALNLCDFYITSHLDNPRFGVKDWDDAKQGKRPEFNFGQLVSGATEIVKPVL
DTLKVTDIAMQIYETSKQAAVSMMVRSQRAGITLGVIRLDYHYPPAPGDVDHPGSYPYEVHYHMVKGLTFEMCQAGK
LTPEVETAFLQGIKDLEAKGVSVITGDCGFMMWLQELARRNTKLPVIMSSLATLPAITASYTRDEQVAIFTANGQSLLG
MEQLIKAECGVDVKDKRYIFVGCEEVPGFEAVALGQKVDVRKVTPGIVKKAQEILTKHPQVRAILMECTELPPYSDAVR
HVCGVPVFDAITACDFYIASHVEHPMMKTWSHAPLG

>Symbiodinium_sp.C1_E CAMPEP_0199605684 /NCGR_PEP_ID=Symbiodinium-sp-C1-20140214|37686_1
/ASSEMBLY_ACC=CAM_ASM_001145 /TAXON_ID=226962 /ORGANISM="Symbiodinium sp., Strain C1"
/LENGTH=227 /DNA_ID=CAMNT_0045451253 /DNA_START=1 /DNA_END=683 /DNA_ORIENTATION=-
XQLDTHFPRIPGDIACRDTYHRPVGVQIIDRAQVASVVNADPNGLDITGFTSALASLENIGADITSKDQIISTSCGFMIYF
QESFSKMTDKTFISSSLIALPELRSRFPDPEIMVITFDADVLASPAYQPALDGFAGPVIGLEKWMHLYEVISQDLVDLDF
HKAELGMDQLITAALRRFPVKAIILECTNLPPYKHVIRRHFAGEIIDCLSVLEAASPGLVKEQHLM

>Symbiodinium_sp.C1_F CAMPEP_0199596160 /NCGR_PEP_ID=Symbiodinium-sp-C1-20140214|27039_1
/ASSEMBLY_ACC=CAM_ASM_001145 /TAXON_ID=226962 /ORGANISM="Symbiodinium sp., Strain C1"
/LENGTH=587 /DNA_ID=CAMNT_0045440493 /DNA_START=1 /DNA_END=1762 /DNA_ORIENTATION=-
XHMAAYSSSPTSTSPTSTSPTSPQSEARARGRLLQLAKVAEEKRSQQTFAVLCVDEGTWPQDVMKVDAGCALKLMS
RCKGVTRACLMADLDAEALRSIRECFERLLAVSESIIGVTADLGNLWLKHHSAIREVISERPVYLTPIVQLPMISASFKSR
EQTLLVTWGSQKEVEMLKPTFCSGCNIVMDLDRLEILGVDVSQDARWARYLNFDTNTDEDMALGSGLLQMVKDRISE
VNVQGKKDMLIKSLIVTTRLSMFSNLLREGTGLPVFNELTMLNLFSSASSLSHFNDAKVLCRLSDKLARSPAKSPGVSP
GGCKLGLLQLEYEYPPAFGDVDHPGTFGFKTCPRIVKGLTFEKAQEGSFDQNILEEMSCEIKKLEEEGVVGITGNCGF
MMYYQCFARHIAKVPVFMSALIQAATMAAAMEPAERVLILTANAATLQPG

>Symbiodinium_sp.C1_G CAMPEP_0199588256 /NCGR_PEP_ID=Symbiodinium-sp-C1-20140214|22040_1
/ASSEMBLY_ACC=CAM_ASM_001145 /TAXON_ID=226962 /ORGANISM="Symbiodinium sp., Strain C1"
/LENGTH=668 /DNA_ID=CAMNT_0045431505 /DNA_START=1 /DNA_END=2004 /DNA_ORIENTATION=+
XAKWPGADEVDNSGXEMVQMQTMVTVQVQDMNDMKASKLAAVDNLKGVDVISTKEVHTACGNEQKQYSNEAFVA
QPVITANAEDAASAPRVAPVAPVAPTLSQRLFKAAKKSKAAQTKETFAVLCFDRGDGDEEVVEVKEGGCKRLLSVCK

GVTKEKLMSDRLDCSAVASIKACVDRILGKAGKESIAGISCDVGYLWLKHQRTLRQALPKFPVLTTPMMQLPFIWTCF
GNEKTLLVTWKDEALCEALCETDSESSESHHLGHLGFLETAGIRLDTDRVEILTLSTSDLEWSPFLSGMTTPSENEKLL
NQLVNMVHQKVIQLLDKDGVKGVKGVGVNSVVLDSMLSPFGKELSAATNLPVFDEVSMLKLFSSASSLSHFSDASVL
CRLDEKSQSQRSKIDGTRMGLVRLEHEYPYGVGDIDHGSTFRFQSCPGVVQGLTFEEAQRGSKDPVILENLKRVVDK
MEAEECFGIAGNCGFMHFYQEFVRDYATVPVFMSALVQVPTMAAALEPDERILILTANESSFMESRDALLSAEGRPFC
DFNRVLVRGCEHVPGFEAVANADLVDNIKVQENLGEYVKEILEQEKESDKGPIKSILLECTQMPHYAAAIRQSTGLPVF
DVVTCVNFFASSLCHPLPKAQVLNGVNGDAKKCDHDRQCPTDVEHQRKG

>C.goreaui clago109
MGGVMTTARGGQALYGAHLGILMLEARFPRIPGDMGNATTWPFPVHYRVVRGASPDRVVRQRAEGLTDAFVAAAR
ELVADGVDGVTTNCGFLSLIQGELAAACQVPVATSALMQVPIIQSLLPPGRRVGILTVSAEDLTAAHLQAAGVPADTPV
VGTEQGREFSRVILNDELEIDVAAASDDLLQAGRRLVSDHADVGALLLECTNMAPYARLLRAELGLPVFDIYSFITWFH
AGLRPHAFGPPVPGAAWRGV

>A.temarense_A CAMPEP_0186397762 /NCGR_PEP_ID=Alexandrium-temarense-CCMP1771-20130823|410328_1
/TAXON_ID=2926 /ORGANISM="Alexandrium-temarense-CCMP1771" /LENGTH=865
/DNA_ID=CAMNT_0029195673 /DNA_START=1 /DNA_END=2592 /DNA_ORIENTATION=+
XAQPSRPPHRAEMKSMAKSHSMTSLSNPFPMMSNYGPGPMQGSDRLDELEMELNQLKAMRQQELYYGQERLAGIA
RPGGGPMPYAQRPASASMQLPPLSPQPYMQAPFQHQQQQQQQQQQQQQQQQQQQQQQQQQQQQQQQFVP
MPYAMPPMQPPPPAGVSSQAHHDLLTAHKQMLVAMKEQEERHSQLLRTHTELMRAHTDLIKKATAAGASKEKKLKK
NPSLGVVRLDYKYPPAEGDIDCPASYGYDVFYRVCPGLTFDMAQDGHFTEAVEREFAEAIKYLEMRGVSAITGDCGF
MMAFQVLARKIASKPIFMSSXAIKYLEMRGVSAITGDCGFMMAFQVLARKIASKPIFMSSMCQCPVISAAFEPNDQILIL
TANGATLKPQKQVLLNSCGFDVNEERFVIKGCQDIEGFDAVAKGEMVPLEIVQPGIVKLTLEILKTRPSIRAILLECTELP
PYADALRAATGLPVWDAITGADFYIRAFRDAANFGFSEWQEEWDKQQDSYAFGQNLVESDKKKLVNFADDPSQPEV
KRSTTSRKALAVSIRTQPEAVEVAKRTQRKLVSKQAPTLGVVRLDYNYPPAPGDIDCPASYSYDVLFRCVPGLTFEMA
QAGKMTATVHEEFVKAVRWLENKGVCGITGDCGFMMAFQPIASKAATVPVFMSSMVQSPMISVAFDKYDKILILTANS
TTLKPQKEILLSHCGFDVDDSRFVIYGCQDVPGFDAVSKGDKVDVEYVTPGIVKMTKDILLKEPTIRAILLECTELPPYA
DALRAASGLPVFDAITCTDFFVSAYKDNPRFGLNKWQCPWDGENEEYELGQNLSADQAGNLIKAEGNLPAARPAVG
QASGLELTLIHGDSAPQA

>A.temarense_B CAMPEP_0186178466 /NCGR_PEP_ID=Alexandrium-temarense-CCMP1771-20130823|2738_1
/TAXON_ID=2926 /ORGANISM="Alexandrium-temarense-CCMP1771" /LENGTH=764
/DNA_ID=CAMNT_0028940589 /DNA_START=1 /DNA_END=2293 /DNA_ORIENTATION=+
XRLEQLEQDIGGGANLRAVGAQIMMSGPGMQGYGGYASMPNRITAADRLEQLEQELYHLKSQRQEEHQYPPGPVLP
DSGVMGPMRLPPLSHNVSVQSGGQLMQQVPNFIPVPYPVHQPAAPATAASAGPSEQEHQELLVAHRELLDTMQKQ
QTRHTELLKSHSELMKAHMDLLKQTMNPGGGAAKKEAKKFPALGVIRLDYNYPPAGGDTDCPASYGYDVFYRAVPG
MTFEMAQAGTFTEAVERRFAEGIKYLEQRGVSAITGDCGFMMAFQVLARKIASKPIFMSSMVQCPVIAAAFEPHDQILI
LTANGNSLRPQKEVLLNSCGFDVNEDRFVIQGCQAIPGFDAVAKGEKVPLDIVQPGIVKMTMDILKTKKAIRAILLECTE
LPPYADALRASTGLPVFDAVTGADFYVNAFKDNERFGVSDWQEHWDQAQEEYEFGQNLTEKDKALLENYHPTKKEE
HRYKKKKQDMAKKKDDAKYLLEKTKKNLTRKAAPILGVVRLDYNYPPAAGDIDCPASYDYEVLFRCVPGLTFDMAQA
GRMTHTVQQEFVAAIKWLEAKGVAGITGDCGFMMAFQPIASEIASVPVFMSSMVQCPMVSVAFDKYDKIIILTANSKTL
KPQKETLLSRCGFDVDDARFIIYGCQDVPGFDAVDKGEKVDVEYVTPGMVKMTKDILKKDPTIRAIVLECTELPPYADA
LRKSTGLPVFDAITCADFFISARKDNPRFGLNQWQNDWDGVVNEYELGGNLSADXEGQAAECMSFPA

>A.temarense_C CAMPEP_0186226756 /NCGR_PEP_ID=Alexandrium-temarense-CCMP1771-20130823|37484_1
/TAXON_ID=2926 /ORGANISM="Alexandrium-temarense-CCMP1771" /LENGTH=781
/DNA_ID=CAMNT_0029000513 /DNA_START=1 /DNA_END=2339 /DNA_ORIENTATION=+
XYGSNFSHVGAQILMQSGSAPGGPPPAWAAMQQQQQQQPPKLNIHDRLDSLEHEIARLRTPRPAQHTDFHSMDPP
MSSRQHPAMAAAAQHALMHHGAMAVAAQQALLPQQVPSSYGVPPLPLANIGPCSTGPASSARTEEHQQLLSVHKQL
LQTMDRQETRHTQLLNSHRELMKAHKDILAMQAKGKGGPTKKKHPGLGVIRLDYNYPPAAGDTDSQASFNYDVFYR
VVPGMTFEMAQRGKFTEEVERNFAEGIKFLEMRGASAITGDCGFMMAFQVKARQIATKPVFMSSMVQCPVIACAYDK
KDQILILTANGHALRPQKEVLLSSCGFDVNEDRFIIKGCQDVPGFDAVAKGEAVPVEVVQPGIVKLALGMMKQHPRIKG
ILLECTELPAYADALRAKTGLPVWDAVTACDFYVSAFKDNPRFGVQDWQSEWDEEQDDYTFGMNLIEADRKELVHKV
GGTKQKKKADPRVRKVAEKTKQKLKRQKAPILGVIRLDYNYPPAEGDIDCPGSYDYDILFRCVPGLTFDMAQSGKMSF

TVQQEFVAAVNYLEKKGACGITGDCGFMMAFQPLAREIANVPVFLSSMVQCPMISVAFDKYDKIMILTANSATLKPQK
DVLLNQCGFDVDDTRFVIYGCQDIPGFDAVEKGEKVDVEYVTPGMVHMVQEILKKQPTIRAICLECTELPPYSDALRKX
DWPPGLRRDHVRGLLRVRQEGQSEVRHEPVAERLGRGGGGVRARAELGGGPAQPPDQRLGGLALPLPDSRGGW
ACWSFEGFARAPT

>A.temarense_D CAMPEP_0186321532 /NCGR_PEP_ID=Alexandrium-temarense-CCMP1771-20130823|165856_1
/TAXON_ID=2926 /ORGANISM="Alexandrium-temarense-CCMP1771" /LENGTH=678
/DNA_ID=CAMNT_0029103759 /DNA_START=82 /DNA_END=2114 /DNA_ORIENTATION=-
MAQSPASAEAKLPPLEELVVQIRKASPGILKQRHASLFCRVSNAATARELVSWMTDQPWCKGKEHACNVSQALLDTG
LMWNAAGSCSIFGTEGASLDMIFQVTQTFTIRDMDVMIDRLECIAKGIKVGDRKFRGQTYPQCFFATEILEYLTAQGIA
GSKAEAIEACHQLCAKHVMHRVDKDPLRRKKGQPFGFDAGRYPYRLSRDEMKLLGFDVRGISHSFVLSTADRIHADL
AKNNQGFFGRECVEWLSREGCGRTEKECTILAEYFVAAWIMRHIDEDQTCKFSAEKGCAFLRDLVTAKVEKKVILPQ
QPLKPKDRPLGVIRLDYGYPPIPGDIDHPHSFDFPVVYRKVPGLLFEVAQAGKLTPDVETGMRKAVQDLEAMGAWGL
TGDCGFMANYQGFVRGIAAVPVFMSSLSLLPLISQSLGPKDTILITTANSNSLKVLFPHVWQGQEPTRDDYQVSNKFL
QEQCGVSLARPEQILVRGFQDIPGFDVVAEATSLLSRKMDPVLISEGIVKDVKSILHARPDIKFILQECTELPHFTVALRQ
ATGLPVMDALSAPETFYHTIKFIIMQSQGVGIAWDIIARSPSRTRRRRDAIQAERGTRPCTRWQDGIMSLTCVWIFCVC
ASSWGTSFGMHVVLSPSGQGAALVAITGLVGSPSLNGDIMRIFDTRRLAAMCX

>P.aciculiferum CAMPEP_0190608078 /NCGR_PEP_ID=Peridinium-aciculiferum-PAER_2-20130926|2150_1
/TAXON_ID=268820 /ORGANISM="Peridinium-aciculiferum-PAER_2" /LENGTH=421
/DNA_ID=CAMNT_0034382289 /DNA_START=1 /DNA_END=1261 /DNA_ORIENTATION=-
PPYADALRAATGLAVWDAITAADFYISGYEDNPRFGVNDWQKEWEQAPAAPNAKGKAKAKSKADAQKVKQLAKKQA
PVLGVIRLDYNYPPAAGDVDCPGSYDYEVVYRCVPGLTFEIARTGMMTYQVQQQFTAAVKWLEMRGASGITGDCGF
MMAFQPLARDVASVPVFMSPMLQCPMVSVAFDYYDRIMVLTADGKNLKDQKEVFLAHCGFDVDDARFLIVGCQDVP
GYNAAAKGEKVNVEKVTPFICELVRGHLDKCPDIRSILLECTELPPYADSLRMATGLPVWDAITCADFFISARKDNPRF
GLNQWQNDWDGVVKEYTLGQNLTPDERSQAKXHQVTLGPCLFAMLPAVSSRGFRWSMTAALAAEVAASSAASAAA
ATAVAAEAGDRDVWPLEARFSHPSHALTRRLAMFRTAGX

>A.ceratii scaffold580
MQAPALCMPDKSTMSKSVRRQLGLPKQLKLRKPVLGVVRLDYNYPPAVGDIDSPLSYEYEVAYRVVPGLTFEMAQA
GTLTPEVGRQLCEAVRYLVEEVEVGAITGDCGFMFNFQPLVRQLHTSRQVPIGLSFRASPKKLSAGFFRHVLAYTLRR
SVKSAPPPYTSLDGMKHLIARECGLHLDEERFLIVGAESVPGFEAVAAGEKVDVERVTPGMVALVKDVAGILLECTQL
PPYAAPIRKATQLPVIDSVTTANFMMEGCLPPD
>Amoebophyra_ceratii2 jgi|Amoce1|13208|g945.t1
MLPTASNAPPKQRGADGEDTSSSSSSASEDAGDPTATAASGSGREEQEPPRATEVEQVREGAAAGGXGTTTADAH
RRGKKHASRRRRKLLAHSRTTGSIMLQDEVSTCRPGMACWFQGLSLQRYEAMFDRVARELRRNQEAATQEDLVTD
VIDEGLRSRAASQSHLTDDDIEVRQLDSSCDEVVEQIIPRPRLRTQVSKASSLPRAASRGALSGVSAALPSSPVLGGLP
ANAAPVTVPPQHCSLRYLNYDSAAAPQQAPPPFGHHGPLGLGEPPGPGPAPGPGPAFATAGGATTTSRRGSSATG
ASTARGADTFGTLLNPDDAEPTSWSVSVSGVYANGKSQQVMQHYNHSTRRESSTMSASAAAPLMRFPSLLQVPVL
GLGILRYDEEEDEPEPGTVAYSSEGVEEEVLELEVDVDSLESEDDDDEDGHYVEGSLASRSLTPAKTRTTGKRSRNT
RSTTTLQPRGAPSVDGAGERNDDEDEHVNEVKARLKAVHKSRRSRGNQHQDHQGPSNMSKSPEESLRDLDVDSIQ
QPSASVSSSSALATGKNGGTSTTGAPGVSSSAATFSTPQKGTSDEGGLLQENALTFSAWDSATSRRIELTSGTTSPT
TATGSTSTSRHHSTHGKTSRRRLRSRADTSLTSGYERVSVFRANASKRFQAGLVRRSTDDIALGLKSEKQTRERQTG
TRSGTALTMDGAAGAGVGPGGNSTLISDAPPAGGSESDRGGPESTSLAPEGPQAIYAATDYPLVLQKVPGLTPSLVY
ECAVLPQGQRKPFPEDAANCLYHSLRKLNDDPSVGFISGEDGLMASLQMVAHQVTRKPCCMSPLLMLPTYSRMYKP
NSKFIVCTVGVSAQDLYLNVDDLAERVGVSPTKFREHFVIERFEDILLEVEDVAKQGALIAWRIQSHFKVCSSAGVSSS
GTGSSSWGNTAQLSELVSALAEEFPHLVNKKNVTRGASTRKQEVSSYLGLPDNQNVVQVPSARPLTTVAGGAPPGA
LMKSQGQEQQQDGALLNLSAGINGGKHRLCGIFLECTEMSVHSDFLREMLAVPVYDMVALSHALMLGYLDNPLFGRI
GWQYHYNQNLYGDYVKSLVDEEGGSTGGSWGKREVESGRQTGGGGGGRGGGDTIMGPLRALKNSEADGAAAGE
AVVAEQVVPYVYAQSMLEKRLTVGGHGRTVDVRKFGTAAFSRGASAVILPAADAGGGEAPEAEMAQAAEGLDGGTS
SAVFNTASFNADHASAERELESYTTRTRSYPAPASAEGTSRSRETSKGVAGDCTDEDRASYQARRGQVQALIDFLLA
KKKAEEQAGGSPMNPPLPPRVGHHTTHSTQPALGCLRIDSLYRPXAAGDVADPRSHEYPLVNRVVPGLTFEVVRSG
KLDRAVRRNFVTAVKYLDKHPAVKVITGDCGFMMFFQELAQAYTRKPVALSSLMLITGLNQFIHPEKQICIVTANSGDL
EPLIPVVSRVCGMGLQVFADADHGHDGVGSMTGSMLDLVSGTGGAGYSNAPGSSTTGGGALALAAAAAQQQPHLP

SFAVGGPGMAPLQNYPFGPGIGMASAAPGGAPASAGAPMLSGTLGGGVGAPTSTSATAGTGPGRPAAGGQLAGSV
SYLEEKNPSVGSSSSIARTDSSALSTASSQPKQEQSKIPAPADQMNRTPVAEAPATSNSKSHEYSKAFQFVSHDGKT
MNLLAEKLQKLSLVDQDRGGEGEQNGDGKISKPQSKTHTTRRGQKVVLSQNNFLDGAFADELSSKLSALTQAFDAVE
QDGKVPGERPPLEAPLLQPGGEQAAEESAVTTQQDQKPPRAAAGREVGAGAAAAELRPAAHHGEHLSGATAQDEK
VDTDVERNGTSEGKIQGGRGKEEDAQGCARALTEVVTSSTTPGDHLTLSAATPAAAAFVDVTTVEHILDETTALAAQD
VFGAQPHEMRTLKEMFHDEQEARKAHEDPHAPSPELKPIAAGGGAPTSALGLGAGGGFSLGGGSSRGGSPGTGAA
PGAALGPTESVSSPRGTGSSVTSLNTSAGPSSIAPSANNSPLAGKAEPAAGGATGDAIFKLVPDAGAAPSGAPGQQN
VKRQWQLPSTGAFHQSSQQQGGAGEQQTASGKTAGSSLTEASVAGTGSATIGTGTSSQRSNESSSFVASASGSIAP
GTQIKLGTGDHPMAPTDSSSGHDFHIWHPRLQHLAGDTLQGPEQDISPCSPELKSIPKLSRTQKLTSIDPFAAGEQTV
RSLSNLDDTDLDSSSFSLSRHGDVFARSRGYHDRSLQGVGAHSMSGSLSLRGRAGGDSLGEYTSRTRQDGRTALG
ALLLSKASRAQAGGTNSPRDQNLADGIHLNFGRQAQTASIKSNSLEMTTSGAPSSNSKNSSLDVVVAGTARTSSSDL
RGSEGGLAARSDSNLSGGLPTTPSHANTNRSSSISGARAHRGSDDDRTSWTGRDGRGLALGQPDFSTMTDDASAA
SRNRMYPVVGLNNNAADNDPPARNRNAPGVLFSKLRGDNMRPNPLSSLLVGGATPTFPNAAASTRKDYLEDIHVPV
ALPEGLTPALTRVAEHGQTILGPQQGGKDASQATAGTLAQGRASVSDDQSYNLEPSSHSSSGAGMPFSAPPPPPSF
LPDTTSSILGPIEQDFILHDVLTPGERAATAMLGPPTAAHDATQLEAITEDKELVLVTSSVTNNSFLLDHHLDVASAQNS
QSLLNVSSSLDTSLAAAMAPAAVDENYPGGGGGATTAADHREISASFQQFLQKANETFQQQQKEKLRVAAAQAVDG
GSCAVAEEGNIADLGPATPDGSALLTYHSPVVGAVKSKSGSSSASLPQSQTVGAAEELRPSDVNYWGTLLGASNEM
RKQVRKKQSVDEMALQPGLSLLCRSASTDEFLRQQQAAAFANEAYPTSQTPFSTAGGGTIASSLTNLNLASLAGIMKP
PGGAPGGGAAGSSDNGHAAEHQVILAGGAHQHLARSNSKGSSHFNLAGGGTGTLHQQYGSSALLSRRDSRYLEVT
VVEHHQTIGNQINACAPQHASYPAAGVHPTPAWLQQRLNAAMLQPSTETRHIDRQRFVVMGIEDLDHFDGIAEGYPL
NFKDCEPLFLQKVNKLLELHPEIDCFLAECTQLPMFSDSVRALFGRPVYDIVCAAHMIMLGFLQSEHQVLEDGEKTQQ
NGYAYGYESGINGYGTAAGNIEGGINEAYIFGMELTLEERSQSAELQEFREMFAAAGIPLLEMKKVGGGGEKR

>Amoebophyra_ceratii3 jgi|Amoce1|7280|g5725.t1
MATGTGQLEAYSVAQRTLFLSLLLREGGTESQFPDAALPHTIPHEEIDRFLQASDGDCELAVRAFCGTLLWRRDTLRL
AYERRRREVESDGPPQQHPIVEANVHQQLTSGSPARLGLEVRPFVPLMNNGTTRFLGFSGFDRHPIVWSDARLSRP
EVVGADHQLAVDNTVYMMECLRRVCYARSLPFKIYVLMDFSGWSLKLASHRLLQRLLVTTLMQHYPETQLRTYIVNA
PALFQVAWRIFKAFAPANTVRKVSFLRLEEAKCRRVFQAVEEKRSRRESREVEREASRVGGELLAELVALGNSQEEL
EEFFLRTPREVLPKEYGGDLLFAPCSMGPGEEVFAGARYMDQELVEAQARANGLPARARDADADHRLLPVRRSGAE
GGDPQDKKPSAEFVSGFAHVDTRLHLQRISDSRQNESAEERALSKSKSSVAAGGLNGGTGGAANVVLPRGGFFAGA
PAPGCRTRPGKASTHNPLRVREISAIGGDHLPDTNVIPPRTVLPLSPAKAPQAVPCEYACKTCDGFDLEDCGSEAESA
HTFASSSTGDVINSAGNGTPGSPRSPRAALGNASRHPHAANALRQPATTSSNPFYVRRSAIVSSPEAARSVGPTSSP
HVGSQRTAARDVGRSAQDEFDSAFGDSQLACSPRIVHPLSEVGSPRALDNVSGVRDSSTAWGTSAGELAGGVQQR
VVENNARPLHGTSTPTGPAVPPTTANANAGTASSARSSGGNTPLQHQANIDQFPHHNLNHTRTTPQRVYNPRPVTF
VEVVEDNPAPTLGIVRIDHEYAPVAGNVAHPSSYAYRSVSLIVPGLTFQMCRKGQMSREVERAFYRVLDTLVFQFRVS
GITSDCGFMLYFQAKARAHLKSRSISHVPVFMSAFAHLPAVICGLDADAEVAILTANGRDLMPMLDLCLSGSGGAAGD
QEKAARGRASGPGGSSFSLSKMFLTPSRGGKASSPNGFHQGVATTSRDDRGRPWSPLGKDHDFDSDLSTQRRHG
HVDSARGHGGTSSTPDSRFVVVSCTRIPGLQEALECGRPLDREEVAHGLAVLTEQVLEAHPRVAVFLLECTELPVYS
DVIRRESGGMPVYDAITNADFFMSGALNNARFGSPMSEQVGGSCFYNSGGGGGGAAFSSAPGRVPGAATRKSSGE
SQESLTWPQGRVGGYGGGFESKYSLCAGDHAGPAFAAAPPPRRYQPASGPQPIMGPPPAQRYPPRNDFYGSYNG
YPSSNLGRRSPAHYVGDIAEIEQGGRNSADCAGGRYPQAEQDYRGLTVSESMGMVDDGSLTRRTSDHLLRSLAGLA
SMGKLGGDFPLDASSTSTRDTNTNTMTPDHGISYGNYNSNAYYAPAPGGLGAGTSTSKQSTDGEQPAVSTAKAPPG
SSVGQRNVELMEEFHRRLKRNLGSSEPVTSALIKRKFLQQTLGNPQCWARVQKKKKGNKDGSAAAGEDEDNGTLEV
GDVWAMDALDALKSDLGRKNNTQRKLAHGGGWWSATGKWKVVESVKSTRLKEKQQREAEKAEEARRKEEQWQK
QEEEKLRAKFNEDVKEKGINVGGIAFSNAGPGAGQNFDAGENSDSDSGYSDDENNSKANKKGGSAGAGGQAALQII
NSMAELLTMELGSLTGYGLYGDGKKHPASVSIQDLDLVVDWTRVEPFLLNAPELRPFLSRWKYLSLASRQLRTVAEY
EERWMKLEALHPLNDLNWLADQFVFLHLSREYRELGFALDTNPDYKPKVAAANGNGNGNRFGKNGEDEGPGGTAE
NPAAEQTGENANDAGAANADIKGGSKVRFARAADGQTAPEKLDLTDDRFFERASNDSSYRSLLEKEHLALTDPDAFK
PGQFGWISLLELQAVCGAFIELRFPDRNLYIEDSMMTQAMFLKGLEIAFSMTAPVSTVSVPFEVSFDTACVVCEICRIL
WLRFQKQGGWGKREVSRLEEIFPVDSGDVSNARVFTLLQKGGIPVDAGEEQTGFIDLLRESDINQDGVTSFEEFCYLL
KRFDDTLRRQEVARIVEHGYVDLLPGTFLDEWGYILKSRLRGGKGSSTAATTSKNASSSYQYHSIADLRDALTTLGLE
QISRERLRQMDKLREELEIDARQMDDYDICFLLGACFQRNTGDFAQLLREWAYEREKKRMGKHAREKKDLLKPTIKLS
VMDHLDDVLSQDFKSDWWFNVAKKTLQLSIWEHERPHFLTQKISGKHGGGSAALKAAKERKSAGTEGSTVTMGILSE
TPGFARAETNENLAEARRTATMGISDYLFSSEPAPFKPVNNLGNQFDTIVQKGFGSPSSPLLWGYSSRNSAPNYIVSS

DIFNIDNTCFDNSLRCPKNAQQNLKNVLFSAGKSPPSKQELNKVRHNYQFELEYSAAKVNSGTGKEEAPATARTAPP
QILHPFFEFKNPVFGPVHVQLREVGYLKGGGTEADNSQDPSSSVDETHNGASVSEIKASLKKGKYVKYGASNPTTQA
KGYFPVVHFDQRFMVGRLGLEEWDKAIYLKDVLPVRLEKMGLSDFVLENGAGRVLESSAKGIFGGGGGSTVTANNV
QTAELGLSRHLRPFSQGSELALRQKVVLHPEMVLSEDDRDVLNALVSDYADNMKRLQAGKETAVDADGRSVLKDILY
NIHTARPLEEAIEDVLKKTKMQAEATGTVSGTVANVASYFGMAGKTVKTAIFKRLFGIDSRAVRKVEVKNIPAAATSAG
EGAEPAASAKKPTRTGALYFLPTTVLQPSLPPQNRGSGDFLLPICSNVARDKSRVMELRVDTEDDKSLSKQSVASGRI
HQSALVVVRETSAIAQCLCNGMHKGCSAPGWPPLEKPYFSEQSSKLMQADLEERLGAREEAAAATALLEVARDLQQ
SEKHMVTQETPDYPPSLDAAGEAHEPFVRVGAEGPLDQKAIIREATQVRVEPNVVMPAPPEMPSSYYADQEKRGGP
FDEHGQVFDVDMQGQKKAWENNEVEERAKLNEYQSRLKRLMVLVGEQE

>G.foliaceum_A CAMPEP_0188240084 /NCGR_PEP_ID=Glenodinium-foliaceum-CCAP1116_3-20130913|6335_1
/TAXON_ID=160619 /ORGANISM="Glenodinium-foliaceum-CCAP1116_3" /LENGTH=332
/DNA_ID=CAMNT_0031571451 /DNA_START=1 /DNA_END=999 /DNA_ORIENTATION=+
FGLNDWQAEWDGVQDEYQFGQELIEADRRELVNKVDNKKPXXXXXXXXXXXXXXXMEKIKKQLIKQQAPSLGVVRL
DYNYPPAYGDIDCPGSYDYDVIYRCVPGLTFEMAQSGKMTFAVQKEFEAAIKYLEGKGVSGITGDCGFMMAFQPLAR
DIAKVPVFMSSMVQSPMISVAFDKYDQILILTANSKTLAPQKEVLLNHCGFDVDDDRFLIYGCQDVPGFDAVAKGEKVD
IEKVTPGMVAMVRKLLEEMPSLRAILLECTELPPYSDALRAEFGLPVFDAITCADFFISARKDNPRFGLNQWQNDWDG
TVDEYQFGQNLSQQERGRLLNK

>G.foliaceum_B CAMPEP_0188434408 /NCGR_PEP_ID=Glenodinium-foliaceum-CCAP1116_3-
20130913|269394_1 /TAXON_ID=160619 /ORGANISM="Glenodinium-foliaceum-CCAP1116_3" /LENGTH=882
/DNA_ID=CAMNT_0031778853 /DNA_START=1 /DNA_END=2643 /DNA_ORIENTATION=-
XFAAMMQHPGMMDHHMAQRLDLLEQELLQVKMQTSMGHPMMMHGGYSPSPMAHMANSRAISPMMGMPGQMPM
QMMPMQQPYMPYGSHDAMFQTHKRIEEMHKKLVESMKLTEERHEKLLASHGFLMDTHKKMMSSRVGLGKDAKKDA
KQHPKLGVVRLDYNYPPGAGDIDSPASFGYDVFYRVVPGLTFEMAQSGKFTEAVERNFAEAIKWLEQKGASAITGDC
GFMMAXRNFAEAIKWLEQKGASAITGDCGFMMAFQVIASKIATRPIFMSSMVQCPVIATAYDPNDQILILTANDQSLEP
QKDVLLNKCGFDVEEDRFLIYGCQSVPGFEAVAKGEAVPLDKVQPGIVEDDDGNHQEKPEHRRHPPRVHGAAALRG
RAPLTMEIIKKNRNIAGILLECTELPPYADALRAATGLAVWDAITGADFYINAYKDNPRFGLNDWQAEWDGVQDEYQF
GQELIEADRRELVNKVDNKKPGAAPKTKSKAKSKAAMEKIKKQLIKQQAPSLGVVRLDYNYPPAYGDIDCPGSYDYDV
IYRCVPGLTFEMAQSGKMSYTVQKEFEAAIKYLEGKGVSGITGDCGFMMAFQPFARDVATVPVFMSSMLQSPLISVAF
DKYDKVLILTANSKTLEPQKQILLNECGFDVEDDRFVIVGCQDIPGFDEVAKGGKVDLEKVTPGMIKLALEVLDQQPSL
RAILLECTELPPYSDALRMATGLPVWDAITCADYFISSRKDNPRFGLNDWQFDWDGQQDEYTWGANVPKNEAKYLQ
HVGEVASAFAAQRCRELHCAHVALAAFVNHRDAMSAGVHLVAWRLEGCASRIRSAGIAPQIIHPVRVAKFGECTAPG
MPEPSSLDKLERADLAREAPLGRAQTRDAPNVCGAVX

>G.foliaceum_C CAMPEP_0188294966 /NCGR_PEP_ID=Glenodinium-foliaceum-CCAP1116_3-20130913|39119_1
/TAXON_ID=160619 /ORGANISM="Glenodinium-foliaceum-CCAP1116_3" /LENGTH=653
/DNA_ID=CAMNT_0031629459 /DNA_START=1 /DNA_END=1960 /DNA_ORIENTATION=-
XASSMSGFGNVPQYGSPAAGGVSIDHHHAVYAMYLDAMERLQQLHDQHSQLMSSHLELMEAHSQVLGYAGDLEVA
GATGGGEKQKKKKPLKKHPCLGVIRLDYNYPVAEGDTDCAASFGYDVIYRVVPGLTFEMAQAGKVTETVERRFAEAI
KYLEGKGVNGITGDCGFMMAFQVLARKIASAPVFMSSMVQCSIIAAAFDENDRILILTANDKSLEPQADILFNHCGIRVD
DDRFLIRGCENLPGFDAVAAGKAVPIHIVQPAIVKMVDDILRREKRIGGILLECTELPPYADALRASTNLPVWDAITACDF
YVSGYKDNPRFGINDWQAEWDQQQDQYVFGQNLLESEKKNLQMKALSEADLKQAQVAKAKAKAAAKLKKLAKKQA
PSLGVIRLDYNYPPAAGDIDAPGSYGYDVIYRVVPGLTFDMAQSGMMTYQVQQRFGEAIKWLEAKGVCGITGDCGFM
MAFQPFARDVATVPVFMSSMLQSPLISVAFDKYDKVLILTANSKTLEPQKQVLMNECGFDVDDDRFVIVGCQDVPGF
DEVAKGGKVDVEKVTPGMIKLALEVLDQQPSLRAILLECTELPPYSDALRMATGLPVWDAITCADYFISSRKDNPRFGL
NDWQFDWDGVQDEYTFGANVPKNEMRYMKNKA

>G.foliaceum_D CAMPEP_0188425482 /NCGR_PEP_ID=Glenodinium-foliaceum-CCAP1116_3-
20130913|264918_1 /TAXON_ID=160619 /ORGANISM="Glenodinium-foliaceum-CCAP1116_3" /LENGTH=879
/DNA_ID=CAMNT_0031769897 /DNA_START=1 /DNA_END=2636 /DNA_ORIENTATION=+
GGSMRAAGAQIAMFHQAQQGAMAAPIAWGTPRVHHGGGVVSLVPGAPGVHDRLDMLEDELSLMRAELGNSRMTK
VLSPTVSYQAPQVQQQMVPAPRYMPQEVADPGPGAGPQEPVVALKDYQQLHQMHRDLVTKMVELQGRHSELLGN
HHQLIMAHADVLNMAANMQPAVAAIANDPAKAEDADKKKHPTLGVIRVDYNYPAAKGDVDCPASFGYQVIYRVIPGLT

FEMAQSGKFTEAVERRFAEGIKFLEQKKVHAISSDCGLLMAFQVLARKIAAAPVFMSAMVQCPIIAAAFDPADRILVLTL
DGPVLKKQKEVLMSTCGFDVDENRFIIQGCKHLPGFDAVDRGEPVPVDLAQPAIIKQTMECLKSNFSLRGILLESTELP
AFADALRAATELPVWDSITCADFFISGYKDNPRFGIDDWQKEFDEEMKNDYAFGDNLTKKEQDALVTNTAQAKAAAK
GKAKAKSKAMLKKLKKNAAKAKAPVLGVLRLDYNYPPAAGDIDCPGSFDYDVIFRMVPGLTFEMAQSGNMSYTVQQ
QFIGAIKWLESKGVCGITGDCGFMMAFQPIAAGRRQRARVLVLDDTLESKGVCGITGDCGFMMAFQPIARDVASVPV
FLSSMIQSPMVSVAFDKYDKILILTANSMTLQPQKEILLSQCGFDVDDDRFIIVGCQDVPGFDAVAKGTKVDVEKVTPGI
CDLVADLLSKQPSIRAILMECTELPPYSDALRATTGLPVWDAITCADFFVSCRKDNPRFGLNQWQNDWDGVVEEYTL
GQNLTDSQRSLGGSYARPPALLARAHALRSRLGIRMFHYLPPVRRPGSTRVANATMRALPAAPPAPGNRPGVAGRA
CGRPNIGRLCAPRARARAARVRLVLPFLPPX

>G.foliaceum_E CAMPEP_0188426016 /NCGR_PEP_ID=Glenodinium-foliaceum-CCAP1116_3-20130913|265187_1 /TAXON_ID=160619 /ORGANISM="Glenodinium-foliaceum-CCAP1116_3" /LENGTH=856 /DNA_ID=CAMNT_0031770433 /DNA_START=1 /DNA_END=2566 /DNA_ORIENTATION=-
XLLEQELLHMKIVGAGPPMPGQSQAMSPMSQAPHMSSMYPAPAQSIQAHVSAPQGHFDPSAMANHHAELVSCHKK
LADTMKQQEDHHAKLLQSHALIMDAHKKLMSAQVGLGKNAKKAARGHPKLGVVRLDYNYPPAKGDTDSPASFGYDV
FYRVVPGFTFEMAQRGQFTEAVERAFAEAIKYLELKGANAITGDCGFMMAFQVLARKIATRPIFMSSMVQCPIIACAYD
PSEQILILTANGKSLKPQKDVLLNSCGFDVDEDRFIIRGCQDIPGFDAVAKGEAVPLEIVQPGIVKMTMQILKENPLIQAIL
LECTELPPYADALRAATGLCVWDAITAADFYINAYKDNPRFGLNDWQAEWDGVQDEYNFGDNLIEADKRKLVNQVPL
GQQAQAKAKAKAKVKSKAMMQKIQKKLIKQQAPSLGVVRLXXXXXXXXXXXXXXXXLGVVRLDYNYPPSAGDIDAPG
SFAYDVIYRVVPGLTFEMAQAGKMTPPVQKEFETAIKWLEAKGVAGITGDCGFMMAFQPIARAIAQVPIFMSSMVQSP
MIDVAFDKYDQILVLTANSNTLGPQKEVLLSHCGFDVDDDRFVIHGCQDVPGFEAVAKGEKVDVAYVTPGIVKEVAAIL
ERIPSLRAILLECTELPPYSDALRKEFDLPVFDAITCADFFISARKDNPRFGLNQWQNDWDGTIEEYSFGQNLTDKSGA
GFSTLEERVATRLLRVSVRARAAEXPVVPRPSTRRGGDKPRCASGPTQADPKRHPTPIRPSSPTPAPIRAQGPGDGP
LFVHEAACLLCGPCRRGSSSPARVGRSRSASRRDRYNGVASCAREGSRPMFCRNSAFQGTRVQKQSTNEEDAEDK
KX

>G.foliaceum_F CAMPEP_0188370660 /NCGR_PEP_ID=Glenodinium-foliaceum-CCAP1116_3-20130913|162165_1 /TAXON_ID=160619 /ORGANISM="Glenodinium-foliaceum-CCAP1116_3" /LENGTH=529 /DNA_ID=CAMNT_0031709055 /DNA_START=99 /DNA_END=1685 /DNA_ORIENTATION=-
MMQHPGMMHHHHMAQRLDVLEHELLQVKMQTGGHPMMMHGGSHYMGGMSPMGHPGIMGDSRGMSPMMMGG
GSPAPMQHAYMPNGSHDVMFMTHKRIEETHKQLVESMRLTQERHEKLLESHGFLMETHKKMMASRVGLGKNAKKD
AKQHPKLGVVRLDYNYPPAAGDIDAPGSFGYDVFYRVVPGLTFDMAQSGKFTESVERNFAEAIKWLEQKGACAITGD
CGFMMAFQVIARKIATRPIFMSSMVQCPVIATAYDPKEQILILTANGKSLEPQKDVLLNSCGFDVEEDRFLIYGCQDVP
GFDAVARGEAVPLDVVQPGIVKMTLDIIKKNPRIAGILLECTELPPYADALRAAATELPVWDSITCADFFISGYKDNPRF
GIDDWQKEFDEEMKNDYAFGDNLTKKEQDALVTNTAQAKAAAKGKAKAKSKAMLKKLKKNAAKAKAPVLGVLRLDY
NYPPAAGDIDCPGSFDYDVIFRMVPGLTFEMAQSGNMSYTVQQQFIGAIKWLESKGVCGITGDCGFMMAFQP

>G.foliaceum_G CAMPEP_0188248466 /NCGR_PEP_ID=Glenodinium-foliaceum-CCAP1116_3-20130913|11173_1 /TAXON_ID=160619 /ORGANISM="Glenodinium-foliaceum-CCAP1116_3" /LENGTH=563 /DNA_ID=CAMNT_0031580203 /DNA_START=1 /DNA_END=1688 /DNA_ORIENTATION=+
XTRLGSPRHSGTSGTSRASDVCCCCIGRSSAAFRAMMLPGMMMQQSHYGSMTPSVNGNSSSPSRLDMLELELNQL
TAQQGGGYPQPSFDKMGMTGMGTSMGGFGTTHQFGSPPAGGVSVDHHHQVYAMYLDAMERLQELHLQHSQLMN
SHLELMAAHSEVLGYAGDLEVTGASGEKAAKKKAPKKHPCLGVIRLDYNYPVAEGDTDCAASFGYDVIYRVVPGLTFE
MAQAGKVTEAVERRFAEAIKFLEGKGVNGITGDCGFMMAFQVLARKIATAPVFMSSMVQCSIIAAAFDNNDRILILTAN
DKSLEPQKDVLLNHCGFRADDDRFLIRGCENLPGFDAVAKGQAVPIHIVQPAIVKMVEDIRKRERNIAGILLECTELPPY
ADALRASTDLPVWDAITACDFYVSGYKDNPRFGINDWQAEWDQQQDEYKFGANLLESEKKHVQMHDLTPEQLKQA
QVAKAKAKAAAKLKKLAKKQAPSLGVIRLDYNYPPAAGDIDAPGSYGYDVIYRVVPGLSFEMAQSGMMTYQVQQRFG
EAIKWLESKGVSGITGDCGFMMAFQP

>G.foliaceum_H CAMPEP_0188362200 /NCGR_PEP_ID=Glenodinium-foliaceum-CCAP1116_3-20130913|144842_1 /TAXON_ID=160619 /ORGANISM="Glenodinium-foliaceum-CCAP1116_3" /LENGTH=323 /DNA_ID=CAMNT_0031700217 /DNA_START=1 /DNA_END=972 /DNA_ORIENTATION=-
GATCRAAGAALVAGGSVFGRRARQALAQAASPTPAAQAGRKRRAVSASPTEDLQRQLDPPCLGIVRLDYNYPPSAG
DVDFPASFGYRVIYRAVPGLTFDMAKSGAMTGGVWEEFKSAIAWLEEKGVCGITGDCGFMMAYQSLARQVAEVPVF

LSSLVQCPMISAVYDTWDRILILTADSKSLEPQKELLLAQCGFSLDDERFIVRGCEDVDGFEAVKNGEKVDEDRVTPG
VRRVVDKALHDFPTIRAIVLECTELPMFGDALRREYRMPVFDAITCADFFIASLSDNPRCGADSWQKKWDGRIEDYKL
GDNLPPESRAKLING
>G.foliaceum_I CAMPEP_0188441342 /NCGR_PEP_ID=Glenodinium-foliaceum-CCAP1116_3-20130913|272877_1
/TAXON_ID=160619 /ORGANISM="Glenodinium-foliaceum-CCAP1116_3" /LENGTH=355
/DNA_ID=CAMNT_0031785811 /DNA_START=1 /DNA_END=1061 /DNA_ORIENTATION=+
XAHRLDVLEQELLHMKMAGGFMPLPGQPPSSPMMAPPGHYMAPGPPATAQYPPPAQPAGPLTMDPGMMSAHHKE
LMDCHKALFDMMKKQEDHHTKVLQSHSLIMDAHKKLMAQQVGLGKNAKKAAKSHPKLGIVRLDYNYPPAKGDTDCP
ASFGYDIFYRVVPGFTFEMAQAGKFTEAVERNFAEAIKFLELKGANAITGDCGFMMAFQVLARKIATRPIFMSSMVQC
PVIACAYDPKELILILTATXFMSSMVQCPVIACAYDPKELILILTANGESLRPQKDVLLNSCGFDVNEDRFVIKGCQDVPG
FDAVAKGEAVPIDVVQPGIVKMTMEILKQRPLIQCILLECTELPPX

>G.foliaceum_J CAMPEP_0188292990 /NCGR_PEP_ID=Glenodinium-foliaceum-CCAP1116_3-20130913|38017_1
/TAXON_ID=160619 /ORGANISM="Glenodinium-foliaceum-CCAP1116_3" /LENGTH=287
/DNA_ID=CAMNT_0031627375 /DNA_START=1 /DNA_END=864 /DNA_ORIENTATION=-
RVNGQMKVGILRIDYDYPPIPGDIACEASYGYQVVFAKVEGLTFEMAQRAVLDDHIMASFQKAVAALEAESVVAITGDC
GFMMAYQPYVRALTKLPVILSSIIQAPAMAAMHSDDAVFALLTANSRTFDKKRLLSQSGVNVDLDNWVLVGLQDVPGF
EAVALGEKVPAEKVQPSIAAMCRDLQARHPKLAGFLLECTELPHYADEIRAKTGLPVFDAITLVDYFQVACCSPPSFSQ
SVLEGVVGALDPRVAASAVAQITFGILGTVVMVPVVVGTSAVKQSSCCTVL

>G.foliaceum_K CAMPEP_0188423942 /NCGR_PEP_ID=Glenodinium-foliaceum-CCAP1116_3-
20130913|264149_1 /TAXON_ID=160619 /ORGANISM="Glenodinium-foliaceum-CCAP1116_3" /LENGTH=752
/DNA_ID=CAMNT_0031768359 /DNA_START=1 /DNA_END=2255 /DNA_ORIENTATION=+
XRSAAEMSGAHPELQKLAPKIGVLHFETSPEASGQTAQARTHAAGFGYEVVERPVPGLNILNVASPTERVKAELRRA
VDELVSLGVKGIAGASHQMWPLQEHVACMARVPVAMSSLMQIALLVPAHHVDQIFLIIATHAEQLKLDLGSVLKSRGL
QFSPARFVVHEPSAEELDDPAKLAERIKGLQDSDNRIAGAIFDCGQLIRHADRVRAISSFSVFDQITCLDFLISASSDNP
YFGTSYGTGAPPVELLVKAKRQLMSPSEEYVRALRSRVNGQMKVGILRIDYSYPPIPGDIACEASYGYKVTFAKVDGL
TFEKAQRAALDDQIMESFERAVKALEAEGVVAITGDCGFMMAYQPYVRTLTKLPVVLSSIIQAPAMAATHSSSAVFALL
TANSSTFDKRLLLTQSGINVDLDNWVLVGLQDVPGFEAVALGEKVLAEKVQPAIAAMCKDLKARHPTLAGFLLECTEL
PHYADEIRAQTGLPVFDAITLVDYFQVACCSPPSFSQAVVQGFLGALDPRVGASAVAQITVGILGTVVMVPAMAGSAQ
QXVLLRRVVGAPRPMPRVRACGSDRARRXXXXXXXALQTFLESRPQAQARTRGARARGAVVALSSHVPRVRAPPV
VAHLCAAGARDSVCVWRLLRVQAQGXPPSADAESLWLTRGHGLLACEFKGASAAGALRRCVCSLAMYRCVMAVEH
FETPDRWGLALARNGRTKFGRRRGMQIHVRAIALPSMCSGGGPNRPPRRLTQSC

>D.baltica_A CAMPEP_0200040072 /NCGR_PEP_ID=Durinskia-baltica-CSIRO_CS-38-20140214|157896_1
/ASSEMBLY_ACC=CAM_ASM_001176 /TAXON_ID=400756 /ORGANISM="Durinskia baltica, Strain CSIRO CS-38"
/LENGTH=710 /DNA_ID=CAMNT_0045966521 /DNA_START=148 /DNA_END=2280 /DNA_ORIENTATION=+
MMPAAVPMDHQFHAMTPITTRAISAGSTPTRLDLLEHEVNRLAQQQTMDGQGGYPISQSMALMKPHAMNSSPMSDA
STSASGGSVSMEEHTQVLKLYLDAVSHLRDLHDQHTSLMEHHAALMKMHTDVIQYTTELEGKVTGGAGGKPKKELK
KHPGMGVIRLDYNYPAADGDTDSPASFGYDVTYRVVPGLTFEMAQAGKFTEQVERRFAEAIKFLEQKGVNGITGDCG
FMMAFQVIARKIATVPVFMSSMVQCPIIAAAFDKNDKILILTANAKTLQPQKEVLLEHCGFQVEDRRFVIEGCQDLPGFD
AVAKGEKVPIDIVQPAIVKMTMGILKREPKIAGILLECTELPPYADALRAATDLPVWDAITACDFYISGYKDNPRFGIDDW
QAEWDKQQEEYSFGQNLIDTEKALLVNKIDAKKAVAKTKAKAKAAAKVKKLAKKQAPILGIVRLDYNYPPAAGDIDHPG
SYNYDILFRCVPGLTFEMAQSGMMTYQVQREFADAIKWLESKGVAGITGDCGFMMAFQPFARDVATVPVFMSSMVQ
SPMISVAYDKYDKVLILTANSKTLLPQKNVLLSECGFDVDDDRFVIEGCQDVPGFDAVARGDRVDVEYVTPGIVKKVTT
LLANQPQIRAILLECTELPPYSDALRKATGLPVWDAITCADYFISSRKDNPRFGLNDWQFGWDGQQDEYTFGENLSA
QQRSHLLNG

>D.baltica_B AMPEP_0199916896 /NCGR_PEP_ID=Durinskia-baltica-CSIRO_CS-38-20140214|5581_1
/ASSEMBLY_ACC=CAM_ASM_001176 /TAXON_ID=400756 /ORGANISM="Durinskia baltica, Strain CSIRO CS-38"
/LENGTH=735 /DNA_ID=CAMNT_0045836133 /DNA_START=1 /DNA_END=2206 /DNA_ORIENTATION=+
XAYAPSALSSLSFEEAAMFHHYPVMMVSSPFPMDHSMTPMTPVSARALSLSPTPTRLDLLEHEVNRLAQQQTVASH
GGYPMNQSLALMKPQAMDHSPMSAASTTASAGSVSIEEHTQVLKLYLDAVSHLHKLQDQHTTLMEHHAALMQMHTE
VIQYTTELEGKVSGAAGGKPKKALKKHPCMGVIRLDYNYPAADGDTDSPASFGYDVTYRVVPGLTFEMAQAGKFTEQ
VERRFAEAIKYLEQKGVNGITGDCGFMMAFQVVARKIATVPVFMSSMVQCPIIAAAFDPNDKILILTANGNSLKPQKEVL
LEHCGFQVEDERFIIEGCQDLPGFDAVAKGEKVPIDIVQPAIVKMVMGILKRERRIAGILLECTELPPYADALRAATDLPV
WDAITACDFYISGFKDNPRFGVDDWQAEWDQQQEEYTFGQNLIESEKALLVNKVDAKKAAAKAKAKAKAAAKVKKLV
KKQSPTLGVVRLDYNYPPAAGDIDHPGSYGYDVLFRCVPGLTFEMAQSGMMTYAVQREFADAIKWLEAKGVAGITG
DCGFMMAFQPFARDVATVPVFMSSMVQSPLISVAYDKYDKVLVLTANSKTLLPQKNVLLSECGFDVDDDRFVIEGCQ

DVPGFEAVAKGEKVDVEKVTPGIVKKVMDLLSNQPQIRAILLECTELPPYSDALRKATGLPVWDAITCADYFISSRKDN
PRFGLNDWQFGWDGQQDEYTFGQNLSAQQRSHLLNG
>D.baltica_C CAMPEP_0200040918 /NCGR_PEP_ID=Durinskia-baltica-CSIRO_CS-38-20140214|160103_1
/ASSEMBLY_ACC=CAM_ASM_001176 /TAXON_ID=400756 /ORGANISM="Durinskia baltica, Strain CSIRO CS-38"
/LENGTH=549 /DNA_ID=CAMNT_0045967419 /DNA_START=1 /DNA_END=1647 /DNA_ORIENTATION=-
VRPAAAPPAALIGRDDVLDDDPHMVNQRLDLLEQELMRAKMSGSQHAMMPSMQQAPQTPMRAMPQAHGFQGAAS
PQTPYMPYGNQDVMFHSHKEIMETHAKLMESMRKQAERHDKLLESHGMLMDAHKKMMASRVGLGKDAKKEAKKH
PVLGVVRLDYNYPPAAGDVDSPASFGYDVLFRVCPGLTFEMAQRGKFTEAVERNFAEAIKYLELKGASAITGDCGFM
MAFQVLARKIATKPVFMSSLVQCPIIAAAFDPKDMILILTANGKSLKPQKDILLNSCGFDVEEDRFIIEGCQDVPGFDAVA
KGEAVPVDVVQPGIVKLTLGILKRNPRIAGILLECTELPPYSDALRAHTGLPVWDAITGADFYINAYKDNPRFGLNDWQ
AEWDGVQEDYFFGDNLIEADKKELVNKIEKPKPKPKAKTKAMMEKIKKKIVKQQAPTLGIVRLDYNYPPAAGDIDCPAS
YDYDIIYRAVPGLTFEMAQSGKMTYAVQKEFEAAIKWLESKNVCGITGDCGFMMAFQPIARDIAKVPVFMSSMVQCP
MVSVAFDX

>D.baltica_D CAMPEP_0200039960 /NCGR_PEP_ID=Durinskia-baltica-CSIRO_CS-38-20140214|157677_1
/ASSEMBLY_ACC=CAM_ASM_001176 /TAXON_ID=400756 /ORGANISM="Durinskia baltica, Strain CSIRO CS-38"
/LENGTH=637 /DNA_ID=CAMNT_0045966403 /DNA_START=1 /DNA_END=1910 /DNA_ORIENTATION=+
QALTDRTRMPRPTAGGDNTSAGAPDAAAGAHIAMQHVPMQAPVAWGTPRVHPSSMPTPGGGVHDRLDMLESELS
QMRSELGHHRATAVMETRFVAPPLALPQLQNHPHAHAHMYSTPPSKGQHPFATPSTVDTALMITDSSPSTNATNDQ
AEPVVSYKDYHFLHQMHKDLIIKMSELQNRHNALLNNHSQLMSAHADVLNMAADLHTKVLTAGGAAAPRAAPKKEDN
RKKHPSLGIIRLDYNYPPAEGDVDCPASFGYDVIFRVVPGLTFEMAQQGKFTEAVERRFAEAIKFLEHKKVNAITGDCG
FMMAFQVIARKIATAPVFMSSMVQCPIIAAAFDPKDKILILTANDLSLKPQKDVLLNSCGFDVDEGRFVIKGCQNVPGFD
AVAKGEAVPIDIVQPGIVKTTMDTLRADYNIKAILLECTELPPYADALRAATELPVWDAITAADFYVSGYKDNPRFGIDD
WQKEWDQEYEEYKFGDNLTLMDQSLLLQKDKQQPAPKAKAKAKAKTKAMVKKMKKIAAKAKAPVLGVIRLDYNYPP
AAGDIDCPGSYDYDVLFRMVPGLTFEMAQSGRLSYTVQQQFVAAVKWLEAKGVCGITGDCGFMMAFQPLARDVAN
VPVFMSSMVQCPMVSVAFDX

>D.baltica_E CAMPEP_0200084984 /NCGR_PEP_ID=Durinskia-baltica-CSIRO_CS-38-20140214|238428_1
/ASSEMBLY_ACC=CAM_ASM_001176 /TAXON_ID=400756 /ORGANISM="Durinskia baltica, Strain CSIRO CS-38"
/LENGTH=792 /DNA_ID=CAMNT_0046017323 /DNA_START=1 /DNA_END=2378 /DNA_ORIENTATION=+
XACARAMYAPRQYGAGDGNMSQRLDVLEQELLNMKASSLQMSTPGRYPQSQAFGGSPQMLPGQIGSPGASPSSA
PQPQYRHIGQEHIANHHKELLDCHKLLTEAMDQQEAHHTKLAASHEALLAAHKKFMAAQVGLGKGAKKAAKGHPKLG
IVRLDYNYPPAAGDTDCPASFGYDVFYRVVPGLTFEMAQAGKFTEAVERNFAEAIKFLEMKGANAITGDCGFMMAFQ
VIARKIATRRSSGRRCGDTDCPASFGYDVFYRVVPGLTFEMAQAGKFTEAVERNFAEAIKFLEMKGANAITGDCGFM
MAFQVIARKIATRPVFMSSMVQCPMIACAFDPKERILILTANGKSLAPQKDVLLNSCGFDVDEDRFIIEGCQDIPGFDAV
AKGEKVPVDIVQPGIVKKTLDILKRVPKISGILLECTELPPYADALRAHTGLPVWDAITAADFYINAYKDNPRFGLNDWQ
DEWDGEHEAYTFGDNLIESDKKELVNKIPDKVPGQPDPKIKAKAKAKSKAIMQKIKKRIIKQQAPVLGVIRLDYNYPPAA
GDIDAPGSFDYDVLYRAVPGLTFEMAQSGKMTYTVQKEFEAAVRWLESRNVCGITGDCGFMMAFQPIARDIAKVPVF
MSSMVQSPMISVAFDKYDQILVLTANSNTLKPQKEILLSQCGFDVDDDRFVIYGCQDVPGFDAVAKGEKVDVEYVTPG
IVKMVRGILDKAPQIRAILLECTEMPPYADALRKEFDFPVFDAITCTDFFVSARKDNPRFGLNQWQNDWDGTVDEYTF
GENLTSHQRARLLN

>D.baltica_F CAMPEP_0200069582 /NCGR_PEP_ID=Durinskia-baltica-CSIRO_CS-38-20140214|230672_1
/ASSEMBLY_ACC=CAM_ASM_001176 /TAXON_ID=400756 /ORGANISM="Durinskia baltica, Strain CSIRO CS-38"
/LENGTH=1010 /DNA_ID=CAMNT_0046001823 /DNA_START=1 /DNA_END=3030 /DNA_ORIENTATION=+
RLDLLEQELMHAKVNGGLHHGQMHGSPMTSMQGASPPPSYMPPPSYMPHGSQDIMFHSHKEIMATHAKLVESMKK
QAERHDKLLASHGMLIEAHKKMMASRVGLGKDAKKEAKKHPMLGVVRLDYNYPPAAGDTDSPASFGYDVTYRVCPG
LTFEMAQAGKVTEAVERNFAEAIKFLEMKGASAITGDCGFMMAFQVLARKVATKPVFMSSMVQCPIIATAFDPKDQILI
LTANGKSLQPQKDVLLNSCGFDVEEDRFLIYGCQDVPGFDAVAKGEAVPVDVVQPGIVKMTQDILKKNPKIAAILLECT
ELPPYSDALRAYTGLPVWDAITAADFYINAYKDNPRFGLNDWQAEWDGVQDDYSFGDNLIEADRKELVNKIEKPKPK
PKAKAKTKAMMEKIKKKIVKQQAPTLGIVRLDYNYPPAAGDIDCPASYDYDILYRVVPGLTFEMAQSGKMTYTVQKEFE
TAIKWLESKGVCGITGDCGFMMAFQPIARDIAKVPVFMSSMVQCPMVSVAFDKYDQILILTANSNTLAPQKEILLNQCG
FDVDDDRFLIYGCQDVPGFDAVAKGEKVDVEYVTPGIVKKVKGILSQVPTIRAILLECTELPPYSDALRKEFDLPVFDAI
TCADFFVSARKDNPRFGLNQWQNDWDGIVDAYEFGQNLTDSQRAKMLNXALRPSSGGQGQLTFGIAARGQRRLMV
SVAFDKYDQILILTANSATLQPQKEILLSQCGFDVDDDRFIIVGCQDVPGFDAVAKGTKVDVEAVTPGIVKLVQGILEKQ
PNIRAIVSECTELPPYSDALRMTTGLPVWDAITCADYFVSCRKDNPRFGLNQWQNDWDGVVDEYKLGENLTAAQRG
AAKMCESSVLASGQRLPSSSMWLHLSVHPFAXNLVTDAFVLTVGNSGRLERTSLCRNLVTDAFVLTGERRGAPKVPP
HDGTERRTSASPVAHRLPELFMPHADLKTLWSIEACAGGAGRQAVPRGRGAPSLAALPVTRQGGDRGGPRAARAA
RKK

>D.baltica_G CAMPEP_0199917990 /NCGR_PEP_ID=Durinskia-baltica-CSIRO_CS-38-20140214|6229_1
/ASSEMBLY_ACC=CAM_ASM_001176 /TAXON_ID=400756 /ORGANISM="Durinskia baltica, Strain CSIRO CS-38"
/LENGTH=282 /DNA_ID=CAMNT_0045837283 /DNA_START=1 /DNA_END=847 /DNA_ORIENTATION=+
XMGKPAMPMGGGKAKHASLGVIRLDYDYPPAPGDIDHPGSFGYDVFYRVVPGMTFEMCQSGKLTPQVQANFIEAVK
WLEAKGVSGITGDCGFMMYFQQLARRNTRKPVFMSALAQLPAVTCAFAYNEHIAILTANGQSLAPMRQLIKDECGVD
PDEHRYIFVGCEDVPGFDAVARGEKVDVQAVTPGMVAKSIRVVKQNPTIRAILLECTEMPPYADAIRQATGLPVFDAIT
ACDFFISSFKDNARFGINDWQQSWDGQQDSYKFGQNLSSAQRGMLVNKIR
>D.baltica_H CAMPEP_0200084306 /NCGR_PEP_ID=Durinskia-baltica-CSIRO_CS-38-20140214|238089_1
/ASSEMBLY_ACC=CAM_ASM_001176 /TAXON_ID=400756 /ORGANISM="Durinskia baltica, Strain CSIRO CS-38"
/LENGTH=351 /DNA_ID=CAMNT_0046016645 /DNA_START=1 /DNA_END=1049 /DNA_ORIENTATION=-
XESQNGRIAGAIFDTGDLIEHADKVRAVTKYPIFDQVTLLDYFVSACSDNPFFGTSYRSGDSPDALYTKAMRRLLSPSE
AYLRDMKSKIDRHIKIGVLRLDYSYPPIPGDVACEASYGYQVVFFTVEGLTFEKAQRDXEVQDRQVVFFTVEGLTFEKA
QAATLDDKIKAGFERGVAALEEQGVAAITGDCGFMMSYQTYVRALAKVPVILSSIVQAPLIAASHKPDAEFAILTANSST
FKTELLLTQSGVKVQPKNWILVGLQDVPGFEAVALGQLVPAEKVQPSIEAMCKDLQAKHPKLKGFLLECTELPHYADAI
RAKTGLPVWDAITMVDYFQNACSADPQLTQKALX

>K.brevis_A CAMPEP_0188921804 /NCGR_PEP_ID=Karenia-brevis-CCMP2229-20130916|56488_1
/TAXON_ID=156230 /ORGANISM="Karenia-brevis-CCMP2229" /LENGTH=600 /DNA_ID=CAMNT_0032408147
/DNA_START=61 /DNA_END=1859 /DNA_ORIENTATION=-
MGCGSSVKAQPSDGKDTTPAAPGAAPASAPASKPAAAPKAASAPATVKKAASAPVLGVVQLDTNEPTKKAFNPCHA
SMGYDIIYKEVEKLTFEMAGKGEMTEEAGKNFDEAIKYLEGKGCSAITGDCGLMMAFQNRARKVAKVPALMSSMVQC
PAIACSLEPEEKILIVTKDAETLRMQKGELLKDCGFDVETARFSIVGISDIDMSKPTKEVQPAIVKKVADTLKGDRSIRAIL
LESTQLPPYADALRAITKLPVWDCVTMVDFITSSYMDNPRFGLNDWQEKMEHTKTQDLSDPKPSKEQIMRVHENLSK
KQHPMLGCIRLDFNFNGAKVGDVQNPASFGYDVIYKAVPGLTFALAQEGKMEGQVLEDFKNSVKFLEEKGASCITGD
CGFFMAYQKQARDVANVPVFMSSMVQCPMVSAMFAKDEKILILTANSESLLPQKEVLLSSCGFNVDDDHFVIEGCQDI
PGFEAVLEGKVADVPYIEKGMMVKMKKIMEKTPNIKAIVCECHGLPQWADAMRTSTGLPVFDIVTCADFFISSMRDNP
RFGMNNWQHPWDQIEQEYQFGKHLTAAELEELRLSPDSAENIRQFTRLLLIEIIX

>K.brevis_B CAMPEP_0188855392 /NCGR_PEP_ID=Karenia-brevis-CCMP2229-20130916|7473_1
/TAXON_ID=156230 /ORGANISM="Karenia-brevis-CCMP2229" /LENGTH=992 /DNA_ID=CAMNT_0032328855
/DNA_START=1 /DNA_END=2977 /DNA_ORIENTATION=-
XQATSLGARRLAHPSTLAPAASCMTSQRRHLNLHEYQAWGIFKDFGVAVPKSTPAFSVSEVAEKSKEFAEEVVIKSQ
VMTGGRGLGHFKESGLQGGVHITATAKVAELAEKMLGNTLVTKQTGESGLPVHTVMLCERFQIKNEKYFAILMDRGS
GGPLIVGSKVGGTSIEDIAESDPSAILKMPIDIMDGITEDQALTFASMMGFSGEQQKNAAANIMGLYKVFIECDCTQIEIN
PLAELTDGRVIVCDAKVEFDDNASFRQKDIFAKRDTSQEDPLEVEAKAVDLNYIKLDGEVAIMVNGAGLAMATMDLVN
QCGGTPANFLDVGGAANRQTCKAAFKLLQTDPNVKVLFVNIFGGIMKCDVIADGVVGAVQEMGLKLPLVVRLEGTNV
EQGKKIINESGLPVYFYDDFTTAAKKAVEISKAPRLASLGIVRLDYDYPASPGDIDHPGSFSYDVFYRAVPGLTFEMCQ
QGVMPEDVKAEFLDAVKFLIDVKGVSAITGDCGFMMYFQKLARTITTHEPVFMSALVSLPAVTCAYAHDELIGIFTANG
ESLEPMHDLIKDECGVDTHEHRYIIVGCEDIPGFEAVALGTKVDYDKVEPGVVKRALETQAAHPTMRAILFECTELPQF
SDAVRHATGLPVYDAITTCNLFMEGLRDNERFGKNNWHAKWDGEQEEYKFGAELTKEQQAKLVNKPAASTRPARRH
FRSAGTQSHSFHPQSLPKTLRKMGKAAKAAPKKKLPSAPFSDKKKKVARNPLFEKAPRNYRLGGDIQPKRDLSRFVK
WPKYVRLQRMKKIMLMRLKVPPSINQFNMTIDKNQAASVLRLMKKYSPETREAKKNRLMEMAQQKKDGQEVKTKKP
QVIKYGLNHVTTLVENKVAKLVVIAHDVEPIELVCWLPALCRKKEVPYCIIKGKGRLGQLVHKKSASCVALTTVNNEDKK
ELDTLAGNFKAQFNDNAEHRRRWGGGIMGIKSQHVTARREKALEIERNKKLGLSIA

>K.brevis_C CAMPEP_0188922230 /NCGR_PEP_ID=Karenia-brevis-CCMP2229-20130916|56810_1
/TAXON_ID=156230 /ORGANISM="Karenia-brevis-CCMP2229" /LENGTH=554 /DNA_ID=CAMNT_0032408693
/DNA_START=76 /DNA_END=1740 /DNA_ORIENTATION=+
MGCCCSSSKPKAKTQQKGKVKAPMLGVVKLDSKYPGSKQCAASMGYDVMYKEVPKLTFDMAVKGEMTAEVGKNF
DDAIKFLQSQGCSAITGDCGFMMAFQKRARQVASVPAFMSSMVQCPAISCALEPAEKIMIMTASAPDLQKQKTQLLK
QCGFDVESSRYIIVGLEGIPAFNGTSKDAPVSEVQATVVKVAAESIKKNPTIRAILLECTELPPYADALRAVTGLPVFDCI
TAADFFKSAFIDDPRFGLNDWQETIKTATKKPNDTQLKKVADNLSKRQNPILGCIRLDFNFDGAKVGDVQNAGSFGYQ
VIYKAVPGLTFELAQKGTMTPQVEQDFKNSIKFLESKGATCITGDCGFFMAYQKIARDAANVPVFMSSMVQCPIVSTAF
DMEEKILILTANSESLTPQKSVLLSSCGFNVDDGHFVIEGCQDIPGFEAVLEGKVADVPKIEKGMMTKIKGVLQKNPSIK
AIVCECHGLPQWADVMRKASGLPVFDIVTCADFFISSVADNPRFGLNEWQAPWDKQMEEYKFGQGLTAEEKERLSK
ESQDRMKNK

>C.cohnii_A CAMPEP_0193854358 /NCGR_PEP_ID=Crypthecodinium-cohnii-Seligo-20130904|11054_1
/TAXON_ID=2866 /ORGANISM="Crypthecodinium-cohnii-Seligo" /LENGTH=411 /DNA_ID=CAMNT_0038406789
/DNA_START=1 /DNA_END=1236 /DNA_ORIENTATION=+
RTMCLQHQNEEFSWTSGNTATTTTPSMQMAPPSSSEVSLAAAAKMRFANGGGGGPPAKPSSSSRGLVSKVVESPT
RQDSKSTTPNSAKSSSTASTASTRSGTTKASAEGSSRGFNCPLKERASFLPNVGEAPQAKREVVKQVILGVMRLDYE

YPPALGDIAHPGSFSYKVVYRVVHGLTFEMAQEGVLTPEVEREFVKAIRDLEMRDVNCITGDCGFMMHFQELARRHA
RKPVLMSALLQLPLITCSLAKKERIAIFTANGQSFSSMHQLLKDEFGVDLDSRCVFVGCEDVEGLEAVALAQKVDVATV
TPGIVDRACAVSKEYPEVRAFLFECTELPPYSDAVRKATGMPVFDALTCCNAFIAGFKDNAFRFECNAEQQRSASKQ
QQQQLQHQQQLQKRSKGVPRTGATKA

>C.cohnii_B CAMPEP_0193926808 /NCGR_PEP_ID=Crypthecodinium-cohnii-Seligo-20130904|174461_1
/TAXON_ID=2866 /ORGANISM="Crypthecodinium-cohnii-Seligo" /LENGTH=666 /DNA_ID=CAMNT_0038517207
/DNA_START=1 /DNA_END=1996 /DNA_ORIENTATION=+
XGARVLASLTPIVLFLEQSFAGLGGKAFRPMSGFHNNGGFSSWASPGAGNSMAATPRGTFGASGALAATARMQFGA
GMAPIAPRITSNGVMHDLESTFKTWVNGSTQGRAQLWKAWHSIDYNGNGRVSLAEIDKWVVEQFVQANNKPALMRA
YKASIINEKDGYVHKHDFPVLLRNVVYFNKLWSVFGGIDTDGDRRLTYDEFTKGLALLGLGGYGKDAHNIFNQLDTNR
GGIILFDEFCKWVASVQCPVDAKVYDPTTMQSPMKATPRAPASNLGQAGRSLMTPRGVPVARQILTPTGPGGVRAS
KAGAGGIESLLKDRPALKPDVNEGIKVLKQKKGVLSKHASLGVLRLDYDYPPAPGDIDHPDSFGYDVFYRVVPGLTFE
MAQSGKMTPAVEREFVEAIRFMEAKDVSGITGDCGFMMYFQELARRHTKKPVFMSSLAQLPAVTCSFAYNEHIAVFT
ANGKSLAPMRDLIKEECGVDPEESRYIFVGCEDVPGFEAVALGEKVDVPAVTPGIVAKALAVLQQFPTIRAFLFECTEM
PPYSDAVRKATGIPVFDSITCCDFFISGFRDNARFGINDWQEDWDGVQDGYVYGQNLSAAAESNAGEQGPLASSSS
HVNSWHPSDHSSPQQERFTQETCLPRHPGHRDVLNGTDLYLLQAGNVASR

>C.cohnii_C CAMPEP_0193865556 /NCGR_PEP_ID=Crypthecodinium-cohnii-Seligo-20130904|19770_1
/TAXON_ID=2866 /ORGANISM="Crypthecodinium-cohnii-Seligo" /LENGTH=749 /DNA_ID=CAMNT_0038422267
/DNA_START=1 /DNA_END=2249 /DNA_ORIENTATION=-
XSPALHSSGTMFGFHPFGHHHEAVHSRLDLLEAEVTQMHLQKHMGQGMPSHQHPNQAFLTFHGPMGSPGSLGSTV
STVALPTRCASPAATPAPSMMTTTLPGSMPSPSNAGPMVPAEHYHQIMQLYSEAVGHLAKLTTKHNDLLVNHQQLM
AAHYDVLTAMAQDGASGASGGKGEAASGKSKAPKMGVIRLDYDYPPAAGDTDSPASYGFSVIYRVVPGLTFEMAQA
GVFTEAVERRFAEAIKYLEHKGVSSITGDCGFMMAFQVLARKIATVPVFMSSMVQCPVIGAAYDHKEQILILTANGTSL
KPQKEVLLKSCGFHVDSERYVIHGCQDVPGFDAVAKGEAVPVELVQKGIIALTRKILQENPRIQGILLECTELPPYADAL
RATTGLAVWDAVTAADFYMSGFKDNPRFGVNDWQEEFNGEMAQYTFGQNLIEKDAALLVNKREVGTDVKADAVVK
QRAIKAHAKAKAARKVRKLVKQQAVCLGVLRLDYNYPPAAGDTDCPGSYNYDVVFRMVPGLTFEMAQSGMMTYKV
QQEFVKAIKWLEAKGVSGITGDCGFMMAFQPLARDVAKVPIFMSSMLQSPMLSVAFDKYDIILILTANSQTLQPQKQIL
LKECGFDVDSERFIIQGCQDVPGFDAVAKGEKVDVEKVTPGIVKLVMDLLSMQPNIRGILLECTELPPYADALRMTTGL
PVWDAVTCADFFISSRKDNPRFGLNQWQNDWDGTHDDYKLGDNLTAEEKSHALNI

>C.cohnii_D CAMPEP_0193933962 /NCGR_PEP_ID=Crypthecodinium-cohnii-Seligo-20130904|195906_1
/TAXON_ID=2866 /ORGANISM="Crypthecodinium-cohnii-Seligo" /LENGTH=820 /DNA_ID=CAMNT_0038526745
/DNA_START=1 /DNA_END=2458 /DNA_ORIENTATION=+
XGPNLPRAAWGLQRVSKDRSAWADSGTQPSNVEVNRCIEALAVAPCDSAGPSHLAMLPPLRAPGIPIAMMGNPSPH
QVAIPVSTAWGTPRLHAGPHYGHPFGGTVHDRLDVLEHELCCMKVEMGTSKGPGSSSPMGDPLSARSLNSPSEALP
YGSPRQTMSGLLPEMSPGKDSLATVTTAASPVANTVSVEVHQDLLKQHADLIRQMAVLQQRHSALLDTHIDLMSAHA
EVLEYTSQLQKGSTAAGKAEEDTKKKKSQASHLSKHPSIGVIRLDYNYPPAEGDTDSPGSFGYDVTFRVIPGLTFEMA
QQGKFTDAVERRFAEGIKFLEHKGVSAITGDCGFMMAFQVLARKIATKPVFMSSMVQCPIIAAAFDPKEKILVLTANGL
SLKPQKEVLLSSCGFDVEDDRFLIYGCQDLPGFDAVEKGLAVPIDVVQPAIVKMTMDILQTHKSRKNKIAGILLECTELP
PYADALRASTGLPVWDCITAADFYVSGFKDNPRYGINDWQAEWDQEQAEYHFGENLVEKDREQILNQEQAKPKAKA
KAKAKSKAHAYKLKKLVKQQAPTLGVLRLDYNYPPAAGDIDCPGSYEYEVLFRCVPGLTFEMAQAGMMTYAVQQRF
VAAIKWLEAKGVCGITGDCGFMMAFQPLARDVASVPVFMSAMMQSPMVSVSFDKYDKILILTANSDTLKPQKETLLSH
CGFDVDDDRFVIEGCQDVQGFDAVAKGQKVHVEAVTPGIVELAQRHIDKCPDIRAIILECTELPPYADALRMATGLPV
WDAITCADFFISARKDNPRFGLNQWQNDWDGVVEEYKLGNNLSDLX

>C.cohnii_E CAMPEP_0193866376 /NCGR_PEP_ID=Crypthecodinium-cohnii-Seligo-20130904|20414_1
/TAXON_ID=2866 /ORGANISM="Crypthecodinium-cohnii-Seligo" /LENGTH=756 /DNA_ID=CAMNT_0038423447
/DNA_START=1 /DNA_END=2271 /DNA_ORIENTATION=+
EREASFTSHLWPSHPSSKREQQQQEQTIQKQQIQSKTNTNVANMHNGAMMQHRLDLLESELMHLKMGGMSGPMM
PMPGQGMVSMALPIPSHPSMGGGSPFGLAATPSTVAPSPGGSPGTQMAPVGGQNDLILSNHKEIMEAHKLLAATME
AQQAKHDLLLESHGKLMEAHKKLMASKVGLGKDAKKEGKKHPILGVVRLDYNYPPAAGDTDSPASFGYDVVFRVVP
GFTFEMAQAGKFTEAVERNFAEAIKFLELKGANAITGDCGFMMAFQVLARKIATRPIFMSSMVQCPVIACCYDPKDQIL
ILTANGLSLKPQKEVLLNSCGFDVDEDRFIIRGCQDVPGFDAVAKGEAVPVEVVQPGIVKLTMEILQQNPKIAGILLECT
ELPPYADALRAATGLSVWDAVTAADFYINAYKDNPRFGINDWQAEWDGEMTDYTYGDNLIANDKQELVNKIGNAVPA
KTLAMPKSKAKAKSKALMQKIKRNAQKHAAPCLGVIRLDYNYPPAAGDIDHPGSYDYDVIFRAVPGLTFEMAQSGKMT
YTVQKEFEAAVKWLEAKGVCGITGDCGFMMAFQPLASDIAKVPVFMSAMVQCPMISVAFDKYDKVMILTANSKTLKP
QKQVLLSQCGFDVDDDRFIIYGCQDVPGFDAVAKGEKVDVEKVTPGIVELARKILDQEPTIRAICLECTELPPYADALRA
EFDLPVFDAITCADFFVSARKDNPRFGLNQWQNDWDGTFEEYTLGQNLTDVQKSALLNA

>A.spinosum_A CAMPEP_0186837182 /NCGR_PEP_ID=Azadinium-spinosum-3D9-20130829|183522_1 /TAXON_ID=632150 /ORGANISM="Azadinium-spinosum-3D9" /LENGTH=421 /DNA_ID=CAMNT_0029731213 /DNA_START=1 /DNA_END=1264 /DNA_ORIENTATION=+
XLKARQVLSRPARFVQLAIPCIADRSPGIMEPMYGGGMGSSSMMMQNGMMNRNSMDVRLSQIEHEVMLHKAQNAQ
SAMMNRSGSAPFLPPVGSQGLPTKEHQELLQSHNKLLGEYSDLMTRHSKVLDKQRELIAKQELALYTPVLGMDKKNT
DLRKAASLGIIRLDYDYPPAPGDIDCPDSYDYDVHYRVVPGLTFDMCQSGKMTRDVEEEFIEAIRYLEKRGVQGVTGD
CGFMMYFQALARQHTKKPVFMSSLAQLPAVTCGFAKDELIAIFTANSTTLTPMKDLIKDECGVDAHTKRFVIVGCQDV
PGFDAVEAGGKVDTAKVTPGMVKKAKDTLKKYPHLRAYLFECTELPPYSDAVRAATGLPVYDSITACNFFMTGLQDN
KRFGLNNWQEAWDKQQDDYSFGGNLTAAQRAKLVNK

>A.spinosum_B CAMPEP_0186768496 /NCGR_PEP_ID=Azadinium-spinosum-3D9-20130829|29318_1 /TAXON_ID=632150 /ORGANISM="Azadinium-spinosum-3D9" /LENGTH=325 /DNA_ID=CAMNT_0029649709 /DNA_START=1 /DNA_END=979 /DNA_ORIENTATION=-
ALLPGSLALEANEEKAGFNEQKAGFLPGEQLSEEVEHRDMGAAAAAELPPGKSLGIDHAVLGVIRLDWHYHPMPGDV
GSPSSFEFPVVYRAVPGLTFEVCQSGQLTPEIEAEFIKAIQWLDRIKGVSVISADCGFFMWFQKLARRHTSKPVVMSS
LAILPAINCAIHGKVGIFTANSESLAPMHDCVKDECGVEWNESRYVLVGCQDVPGFEAVAHGTEVDFQKVMPGVVGK
AKQVLADHPDISAFLFECTQLPPFSDDVRAATGLPVYDAITCCDSVMMGFVDNPRFGLNTWHKSFDGEREQYKLGDC
LGPEGKRRLVNLPANG

>A.spinosum_C CAMPEP_0186753562 /NCGR_PEP_ID=Azadinium-spinosum-3D9-20130829|17069_1 /TAXON_ID=632150 /ORGANISM="Azadinium-spinosum-3D9" /LENGTH=784 /DNA_ID=CAMNT_0029630991 /DNA_START=1 /DNA_END=2353 /DNA_ORIENTATION=+
XAQRSESPDVRLARLNRLESELRDLRERTPDVQLARMTRLEHELRDLRESHQHKIQVDAARASLAGSLTPTDLHLSW
MVSQQSQTPGAGSCPPSGWQLSCAQVADQSQQKSGWSPSADSNSQVQTQKGMLQASRELLAALQVQHKSHAEM
LESHMQFAEAQSDNLHLAMSLIGASHRMAQAQPQAQAQAQAQADAKAQGEAKAQAQAQAPGECLTQPLTHGEGKD
PDAVRMRLQEHAGGLGIVLEGKRRRPLLGVLRLDHDYPPASGDINSAASFGYEVVYHSVPGMTLSMARSGSFTEQA
EHNFAEGVRTLEQRGASAITGDSGFMMAFQGNARRIASRPIFMSALVQCPPIMASLETGDKIMVLTSNAERLRLGREIL
LSNCGVDLDESRFVIEGCEHVPKFDAFEKGEATPFSIVQQGIVRLTLRTLKKHPAVKIILLESTQLPPYADALRAATNIAV
FDCITAANFYISAFQDNPRAGLNSWQEKKADGGDESKAEEAAMAKATEKVVKAKKGGKSLKKRMNPILGILRFDRSG
SAIATGSAGSEELEIRRPCAYGYEVISRTVPGLTQEMAMAGHLGDEDEKIRHAFAEAIRWLEAQGACCITGDCGFLLTL
QAAAREAAEVPVFMSPLAQWPLLAAAFDPSGEQVLVLTAAERGLQIQRESLKQLCGVDLGDERLLLCACEGVPSFSA
LLLGGPVDLDHVSAGLVRLVVEKLERAPKVRALCLECVELPPFSDALRAATGLPVFDAVTCADFFVSSRADNPRAGLN
NWQEDWDGMDSD

>A.spinosum_D CAMPEP_0186759428 /NCGR_PEP_ID=Azadinium-spinosum-3D9-20130829|21942_1 /TAXON_ID=632150 /ORGANISM="Azadinium-spinosum-3D9" /LENGTH=620 /DNA_ID=CAMNT_0029638239 /DNA_START=1 /DNA_END=1858 /DNA_ORIENTATION=+
XSSFSFFARTAVSDNSFPALQHWWAMAKYGTHPELAKHEQKFGILCFESAGATDKTLSLGYGYSAIQKVVKGLTPAT
AKEPNPKVLQALKRAVQELEAAKVEGITGESCLVLPLQAEVRSMSRVPIFLSALLQAALLIAAHDKQQKFLIVTKHADEV
NARKEELLKMCGVMVAPERFMVLDLLGRSPDALAKELGTMKDATPELAGVIFDCIHIVQHADQLRSITKFPVFDQVTL
MDFYFCARSDNPYFGTCYKEVTGNMKKGEVSHLVKKAQRRLFSPYENYQANLVSKTDASVAVGVLRIDYSYPPIPGD
VDHEGSYGYRVHFEQVDGLTFEKAQSAVLDEAIQESFKKAVKKLEEKGVVAITGDCGFMMAYQPYVRALTEIPVVLSS
ILQAPMLVATHEPEAKFAIFTANSSTFDKEKLLAQSGMTVDTSQWIVVGLQDLDGFEAVALGQKVPAEKVQPGILKCVK
NLITKEPQLRGILLECTELPHYADAIRADSGLPVVDAITLVDFFQSSCTTAPAFSKKTLEAFLGVLDPRLAISTVAVVTAG
TLGAAVGVTGKVAQTASNTASVVAYCSESRESYVEPPGESPIALGIMYQASTAGEASDEESHGCQAVWRX

>S. Kawaguti A jgi|Fugka2468_1|26563|SymbF.scaffold994.5.m1
MPVLMLGGLGVATGYAFALWVKGQEPPRGKDPNIKHASLGVIRLDWHYHPLPGDVGSSDSFEYPVYYRAV
PGLTFEVCQSGKMTPEIREHFKEAIRYLDQDKNCSVITSDCGFFMWFQKEARQYTAKPVVMSSLAILPAV
HGALGPDGKIAILSANSESLAPMHDTVAREMGVDWNADCYVLVGCQDVEGFDAVATGEVVDLAKVTPGIV
KKAVEVTKKHNIHAILMECTQLPPFSDDVRAATGLPVYDAIVCADFFVRGFVDNPRFGLNDWHQRWDGKQ
EAYRFGDVEDKAKLIHYSPTNN*

>S. Kawaguti B2 jgi|Fugka2468_1|14610|SymbF.scaffold3123.3.m1
MAGNVGALLPSIGRAEVPADDKTIGSATAPSVTMPLLPGLQQSAEGPAAPMTAPPKKPEQPAAQRLSSEL
TKRRLQLSENKEQKHPGNASNDQSPQDLQVDSKVSARTISTSTASEIASPDKSQDVKGNPEDDDAASEGN
ESEVLSLGEEEKDEFGDFNQARRTEDERDDWRAAGDGRRASARKSVALTLSQKLFKAAKKSKEALTKDAF
AVLCFDEGDADDEVVEAKDRGCRRFFSICKGVTKEKLMSDDLDSAAVESIKACFDRIVKKAGQESIAGVS
CDVGYLWMKHQRTLRQAFPDFTVLTTPMMQLPFIWTCFGTNDKTLLVTWKDETDDSKAHHLHMLEDAGIR
LDTHRVEILTLNASDPKWSLFLAGMTTQKENQKLVNEVVKMVHQQIIQLFDQDIDVNSIMLDSVLSKFSE
VLGKATDLPVFDEASMLKLFSSASSLSHFSDASVLCRLQEKKSKAERKSYEADKGTVGLIQLEYEYVAAV
GDIDHGSTFQFNSCPEVVPALTFEEAQRGSQNPVILENLKLAVDKMEREECFGITGNCGFMHFYQQFVRD

YATVPVFMSALVQVPAMAAALEPDERLLVLTANESSFMESRDALLSAEGRPFCDFNRVLVRGCENVPGFE
AVANAEQVDTIKVQQHLQKYVQEIKHSDLGPIKAILLECTQMPHYAAAIRQSTELPTFDVVTCANFFAQV
LNKDQLYQKEAPEEINCTKAKDSSATRVWDKVSCL*

Bacillariophyceae

>S.grethae_A HBLG01002042.1 Skeletonema grethae MMETSP0578-249982-11834
EQTQYQNEFVAFMGELDMGQNFDYVFPKIETEGGPFQIIPNSVYCVKPDTVMAEAQKHTDDSAPFAVGPYTLGDADV
TLVSTRKVFKVPFSLIGQFLAIPVGADVAKYFWAVIYPVIKAAGNGKIKECKSLIDYFLIAVTIREEGEVMQQVRLSEVQL
PRPVERDIVLKHYRLRELGRLFPKLFPVPVDVPAMDGAEVLDDHHYGQNNDHNRHQTESISTQRTRRMSVDLGVQG
TLYLSYKNDITQKNMSTPKLGILRLDYNYPPAVGDIDNAESFDYPVIYRVIPGLTFEMCQSGELPDNVKAECLRAVKYLE
QQGVSGITGDCGFMINIQDLISDATNKPVFTTSLVQLPSLCMCFDSDETIAVFTANSNTLLPILPRMLEMCALRQEDKD
RIQIVGCQDVDGFDAVEKGEAVDVVKVDAGIVALAKSVIAANPRIKAILLECTELPAYADSLKLETGLPVYDSISVCNSF
MAGFLDNPRIGLNDFHEAWDGTQAEYSFGACLSRASQAKVVSRISSLFPLSTFEEIDDIDIDTDDEE

>S.grethae_B HBLG01015097.1 Skeletonema grethae, MMETSP0578-23584
MIYTQNSNSTLNLTNFSYIYVDHPESFDYPVVYRVIPGLTFEMCQAGDLTEEVKRECRHAVQYLERQGVSGITGDCGF
MIHIQDLISEVTNKPVFMTSLIQLPSLCMCFDQDETIAIFTANSNSLNPVLPQLLEMCALRQEDKDRIQVVGCQDVDGFD
AVEKGVEVDVLKVDAGIVALAKRVIEENPRIKAILLECTELPAYADSLKFATGLPVYDSISVCNSFMAGFLDNPKIGLNGF
HEEWDGTQKEYTFGACLSENELTKLCSQTTILG

>S.marinoi_A HBJE01014325.1 Skeletonema marinoi, TRINITY-DN1794-c0-g1-i1
LRFGCSSKKKTKDGEAGIDDVKTTGHTLDNSATTTNSSEGDVAGTARAEVHVQHYGEKNCRPQPPSMQRQSSTLSV
QDRICLTYQKAISEKNFSTPKLGILRLDYNYPPAVGDIDSPESFHYPVIYRVIPGLTFEMCQSGELSEEIKVECRQAVQY
LEKQGVSGITGDCGFMIHIQDLISDVTNKPVFMTSLVQLPSLCMCFDSDETIAIFTANSTTLLPVLPRLLEMCALRQEDK
DRIQVVGCQDVDGFDAVEKGEAVDVLKVDAGIVALAKRVVAENPRIKAILLECTELPAYADSLKFATGLPVYDSISVCN
SFMAGFLDNPKIGLNHFYEAWDGKQAEYTFGACLSENEQGKLKSAKAG

>S.marinoi_B
MGLKSAFPWWALRFGCSSKKKAEDGKAGIDGMKTTGHTLDNSATNSSEGDVAGTARAEVHDQHYVENNCRPQSPS
MQRQSSTLSVQDSICLTYKKAISEKNFSTPKLGILRLDYNYPPAVGDIDSPESFDYPVIYRVIPGLTFEMCQSGELSDEV
KVECRQAVQYLEKQGVSGITGDCGFMIHIQAMISDVTNKPVFMTSLVQLPSLCMCFDSDETIAIFTANSTTLLPILPRLL
EMCALRQEDKARIQVVGCQDVDGFDAVEKGEAVDVLKVDAGIVALAKRVVAENPRIKAILLECTELPAYADSLKFATGL
PVYDSISVCNSFMAGFLDNPKIGLNDFHEAWDGKQAEYTFGACLSENEQGKLKSGRNLTSQISSLSLPNFDDNDISEE
DSVSHAMTL

>S.japonicum HBLJ01001483.1 Skeletonema japonicum MMETSP0593-11330
HAAACSRYGERLIAFKTTVPVPKLGILRLDYNYPPAVGDIDHPESFDYPVIYRVIPGLTFEMCQCQTGDLTEEVKIECPA
VQYLERQGVSGITGDCGFMIHIQNLISDVTNKPVFMTSLVQLPSLCMCFDSDETIAVFTANSDSLSPLLPRLLEMCALR
EKDKERIQLVGCQDVDGFEAVEKGEQVDVLKVEAGIVALAKRVIEENPRIKAILLECTELPAYADSLKFATGLPVYDSIG
VCNSFMAGFLDNPKIGLDDFQDSWDGIQEEYSFGACLTKKERAKLVSKIRRSVRLSIIVNKLRSAVSAE

>S.robusta jgi|Semro1|16691|Sro2472_g328650.1
MPRRASVVAHHQHSLSLAQTIRSAIHDGAKLFNDGKKKECYELYLATAYGGLSQQTMHAMHAHNRVEGAVETMGHQ
HDKYTDAQSEKIRELLTKATVDANVVASKEDYGEAAWVIRRAFDLILKMTKRASKASELKPAKTGGETQDDVSFITISD
GATEREQAETLSGLCNKMADILDIGNHKYRLTTYPDTFVGKEAVTKLVKTNLCTSREDAVKKLNKLMKYGLIYHVTKEH
KFKDEPLFYKLTASTDLRFELDKFGTKTNTLEGEELVHYATLLGRFKSFHKHPSLGVLRLDYDYPPAAGDIDHPDSYDY
PVYYRVVPGLTFEMCQSGVLSAEVGAAFDDAVLWLAIHKDVSVISGDCGFMFWFVERCRKIASKKIISLSPLMQLPIMV
AACAPKDKILVLTANGKSLDPMHDLIKRQCGIDGQNTQFIIVGCENVPHFGVEVAKGLQVDVDKACPGIVELAKEMVR
DHPDTRMILMECTELPPYSDAVREATRLPVWDAITNCNFFMQGFLDSVNFGLNGWYEEWDGIQEEYVLGQNLNDTQ
RLQCVWCRNNELNYAGK

>Pseudo-nitzschia multi jgi|Psemu1|327799|estExt_fgenesh1_pg.C_8290001
MRVQPLRQSTLSPTLSDPKHSFDVRVATIATEANGKKEEKSDSFNPKIAVLRLDYFYPVRKGDVDSPDSFDFPMVYET
VEGLTFACSQKGLDNNSVLSDSKIAKSDADAWNARYGQERKYEYSLEREREANDQTYLCYKSETIERSLRETIRRLLQ
REGKNIIGITGNCGFFMSLQDKVEDLSSQVAEELNIPAPDVMMSSLLQVAFIQQTIGRRNKLAIVTANSKSFLKEFKALL
PFGARASNIVVVGMEDVEGFGEEVEKGWAVQPCAGDNIVEKLRMALNQDTSIKGILMECTELSFVSDRVRAEFEMVV
VDSLTMLNYFYRSRTGGFLA

>P.fradulenta_A CAMPEP_0199812070 /NCGR_PEP_ID=Pseudo_nitzschia-fradulenta-WWA7-20140214|96008_1
/ASSEMBLY_ACC=CAM_ASM_001165 /TAXON_ID=183588 /ORGANISM="Pseudo-nitzschia fraudulenta, Strain
WWA7" /LENGTH=421 /DNA_ID=CAMNT_0045699945 /DNA_START=1 /DNA_END=1264 /DNA_ORIENTATION=-

XLKARQVLSRPARFVQLAIPCIADRSPGIMEPMYGGGMGSSSMMMQNGMMNRNSMDVRLSQIEHEVMLHKAQNAQ
SAMMNRSGSAPFLPPVGSQGLPTKEHQELLQSHNKLLGEYSDLMTRHSKVLDKQRELIAKQELALYTPVLGMDKKNT
DLRKAASLGIIRLDYDYPPAPGDIDCPDSYDYDVHYRVVPGLTFDMCQSGKMTRDVEEEFIEAIRYLEKRGVQGVTGD
CGFMMYFQALARQHTKKPVFMSSLAQLPAVTCGFAKDELIAIFTANSTTLTPMKDLIKDECGVDAHTKRFVIVGCQDV
PGFDAVEAGGKVDTAKVTPGMVKKAKDTLKKYPHLRAYLFECTELPPYSDAVRAATGLPVYDSITACNFFMTGLQDN
KRFGLNNWQEAWDKQQDDYSFGGNLTAAQRAKLVNK

>P.fradulenta_B CAMPEP_0199831756 /NCGR_PEP_ID=Pseudo_nitzschia-fradulenta-WWA7-20140214|209615_1
/ASSEMBLY_ACC=CAM_ASM_001165 /TAXON_ID=183588 /ORGANISM="Pseudo-nitzschia fraudulenta, Strain
WWA7" /LENGTH=497 /DNA_ID=CAMNT_0045724545 /DNA_START=1 /DNA_END=1488 /DNA_ORIENTATION=-
XAKEPNPKVLQALKRAVQELEAAKVEGITGESCLVLPLQAEVRSMSRVPIFLSALLQAALLIAAHDKQQKFLIVTKHADE
VNARKEELLKMCGVMVAPERFMVLDLLGRSPDALAKEXXXXXXXXXXXXXXXSSSSXADQLRSITKFPVFDQVTLMD
FYFCARSDNPYFGTCYKEVTGNMKKGEVSHLVKKAQRRLFSPYENYQANLVSKTDASVAVGVLRIDYSYPPIPGDVD
HEGSYGYRVHFEQVDGLTFEKAQSAVLDEAIQESFKKAVKKLEEKGVVAITGDCGFMMAYQPYVRALTEIPVVLSSIL
QAPMLVATHEPEAKFAIFTANSSTFDKEKLLAQSGMTVDTSQWIVVGLQDLDGFEAVALGQKVPAEKVQPGILKCVKN
LITKEPQLRGILLECTELPHYADAIRADSGLPVVDAITLVDFFQSSCTTAPAFSKKTLEAFLGVLDPRLAISTVAVVTAGTL
GAAVGVTGKVAQTASNTASACCVLQ

>Attheya_sp.
MVRVARKEVPSLEISIRKALIKGATVYNEGKQQECFEIYLATAYSGLSQQTMHGHGYHDDYTDEQSKTIIQVLIQATIDG
NAQASASSKDFGEASWVLRDAFDAILKMSKKKTDDEDGQDHDKDEGGILANLCNTLPDILDIKEHRHHLQTYPDTFVG
DEAVSKLVESHVCMNRQHAVEQMNKLLRFGMLYHVTKEHTFEDERLFYQLASSSDLKLELNKFGNKTKSLRGDDLVH
YTAILRRYITFHKHPSLGVLRLDYDYPPAPGDIDHPDSYEYPVYYRVVPGLTFEMCQEGVLTPEVAAAFDEAVLWLAN
HKDVSVITGDCGFMFWFVERVRKIASKKCISLSPLMQLPGMAAACGPKDKILVLTANGTSLNPMRDLIKRQCGVDSQD
TKFIFVGCENVPYFGEEVANGTKVNEEKAEPGIVQLTQEMVRTYPDTRMILFECTELPPYSDAVREATRLPVMDAITNC
DFFMQAFLDNGKFGLNGWYEEWDGVQDEYVMGQNLDEGQRLQCIWCAKTDVHQFVT

>Ta_antarctica CAMPEP_0200113894 /NCGR_PEP_ID=Thalassiosira-antarctica-CCMP982-20140214|20162_1
/ASSEMBLY_ACC=CAM_ASM_001185 /TAXON_ID=420261 /ORGANISM="Thalassiosira antarctica, Strain
CCMP982" /LENGTH=286 /DNA_ID=CAMNT_0046052863 /DNA_START=1 /DNA_END=859
/DNA_ORIENTATION=+
XKDKAIKGAASARARAKKANASLGVVRLDYDYPPAPGDIDSPDSFGYDVFYRAVPGLSFGMCQRGELTDKVKAEFIE
AVQWLDKEKGVSAITGDCGFMMWFQDLARQHTKKPVLMSSLVQMPSIAAAYSKTEDIAIFTANSVTLTPMNSLIEEQC
GVNPEDTRFHIVGCQDVDGFEAVAVGGKVDVPKVTPGIVALAKAELAKNPSIRAILLECTELPPYADALRAATGLPVFD
AITCCNMFIEGMVDNPRFGINDWQEGWDGVQEKYELGQNLDAEDKKALESAV

>Tx_antarctica CAMPEP_0200964928 /NCGR_PEP_ID=Thalassiothrix-antarctica-L6_D1-20140214|17297_1
/ASSEMBLY_ACC=CAM_ASM_001238 /TAXON_ID=1049557 /ORGANISM="Thalassiothrix antarctica, Strain L6-
D1" /LENGTH=453 /DNA_ID=CAMNT_0047172213 /DNA_START=19 /DNA_END=1380 /DNA_ORIENTATION=+
MSEGGSNSGETPVDVVETTKEEEVSKTEEETVKTKEEAVKTKDEIVKTKDEIVKTEDEAAQTEEEVAKTEEEVVKTEE
EAAQTEEEIVKTEEVVEGKEEEEIVKTEEVGAKEEEEEVVEEVVEKEEDAEQEETIKEENTTNKNEISTDIVASKEKEEM
IDYFSATPPSLGILRLDHDYPPALGDINNAETFTYNVYYRVVPGLTFEMCKSGKLSKDVQEKLFEAVDYLEKEKDVSGI
TADCGFMMFFQSLVRSRTKKPVFLSPLTQLPAVTSAYSENEKIIIMTSNGKSLEPMRDLIRKECGVDTQQERCIIVGCE
DVDGFEAVGEKVDVAKCTPGIIQKAMDILQEHPTARAIVLESTELPPFADALRFYTRIPVYDAITNANFFMTGLQDNERF
GMNNMEKWDGKQKENYTFGQNLAEDEKENPTEEDTGKLANKSRVVPIEVEDEDDVWC

>F.kerguelensisL2C3 CAMPEP_0199381286 /NCGR_PEP_ID=Fragilariopsis-kerguelensis-L2_C3-
20140214|35188_1 /ASSEMBLY_ACC=CAM_ASM_001141 /TAXON_ID=186038 /ORGANISM="Fragilariopsis
kerguelensis, Strain L2-C3" /LENGTH=492 /DNA_ID=CAMNT_0045159857 /DNA_START=1 /DNA_END=1478
/DNA_ORIENTATION=-
XTPFGLSLFPGDTTNQVKAEPTSIRSLLRKKGLVVFAAVSLGVAGGAAGNQMYASYSNTIVEASVMASKNQFGDIPKG
TELPFDNFLGIYGPAFKSGVTSDLDATCSSNVGCLEAGLLGYQCQFFDSCFATEPPTQTKAQQAEVKEAITEYMASEL
EAQAKNLHDDTHGDESAIKNSINFAENLNLKSKYGILRVDYAYRPTRGDPGSPLSFSSNDSVSFKVEGWSFTAAQEGL
AADGTYPVGEYNMSDDSRNRYWKTPPVVNGTAVKRSGYHTTIINNDVIHRYAPDVMKKNMKEAIEYLESQNVSGITA
DVGFSQAFQESIASMASVPVVASSLQQLSFVAPMFNLNDPSKKKKILVTTSNSVNFDKDKLIPKGINHTSVVVLGMEEN
SFGKWVGIGNSFSRFKSDAFDEASVEEALAAVVEACRTTIEEEKKNGTEIVAIVQECSELPAYSNALRHEFDLPVYDFG
TAIEFVRMGRKFGNYGSYMVPSK

Phaeophyceae

>C.okamuranus_A

MQRGGKTVYGAGIGVLMLETRFPRIPGDIGHAESFPFPLQYRVVRGATPDRAVRQDPRALVDDFIKAGRDLVDMGCD
GITTTCGYLSLIQDQIKDALGVPVAASSLMQVPMVQALLPMGQKVGILTVSQAALTDSHLAAAGVPLDTPIVGTEGGRA
FYDGFLNNRVEIDITACRADLLDAANTLCRDHPEVGAIVLECTNMVPFAHDIRRDTGLPVYSIESFLTWFQAGLMPRRF
PIELADPRWR

>C.okamuranus_B
MIATGGKTVYGASVGILMLEARFPRIPGDMGNARTWPFPVMYKVVRGASPDRVVRQGAEGLVDAFIAGARELVADGV
DGITTNCGFLSLVQEELSRAVPVPVVTSSLQQVAMVNRLMPAGKRAGILTISGSTLTQAHLEAAGVPDGTPIGTTEGLQ
EFTRAILDNELTLNVEAARQDNVEAALALKKNHPDLGGIVLECTNMCPYAADITEATGLPVWSMASFVEWFQAGLAPR
HYQV

Pelagophyceae

>Pelagophyceae_sp.A
MGVGGVVDKGRARRPARDALCREAQIWFGDDDYGHEAAPARRKAQSIFPGEEPLRGRQELEAVCCPESPLTTQRK
RAQSSGVSAQEDRSPRHPSLGVLRLDYDYPAAPGDIDHPASYGYPVHYRVVPGMSFELCQSGVLPDDVKRAFIKAIR
WLDDAKGISAITSDCGFMMWFQQLARLHTKKPVFLSSLVAIPTVALTLSRRRKIAVLTANSVSLARMTDLVRRDCGGC
TFGDRLVLVGCQDVDGFEAVALGEKVEMQRVGPGVVRLALESCAQHDIGAFIFECTELPPFSEAVRRATGLPVFDAIS
CCDFFMSGYTRHERGTAGPDWFAAWDGMQEKYELGRELGLQEKARLNTTELGRRRAIANANAARAAVAATSLVAA

>Pelagophyceae_sp.B jgi|Pelago2097_1|111540|CE111539_15685
MHPVYAFVFLAGRTVAFTALRTRLGGGRAGACTSRPGRGAVGCASSKAAVEESRFPHPLTRSALGVLRLDYDYPPA
PGDVDSPLSWDYRVYYRCVPGYTFDMCKSNKLSQDVKVELEAATKWLEAQGCSGITGDCGFMMQIQPLIRNFTKLP
VFMSSLVQLPTVRAGFSNRERIAIFSANGGDLHHMFETFRSECNLDSKDEAFVIVGCEKVDGFDAVELGGKVNVTKV
QPGIVAKAVEIVKAHPDLRAILMECTELPAYSDAVRNATGLPVYDAITCCNMQSPRRPAPQRGRGAEGTPKGEKPHC
SRFMAGVQNNPRFGLEEWQRKFDGEQEAYVFGQHLTPAQRANLVNKPALVIEPASDALVIDVNKPASDALFDKPAKV
TAAAA

>Pelagophyceae_sp.C jgi|Pelago2097_1|552301|fgenesh1_kg.1075_#_4_#_Locus30470v2rpkm0.28
MGCASSNAPKVAGLPPGRFPTPLAASALGVLRLDYDYPPSPGDVDSPLSWNYRVYYRCVPGFTFAMCRSNDLSPEV
LVQLEAAVKWLVAKGCSGITGDCGFMMQLQPLIRTRNKLTVFMSSLMQLPTVRAGFAPGEQIAIFSANGGDLNSMFE
TLKAECREDARDDAFVIVGCEDVDAFEAVESGEKVKVDRVQPGIVAKALAVVAAHPNLRAIVMECTELPVYSDAVRHA
TGLPVYDAITSCNMFMAGVQNNPRFGLEEWRHKFDGEQEVYRFGQHLSPAQQALLVNKPAAT

Pyramimonadophyceae

>C.tetramitiformis_A
MRVLSADAKQATQHSSASIPESDVRIRGLDDIDTLCRELIRAGLEIRDRSWLFTQYPRCFLGSDLVKLLISLKVAKDVPS
AVAVGNELISKHVFHHVWDYDIEMRDAYLFYRMSVHEEILGKRDLSSISHTFVLKTNERMNNPNTGVKLSEYQKGKYG
YTGTAAAEWLEKEGIARSQEEANAFGDLLIAFRLITTVVKQEQRPFRNDATLYRPCVKSSKKAGQSSETKSDTLGVLR
LDYGYPPIPGDIDHPNSFEYNVVYRKVKGLTFELAQSGEFKDDVEQNMLQAIRDLESLGAFGITGDCGFMANYQRFVA
EHAKVPVFMSSLCLIPFIFSSLRREGKLLVITANSDSLRALLPNVYDKPLEAEVSAQACAVTNDFLGEQLGCTILKPERII
NMGLQDLPGFDVVASATSLLSTEVDRVHVGNEIVKRVRDYLQQEPAIQAILLECTELPHFSSPLLRKYTRLPVFDALSV
CDFFQSSVEIKKSQQRRAVCVGHHCQRSLPDTPV

>C.tetramitiformis_B
MALGARGIWGLALNGVQAAPPVQDQDEVMEETVSMEDTSTVAAPKIKGKQRQSVWCHKEFDSSKVIGVLRIDYDYP
PAVGDIAHPDSFGYTVHYETVHGLTFERCQAGALDDEIMTNLQNSILKLQEYPGLVGITGDCGFMMHYQCPVRFMAK
VPAFMSSLIQCPAIGAIFKLFEKVLVLSANSKSLEPMKETLLTQAGFDVDDPSRYEVYGLQDLDGFDAVAKGEAVDLPR
VEVAITEAVTKKLAGDASIKAILLECTELPAYADALRRATGLPVFDAITAVNFFRNATVSSHWNKKAFIPHNPAYWTTKY
SKFGNEDQEAVEEVEEVEDETLSAPPFPCEDCDAIKIIGVLRIDYNYPPAPGDIDHPDSYGYKVVFRVVHGLTFAKCQD
GVMDDQIAESFRRAVDELEAVEGLVGITGDCGFMMHYQCLVRHMAQKPVFMSALLQAPLISAAFKNFEKILVVTANST
TLLPARDTLLTTCGFKVDDPEQFVIHGLQDLEGFDAVEKGEEVDTLLVQHHITERIKQVVKDEPSIRAILLECTELPHYA
DRLRRKIGLPVFDAITCVNYFRSSAVTSHWNKAFPPHNPDFWKDKFAKFGLEEKNMAEAAK

>C.tetramitiformis_C
AGVGVLRLDYHYPPLAGDIDSADSYGYKVIFKQVDGLTFEMAQSGKMTKAVEKNLVSAIHWLESQEVVGIAGDCGFM
MAYQVFVRRQTDVPVFMSSMLQAPMLIAAFDQDAKFGILTANSKSLEPNLDMLLQDCGVDVDVDRFVVLGLQNVPGF
DAVAKGEAVDASVVQPGIVAAVQKMCTDEPDVAAIILECTELPAYADAIRAGTGIPVFDAITLVDYFHSSMADNPRFGT
VYTTNPSTKITKALAAASNQDSA

>C.tetramitiformis_D
MAAAVGAKQKKFDVEFAEGADHPDLGPVAVGVLRLEYHYPPLEGDIDSADSYGYEVLFKQVDGLTFEVAQSGKMTK
NVEVNMISAIRWLEAKQVVGITGDCGFMMAYQVFVRRNTELPVFMSSMMQAPMLIAAHDQDAKFAIFTANSKSLEPN
LEFLLTDCGVEVDVDRFVLVGLQECDGFDXVAKGERVNPRLVQPAILKRVRKLVKDEPELAAIILECTELPAYADAIRAE
TGLPVFDAITLVDFFHSASSDNPHFGGAMKKGSFSGKKKGKGAVSKAMKKRFRPGADHPDLGPVGTIPAACSWLCD
CIPPVGFKARL

>C.tetramitiformis_E
MVDYFHSSCADNPNFGCSYQQQEQGHLKWQQEFAEGADNPDLGPVAIGVLRLDYHYPPLGGDIDSQDSYGYQVVF
KQVDGLTFEMAQSGRMTKAVEKNMKGAIQFLEEKDVVGITGDCGFMMAYQVFVRQQTDLPVFMSSMIQAPMLVASN
DSEAKFAILTANSKSLAPSLDWLLSECGVTVDVDRFVVVGLQDVPGFDAVAKGEAVDPKVVQPGIIDMLRETQKDEPQ
IAGIILECTELPAYADAIRSVLYIPVCLFIVSL

>C.tetramitiformis_F
AKKRTRKNKFREGADNPNLGPIAVGVLRLEYHYPPLAGDIDSEDSFGYKVFFRQVDGLTFELAQRGRMNKVVEKNMV
DAITWLEAKQVVGITGDCGFMMAYQVFVRQHTDLPVFMSSLLQAPILVAAHDREAVFALVTANSDSLEANIDTLLEDV
GMDVDADRFKIVGLQDVPGFDAVAKGEAVDPLVVQPGIIARLTQLADDEPDLQAVLLECTELPHYADAIRAELGLPVFD
VITLTDFFHSSRADNANFGQSFQPRIDNDDRTLMTMD

<u>Chlorodendrophyceae</u>

>P.subcordiformis
MSELMARRVFHHVWETGFDFKNSTLFYRFSFHEQLLGGADLTNISHTYVLKTCERMKGQVEMVEYKDGHHAFTGRA
GVEWLMKEGTVLTEAEATQLCNLFVACRLIAPVVSKTPLAPFSDSVLYRLVSAGTTGNTKEAAKTSLGVLRLEYGYPPI
PGDIDHPNSFPYSVVYRQVPGLTFELAQSGGLPDDVRAAFLNAIRDLEKLGVFGITGDCGFMANYQRFVADSTTACPV
FMSSLCLIPSIMAGMKTDAKLVVVTANSESLSVLFPNRFDAPLPESEHHRCAATNKFLSQQMGCTVARPECIVNMGLQ
DVIGFDVVAAATSLLSPEVDRVLVGQEIAKRVVDFIGSDTAVQAILLECTELPHFSAPMLRHYTQLPVFDALSVCDFFQA
ATDYGPEQTTRRAHCVGQHCQRGLPDTLVSL

>T.striata_A jgi|Tetstr1|462051|TSEL_007121.t1
MAGKHFDRQFQLKQAPSTQSAGSRESGLDDLDVIARRLMGSGQPADGAAVFHHVWEAGFEFKNSTLFYRFSFHEEL
LGGTDLTNISQTYVLDTCKRAKGNVEMVEYKSGKFAFQGKALVDWMLKEGTVLTQEEAMKLANLFVACKLITKVVSD
KGPMQAFSLSCLYVLTIKPSEGDKAKANSALGVLRLEYGYPPIPGDIDHPNSFPYKVVYRQVPGLTFELAQSGALPDE
VRESFVTAIRDLERLGVFGITGDCGFMANYQRFVSETTSVKPVFMSSLCLIPSIMAGLKADAKLVVVTANSDSLAAIFPN
RYDLPLPEEEFHKCSVTNEFMMAQMGCSIARPERIINMGLQDVTGFEVVAEATSLLSPDVDRVLVGNEIAKRVVDFLEI
DSTVQAILLECTELPHFSAPLLRHYTQLPVFDALSVCDFFQDASDSGEGQAGRRAACVGQHCQRGLPDTLIN

>T.striata_B jgi|Tetstr1|454920|TSEL_041783.t1
MAGKHFDRQFQLKQAASTQSAGSRESGLDDLDVIARRLMGEVEIKDRTWLFKTYKRCFLGTDLVKAMVKLQIAPDVK
GAVAVGNQLMERRVFHHVWEAGFEFKNSTLFYRFSFHEELLGGTDLTNISQTYVLDTCKRAKGNVEMVEYKSGKFAF
QGKALVDWMLKEGTVLTQEEAMKLANLFVACKLITKVVSDKGPMQAFSLSCLYVLTIKPSEGDKAKANSALGVLRLEY
GYPPIPGDIDHPNSFPYKPSATWSGLGVFGITGDCGFMANYQRFVSETTSVKPVFMSSLCLIPSIMAGLKADAKLVVVT
ANSDSLAAIFPNRYDLPLPEEEFHKCSDVTGFEVVAEATSLLSPDVDRVLVGNEIAKRVVDFLEIDSTVQAILLECTELP
HFSAPLLRHYTQLPVFDALSVCDFFQDASDSGEGQAGRRAACVGQHCQRGLPDTLIN

>T.striata_C jgi|Tetstr1|422311|TSEL_013155.t1
MAEVAVANHTWFFKTYRRSFTGDDLIRAIVRLRIAPDTKAALMVGNALMRHHVFHAVWGGDVEVMSTALLYRFSFHE
ELLGGVDLTNISVTYVLKTAERMEKHIDMVGYDKAGRPMFQGQSMLEWLTKEGTVLTEEEGMQLCNLFVACKLTSCV
TDKDCRAPFSPAALYIMSRRSARASNKETEKAAPLGVLRLEYGYPPIPGDIDHPSSFPYDVIYRKVPGLTFELAQSGDL
PEHVRDCGFMANYQRFVAESTAAAPVFMSSLCLIPTIMAGMKVDAKLLVLTANSALLAALLPNRYDAPLAPAEWGKCE
VTNAFLRKQCGCAVERPERIVNMGLQDLPGFQVVAEATSLLSPEVDRVLVGGGIAERVVSYIARDATVQAILLECTELP
HFSAPLLRNCTGLPVFDALSVCDFFQAAVDHGAEQAGRRALCLSRGRQRS

>T.striata_D jgi|Tetstr1|425072|TSEL_015536.t1
MAEVAVANHTWFFKTYRRSFTGDDLIRAIVRLRIAPDTKAALMVGNALMRHHVFHAVWGGDVEVMSTALLYRFSFHE
ELLGGVDLTNISVTYVLKTAERMEKHIDMVGYDKAGRPMFQGQAMLEWLTKEGTVLTEEEGMQLCNLFVACKLTSCV
TDKDCRAPFSPAALYIMSRRSARASNKETEKAAPLGVLRLEYGYPPIPGDIDHPSSFPYDVIYRKVPGLTFELAQSGDL
PEHVRDCGFMANYQRFVAESTAAAPVFMSSLCLIPTIMAGMKADAKLLVLTANSALLAALLPNRYDAPLAPAEWGKCE
ITNAFLRKQCGCAVERPERIVNMGLQDLPGFQVVAEATSLLSPEVDRVLVGGGIAERVVSYIARDATVQAILLECTELP
HFSAPLLRNCTGLPVFDALSVCDFFQAAVDHGAEQAGRRALCLSRGCQRL

>T.suecica
TIVSDGCPRERAILRNHVPLTVIVIVSLLRFGSGMLTRWCDALLCTGDCGFMANYQRFVADSTTACPVFMSSLCLIPSI
MAGMKTDAKLVVVTANSESLSVLFPNRFDAPLPESEHHRCAATNKFLSQQMGCTVARPECIVNMGLQDVIGFDVVAA
ATSLLSPEVDRVLVGQEIAKRVVDFISSDTAVQAILLECTELPHFSAPMLRHYTQLPVFDALSVCDFFQAATDYGPEQT
TRRAHCVGQHCQRGLPDTLVSL

Ulvophyceae

>U.mutabilis_A jgi|Ulvmu1|5019|UM021_0036.1
MEGKMKALSTEERHTFSKIIKKLACVARKVDDAVFGPADCGKKRAVTDKAPTAADTEAGAPAEAGGELEPERLLPRLR
PQVKLGILRLDYNYEAAPGDIDYPMSFKYKVIYRVVPGLTFEMCQSGKLTAAVDRAFREAIRWLDEEQNVDAITGDCG
FMFWFQDMARRQTSKPVLLSSLVQLPAITCGFAKHKKIAIFTANGHELMPMCELINKECGVNIEDSRFHIVGCEDVPGF
DAVAQGRKVDVGRVQPGIIELARHVTTQDLNVRAILLECTELPPYADALRRYLNIPVYDAVTCADMFMDGLQDNPRFG
DMDWREPFDGNDGEYEFGGELEEGELEHCINCIRKMDAQEKAQQLTA

>U.mutabilis_B jgi|Ulvmu1|6706|UM030_0039.1
MDTNLLKAVKELPPQSRRKLLQLADNLGMHLRKMDLDDPSGSKANPEQPKLGVLRLEYDYEPAVGDIDHPGSFAYPV
EYCTVKGLSFKICQAGERNEELDKNFQDAVQWLIDQGCDAITGDCGFMYWYQEDVRLVTDRPMMLSALCQLPAVTA
AFDDKEKIAIFTANSETLKPMLPKIKKNAGVDLEKDRYVIIGCQDVPGFDAVEKGEPVIYDEVAPGMVDLTRKVLEDYPS
ITAVLLECTELPQFADVLRKEFKLPVYTAVTCTDHFMAGLLDNPRFGLQDWHDEATVGKEKRKEKRSGMEGDAAAAG
GSEQRPVEDILKGLQMLVEA

>U.prolifera_A
MEEKFKALSKEEQHTLSRVLKKVACVARKFDQNVLKKPAAYIAGRGVDDDKPTNDAQASAGTHAPADSPKPSNFMP
HPRPQLKLGILRLDYNYEAAPGDIDYPKSFKYKVVYRVVPGLSFEMCQSGKMTAAVERAFKDAIRWLDEEKSVDAITG
DCGFMFWFQEMARRQTSKPVLLSSLVQLPAITCAFAKHKKIAIFTANGHELMPMCELINAECGVNIEDKRFCIVGCEDV
PGFDAVAQGCKVNVGKVQPGIIELARKVVTEDLNIRAILLECTELPPYADALRRHVNLPVYDAITCADMFMEGLQDNPL
FGDMEWREPFDGLHEEYAFGGELESGELENCINCMLKLDADGKAVPATG

>U.prolifera_B
MESHLSKAIDSMTPQARHKLMRIADHLTMTLRKSDMKPSTEDGVPEASVTQPMLGVVVLDYDYEPAIGDIDHPGSYS
YPVLYRKVPGLTFELCQAGGHTPEVVQNFKDAVKWLVEQGCDAITGDCGFMYWYQEHARAITDRPILLSALCQLPAV
APAFADYEKIAIFTANGKSLAPMLDLIKSSSGIDLESDRYIIVGCENVPGFDAVANGDPVIYDDVAPGMVDIAKKTLATNP
DVKAFLFECTELPQFADAVRAETGLPVYDSITCTNAYMSGLMDNPRFGLNEWQDEAEDAQESRKQHKKGCKHNKKH
GRGRKHAHRSVEDIVSALHVLLEGQ

>U.prolifera_C
MESVLSKAIDSMTPQARNKLLRIADHLTMTLRKSDIKPSTEDGVPEASIAQPMLGVVVLDYDYEPAVGDIDHPGSYSYP
VLYRKVPGLTFELCQAGGHTPEVVQNFKDAVKWLVEQGCDAITGDCGFMYWYQEHARAITDRPILLSALCQLPAVAP
AFADDEKIAIFTANGESLAPMLDLIKSSSGIDLSSDRYLIVGCEDVPGFDAVANGDPVIYDVVAPGIVDMAKKTLATNPD
VKAFLFECTELPQFADDVRAETGLPVYDSITCTNAYMSGLMDNPRFGLNEWQDEAEDAQESRKQHKKGCKHTKTHG
RGRKHAHRPVEDIVAALKVLLEGH

>U.prolifera_D
MVADGSVSASSPLAIAVAGLSPESRLKLLRVADDLVMHLRKSELPTPPAAAPSDFKFVQYSDSGGLQAVPPTPIGATA
VLSPKLGVIRLDYDYPPALGDVDHEDSFEYPVEYEIVEGLTFKLCQEGDLSKHPHVIANFESAVEKLYEKGCDVIIGDC
GFMFQFQHHAVTLPSRPSMMLLSPVSQLPSLISGLDPGKKVAIMTANSTTFNGMLELLAKEEADSGHASKYSISRAPR
DRYIVVGCQDVPHFGDEVARGEAIVYEDAERGIVQLVQDTLRKHPEVDMFLLECTELPQFANTLRFKTGKPVYTAITLA
NMAMSGRQSNPRFGRQYWAGMWLEQQRGRMAMRLRREGDPAGARAAAEGMRDTELMRSIANAPLLQHASGAHA
LSVLSQLADL

>U.prolifera_E
MDEDGNGAAMSPLASAVASLDPESRLKLLRVADDLVMHLMKSELPAPSRPEEIAQHGDSGGLQAVRSALTTVEQAAI
LNPKLGVLRIDYKYPPALGDVDHKDSFEYPVKYKKVKGLTFELCQKGDLTDHPQVNKNFKKAVNKLIEKGCDVIIGDCG
FMFQFQQDAVTNPNRPSILLLSPLSQLPTLITALGPEKKVAIMTANSESFEHMFKLLEKQEKISGHTSAYSISRAPRDRY
VVVGCQGVKHFGEEVANGWPIVYKLAEPGIVALAKDTLRKHPEVDMFLLECTELPQFANTLRYKTGKPVYTAITLANM
AMSGRRSNPRYGRQYWEKVWLEQQRRRMAMWLQREGDPGGAQAAAEGLRMRDTELMQSIADAPPPQHSNGDDL
EVLRQLADL

>U.prolifera_F

MDADGNGDAIAMSRDPLASAVASLDPESRLKLLRVADDLVMHLRKSELPAPPAPGEFVQRDESGGLALQLQAVPLTP
VEDTAILNPTLGVLRIDYKYPPALGDVDHKDSFEYPVKYKKVKGLTFELCQKGDLTDHPQVDQNFKKAIKKLHDKGCD
VIIGDCGFMFQFQQDAVTIPNRPSILLLSPLSQLPTLITALGPKKKVAIMTANSKSFEHMLELLKKQERSCGHTSAYSISR
APRDRYVVVGCQDVPYFGCEVAHGEPIVYELAEPGIIKLAQDTLRKHPEVDMFLLECTELPQFTNTLRYKTEKPVYTAI
TLANMAMSGRLDNPRFGRHKWRREWLEQQRGRMAMRLRREGDPAGAQAAAEGLRDTELMQSIANAPPPQHAAAR
RSTPQHADGDHADGDHMHVLNVLGQLADL

Anthozoa

>Acropora_millepora (Identical to A.milleporaH)
MSGETAKSAAACLGVIRLDYDYPPAPGDIDHPDSFSCDVYYKVVPGLTFEMCQKGQMTKEVQKRFKQSIKWLVKEKN
VNGITGDCGFMMNFQGFARKVTKIPIFMSSLCQLPAVTCGFAQKEQIIIMTANGKSLEPMRDLIRDECGVDTQDKRYNI
VGCEDVPHFGPAVANGDKVDTKKAQPGIVKKAVEALKKYPHSRAFLLECTELPPYSDAIRFHTGLPVYDSITACHFFIS
GHKDNVRFGLQDWQDDWDEEQEEYDYGDNLNRKEKKELVNKVE

>A.millepora_A Amillepora37625-RA protein AED:0.02 eAED:0.02 QI:0|0.5|0|1|0|0|3|0|326
HNSIAAVNTSFEVIEQLSIFRRLFDVRRIPQKKMSGEKKKSAAACLGVIRLDYDYPPAPGDIDHPDSFNCDVYYKVVPG
LTFEMCQKGEITDAVKQRFEESIKWLVEEKKVKGITGDCGFMMNFQSLARNITKIPIFMSSLCQLPAVTCGYAQKEQMI
IMTANGKSLEPMRDLIRDECGVDTQDKRYNIVGCEDVPHFGEAVAKGNKVKVEDATPWIVRKAVHALLKYPKSRAFLL
ECTELPPYSDAIRFYTGLPVYDSITACHFFISGHKDNPRFGLQDWQDQWDGKQDKYTYGANLTEEEQKELAPPSFES
LPIEFSRKKSLFR

>A.millepora_B Amillepora26932-RA protein AED:0.26 eAED:0.26 QI:0|-1|0|1|-1|1|1|0|278
MSGKTKKSSAASLGVIRLDYDYPPAPGDIDHPDSFSCDVYYKVVPGLTFEMCQKGKITDEVKERFEESIKWLVQEKKV
KGITGDCGFMMNFQSLARNITKIPVFMSSLCQLPAVTCGYAQKEQMIIMTANGKSLEPMRDLIRDECGVDTQDKRYNI
VGCEDVPHFGEAVAKGEKVKVEDATPWIVRKAVHALFKYPKSRAFLLECTELPPYSDAIRFYTGLPVYDSITACHFFIS
GHKDNPRFGLQGWQDQWDGKQDEYTYGANLTEEEQQELVNPCK

>A.millepora_C Amillepora26933-RA protein AED:0.00 eAED:-0.00 QI:0|-1|0|1|-1|1|1|0|246
MSGETRKSVAACLGVIRLDYDHPPAPGDIDHPDSFNCDVYYKVVPGLTFEMCQKGQMTEEVEKRFEDSIKWLVQEKK
VKGITGDCGFMMNFQSLARNTAKISVFMSSLCQLPAVTCGFDQKEEIIIMTANGKSLKSMRGLIRDECGVDTQDKRYNI
VGCEDVPHFGEAVAKGEKVKVEDATPWIVRKAVHALVKYPKSKAFLLECTELPPYSDAIRFYTGLPVYDSITACHFFIS
GHRDCILFVRI

>A.millepora_D Amillepora34779-RA protein AED:0.25 eAED:0.25 QI:0|-1|0|1|-1|1|1|0|284
MSGETAQSAAPCLGVIRLDYDYPAAPGDIDHPKSFSCDVYYKVVPGLTFGMCKEGKPMPPEVETRFKESIKWLVEEK
KVKGITGDCGFMMNYQKLAREVTKIPIFMSSLCQLPAVTCGYAENEQIIIMTANGKSLEPMRDLIRDECGVDTQDKRYN
IIGCEDVPHFGKAVEDGHKVKVEDATPWIVRKAVHALLKYPKSRAFLLECTELPPYSDAIRFYTGLPVYDSITACHFFIS
GHKDNPRFGLKDWQDEWDGVQEKYKYGDNLTPEEKEQLKHKVKQGHSI

>A.millepora_E Amillepora18269-RA protein AED:0.26 eAED:0.26 QI:0|0|0|1|1|1|3|0|789
MTDDVEKRFKQSIKWLVKEKKVNGITGDCGFMMNFQSIARQITKIPIFMSSLCQLPAVTCGYAQHEQMIIMTANGKSLE
PMRDLIRDECGVDTQDKRYNIVGCEDVPHFGQAVANGDKVNVKDATPWIVRKAVHALFKYPKSRAFLLECTELPAYS
DAIRFYTGLPVYDSITACHFFISGHKDNPRFGLDDWQDEFDGKQEDYEYGDNLTKKEKAALVNKVRRILQKNMSEETK
KAAAACLGVIRLDYDYPPAPGDIDHPDSFNCDVYYKVVPGLTFEMCQKGKMTDDVEKRFKQSIKWLVKEKKVNGITG
DCGFMMNFQSIARQITKIPIFMSSLCQLPAVTCGYAQHEQMIIMTANGKSLEPMRDLIRDECGVDTQDKRYNIVGCED
VPHFGQAVANGDKVNVKDATPWIVRKAVHALFKYPKSRAFLLECTELPAYSDAIRFYTGLPVYDSITACHFFISGHKDN
PRFGLDNWQDEFDEEQEEYQYGDNLTKKEKAALVNKVKRIPQKKMSGETKKSAAACLGVIRLDYDYEAVPGDIDHPD
SFNCDVYYKVVPGLTFEMCQKGKMTDDVEKRFKQSIKWLVKEKKVNGITGDCGFMMNFQHIARQITKIPIFMSSLCQL
PAVTCGYAEKEQMIIMTANGKSLEPMRDLIRDECGVDTQDKRYNIVGCEDVPHFGQAVANGDKVNVKDATPWIVRKA
VHALFKYPKSRAFLLECTELPAYSDAIRFYTGLPVYDSITACHFFISGHKDNPRFGLDDWQDEFDGKQEDYDYGDNLT
KKEKAALVNKA

>A.millepora_F Amillepora26941-RA protein AED:0.00 eAED:0.00 QI:102|1|1|1|0|0|2|1016|302
MLHRGSLFLSLSRFSKVRRIPQKKMSAETKKSAAPRLGVIRLDYKYEAALGDIDHLDSFNFHVYYKVVPGLTFEVCQE
GKMSDEVKDRFKESIKWLVREKKVNGITGDCGFMMNFQDFARKITKIPIFMSSLCQLPAVTCSYEEKEQIIIMTANGKN
LEPMRDLIRDECGVDTQDKRYNIVGCEDVPYFGQAVANGDKVNIEEATPWIVRKAVHALFKYPKSRAFLLECTELPAY
SDSIRFYTGLPVYDSVTACQFFINGHTDNVRFGLQNWQDEWDGNQEEYQYGDNLTKEEQNQLVHKVE

>A.millepora_G Amillepora18267-RA protein AED:0.41 eAED:0.41 QI:0|-1|0|1|-1|1|1|0|278

MSEETKKAAAACLGVIRLDYDYPPAPGDIDHPDSFNCDVYYKVVPGLTFEMCQKGKMTDDVEKRFKQSIKWLVKEKK
VNGITGDCGFMMNFQSIARQITKIPIFMSSLCQLPAVTCGYAETEQMIIMTANGKSLEPMRDLIRDECGVDTQDKRYNI
VGCEDVPHFGQAVANGDKVNTKNATPGIVKKAVEALKKYPKSRAFLLECTELPPYSDAIRFHTGLPVYDSITACHFFIS
GHKDNPRFGLDNWQDEFDEEQEEYQYGDNLTKKEKAALVNKVK

>A.millepora_H Amillepora26930-RA protein AED:0.33 eAED:0.33 QI:0|-1|0|1|-1|1|1|0|278
MSGETAKSAAACLGVIRLDYDYPPAPGDIDHPDSFSCDVYYKVVPGLTFEMCQKGQMTKEVQKRFKQSIKWLVKEKN
VNGITGDCGFMMNFQGFARKVTKIPIFMSSLCQLPAVTCGFAQKEQIIIMTANGKSLEPMRDLIRDECGVDTQDKRYNI
VGCEDVPHFGPAVANGDKVDTKKAQPGIVKKAVEALKKYPHSRAFLLECTELPPYSDAIRFHTGLPVYDSITACHFFIS
GHKDNVRFGLQDWQDDWDEEQEEYDYGDNLNRKEKKELVNKVE

>A.millepora_I Amillepora37624-RA protein AED:0.26 eAED:0.26 QI:0|-1|0|1|-1|1|1|0|279
MSGETAKSAAACLGVIRLDYDYEAAPGDIDHPDSFSCDVYYKVVPGLTFEMCQKGKMTEEVEKRFKESIKWLVKEKS
VNGVTGDCGFMMNFQSIARKITKIPIFMSSLCQLPAVTCGYAQKEQIIIMTANGKSLEPMRDLIRDECGVDTQDKRYNIV
GCEDVPHFGEAVANGDKVNTKKATPGIVKKAVEALKKYPQSRAFLLECTELPPYSDAIRFHTGLPVYDSITACHFFISG
HKDNPRFGLQEWQADWDEDQEEYQYGDNLTQKEKDQLVNKVNS

>A.millepora_J Amillepora34780-RA protein AED:0.31 eAED:0.23 QI:0|0|0|0.33|0|0.33|3|0|309
MSGETGQSAAPCLGVIRLDYNYEAAPGDIDHPDSFNCDVYYKVVPGLTFKMCQTGVITVEVKKRLEESIKWLVEEKKV
NGITGDCGFMMYFQKLARKVTKIPIFMSSLCQLPAVTCGYAENEQIIIMTANGNSLEPMRYLIRDECGVDTQDRRYNIV
GCEDVPYFGEAVAKGEKVNTTEAQTGIVKKALEALRKYPQSRAFLLECTELPPYSDAIRFATGLPVYDSITACHFFISG
HKDNVRFGLEDLDWQADFDEEQEEYNYGDNLTPEEKKHLIKGRHRNVHFNLQVCLSRRYLTDFLNLKDQEIRM

>XP_015777049.1 closest to A. millepora B PREDICTED: dimethlysulfonioproprionate lyase 7-like [Acropora digitifera]
MRSLARKLQKKKNKSGDSLEENNKEKLRGNTSETDNKIKEEKSASPRLGIIRLDYDYTPADGDIDCPDSFDYDVYYKV
VPGLTFEVCKEGKMTEEVENRFKESIKWLVEEKKVNAITGDCGFMINFQGIARQVTKIPVSMSSLCQLPTVTCSYAEN
EQIIILTANGKDLRHMKNVISVECGVDTEDLRYNIVGCEDVPYFGEAVAKGEKVKVKDAQPWIIRKAFNALLKYPKSKAF
LLECTELPPYANAIRFYTGLPVFDAITACHFFISASKDNTMFGLQGWQDEWDGKHEAYHYGDNLTEAEKKELVNPVHS
E

>XP_015777050.1 closest to A. millepora B PREDICTED: dimethlysulfonioproprionate lyase 7-like [Acropora digitifera]
MLSLARKLLKKKNKSGDSLEENNKEKLRGNTSETDNKIKEEKSASPRLGIIRLDYDYTPADGDIDCPDSFDYDVYYKVV
PGLTFEVCKEGKMTEEVENRFKESIKWLVEEKKVNAITGDCGFMINFQSIARQVTNIPVSMSSLCQLPTVTCSYAENE
QIIILTANGKDLRHMKNVISVECGVDTEDLRYNIVGCEDVPYFGEAVAKGEKVKVKDAQPWIIRKAFNALLKYPKSKAFL
LECSELPPYANAIRFYTGLPVFDAITACHFFITASKDNTMFGLQGWQDVWDGKHEAYHYGDNLTEAEKEELVNPVHS
D

>XP_015765068.1 closest to A. millepora J PREDICTED: dimethlysulfonioproprionate lyase 7-like [Acropora digitifera]
MSGETTQSAAPCLGVIRLDYNYEAAPGDIDHPDSFNCDVYYKVVPGLTFDMCKKGVITVEVKKRLEESIKWLVQEKKV
NGITGDCGFMMYFQKLARKVTKIPIFMSSLCQLPAVTCGYAENEQIIIMTANGKSLEPMRYLIRDECGVDTQDRRYNIV
GCEDVPYFGEAVAKGEKVNTTDAQTGIVKKALEALSKYPQSRAFLLECTELPPYSDAIRFATGLPVYDSITACHFFISG
HKDNVRFGLEDLDWQADFDEEQEEYKYGDNLTPEEKKHLVNKPEKGHSK

>XP_015765069.1 closest to A. millepora D PREDICTED: dimethlysulfonioproprionate lyase 7-like [Acropora digitifera]
MSGKTAKSAAASLGVIRLDYNYEPAPGDIDHPDSFNCDVYYKVVPGLTFKMCKGEIPSTDEVETRFKESIKWLVEEKK
VKGITGDCGFMMYYQSLARKVTNIPIFMSSLCQLPAVTCGYAENEQIIIMTADGGCLEPMRDLIRDECGVDTQDKRYNI
VGCEDVPHFGQAVKKGDKVNVKEATPWIVRKAVHALLKYPKSRAFLLECTELPPYSDAIRFYTGLPVYDSITACHFFIS
GHKDNPRFGLKDWQDQWDGKQDEYKFGDNLSPEEKKHLKFKAKQGHST

>XP_015780071.1 closest to A. millepora H PREDICTED: dimethlysulfonioproprionate lyase 7-like [Acropora digitifera]
MSGETAKSAAACLGVIRLDYDYPPAPGDIDHPDSFSCDVYYKVVPGLTFEMCQKGQMTKEVQKRFKQSIKWLVKEKN
VNGITGDCGFMMNFQGFARKITKIPIFMSSLCQLPAVTCGFAQKEQIIIMTANGKSLEPMRDLIRDECGVDTQDKRYNIV
GCEDVPHFGPAVSNGDKVDTKKAQPGIVKKAVEALKKYPHSRAFLLECTELPPYSDAIRFHTGLPVYDSITACHFFISG
HKDNVRFGLQDWQDDWDEEQEEYDYGDNLTRKEKKELVNKVE

>XP_015760108.1 closest to A. millepora C PREDICTED: dimethlysulfonioproprionate lyase 7-like isoform X1 [Acropora digitifera]

MSGETRKSVAACLGVIRLDYDHPPAQGDIDHPDSFNCDVYYKVVPGLTFEMCQKGQMTEEVEKRFEDSIKWLVQEK
KVKGITGDCGFMMNFQSLARNTAKISVFMSSLCQLPAVTCRYAEKEQIIIMTANAKSLESMRGLIRDECGMDIQDKRYN
IVGCEDVPHFGEAVAKGEKVKVEDATPWIVRKAVHALVKYPKSKAFLLECTELPPYSDAIRFYTGLPVYDSITACHFFIS
GHRDCILFVRI

>XP_015768696.1 closest to A. millepora G PREDICTED: dimethlysulfonioproprionate lyase 7-like [Acropora digitifera]
MSGETKKSAAACLGVIRLDYDYEAVPGDIDHPDSFNCDVYYKVVPGLTFKMCQKGKMTDDVEKRFKESVKWLVKEK
KVNGITGDCGFMMNFQSIARQITKIPIFMXSLCQLPAVTCGYAEKEQIIIMTANGKSLEPMRDLIRDECGVDTQEKRYNI
VGCEDVPHFGQAVANGDKVNVEEATPWIVRKAVHALLKYPKSRAFLLECTELPAYSDAIRFYTGLPVYDSITACHFFIS
GHKDNPRFGLDNWQDEFDGKQEDYEFERNLTENEKNALVNKVK

Species we used MMETSP for expression, already defined above:

>CAMPEP_0187698862 Alma 4/5 like /NCGR_PEP_ID=Emiliania-huxleyi-CCMP370-20130905|29607_1 /TAXON_ID=2903 /ORGANISM="Emiliania-huxleyi-CCMP370" /LENGTH=645 /DNA_ID=CAMNT_0030801207 /DNA_START=1 /DNA_END=1935 /DNA_ORIENTATION=+
XPETRRGDTDPTWAARTPPQRPRARRSSPGERSNPEPSAPASLSSGSYAATEHDGSAAQAIVSAADRAEYHYRIVY
AVVEGLPYQHVRAGKPLSASQEEALRVAVEELDLAECVCITGDCGSFVHYQGAVRSLTTRAVVLSPLVQASLIATMYQ
RTEKVIVVTNDTNDYSQADLEQNLVAIGLEQDDAKRFIVVGLQHIRGFATADVADQEVSEWGLVDESVTTVERKAIITT
VERKAKERNAKAIILESTLLPSFSDALRARLGLPVFDALTLVDYFVHASTDNPRFGLDFDERQAGRCNAYDGLDPAKLP
AIGILRIDYEYPPALGDIACEKSYGYRTTHEVAEGLTFEVAQEGGPLCSEKRENVAAALQRLQATPGVVGIAGDCGFLM
NYLADARSVSKLPCFISALIQCDLMATAYGDEAARASLSQAHAQAQAQARQPRPCGAPEGATAVGXSDAALPAGRAL
LLLLALRNYLVLTANGAALEPKFAEMLALANVTEAEVQARFHVLGLEDVDGFDAVAKGEAVDTARVEPGIVALAKAAVA
RFPDVRAILLECTELPPYADALRSALGVQVFDAITLVDYVHSSTADNPHFGVKFQTTVHAASTARFGRDSARNSERNS
VPETPSTPSSTPWRKSSPTL

>CAMPEP_0187624632 Alma 4/5 like /NCGR_PEP_ID=Emiliania-huxleyi-379-20130905|8113_1 /TAXON_ID=2903 /ORGANISM="Emiliania-huxleyi-379" /LENGTH=710 /DNA_ID=CAMNT_0030721735 /DNA_START=1 /DNA_END=2133 /DNA_ORIENTATION=+
XRPGGCPARLTQARNTNLRGEAAHAARWGGTGRAPSNPPALGLGFDAWGHSRETRELAGAVGEGATPCTDGRPP
ENTPAWIASHQSSLPNAAKGYPPREPRENTKFGGPDPTSTGEKAETRRGDTDPTWAARTPPQRPRARRSSPGERS
NPEPSAPASLSSGSYAATEHDGSAAQAIVSAADRAEYHYRIVYAVVEGLPYQHVRAGKPLSASQEEVLRVAVEELDS
AGCVCITGDCGSFVHYQGAVRSLTTRAVVLSPLVQASLIATMYQRTEKVIVVTNDTNDYSQADLEQNLVAIGLEQDDA
KRFIVVGLQHIRGFATADVADQEVSEWGLVDESVTTVERKAIITTVERKAKERNAKAIILESTLLPSFSDALRARLGLPVF
DALTLVDYFVHASTDNPRFGLDFDERQAGRCNAYDGLDPAKLPAIGILRIDYEYPPALGDIACEKSYGYRTTHEVAEGL
TFEVAQEGGPLCSEKRENVAAALQRFKRRLASSALRVIARLPHELSRRRTLRLQAAVLYLGPHPVRLDGDRLRRRGE
ALVLTANGAALEPKFAEMLALANVTEAEVQARFHVLGLEDVDGFDAVAKGEAVDTARVEPGIVALAKAAVARFPDVRA
ILLECTELPPYADALRSALGVQVFDAITLVDYVHSSTADNPHFGVKFQTTVHAASTARFGRDSARNSERNSVPETPST
PSSTPWRKSSPTL

>CAMPEP_0187639246 closest to Alma 6 /NCGR_PEP_ID=Emiliania-huxleyi-379-20130905|18426_1 /TAXON_ID=2903 /ORGANISM="Emiliania-huxleyi-379" /LENGTH=325 /DNA_ID=CAMNT_0030737195 /DNA_START=126 /DNA_END=1103 /DNA_ORIENTATION=+
MLRPKTPRIGIVRLDQPRTAKTAVLGDLVAPQSLQGVAPQSLQGSESLSQTDIIYRRAEGYTFSVCVSGHPVNAPDQL
QLTRGEVRIRGTQGDPQHLFEVGGNQPPCARNPSFTEPWVVNGRAVGVVYIYLPELILANLKQTLAELRELECRCLSS
SCGFMANINSFCAARSGMPTLMSTLDLLPTLLLSLPPEDVILVLTSNGTNFKRHAETLIPPHVPLKRLRVCGLEQVRGF
GAEVASGTTVDPLLAEAGIVGAVVEAFTATAPSPIGAILLECTELPGYSNALRQKFNLPVYDVLTLCSLLIASVAVSPTLG
AHAHFVNEM

>CAMPEP_0187748300 Alma 3/6 /NCGR_PEP_ID=Emiliania-huxleyi-PLYM219-20130905|8612_1 /TAXON_ID=2903 /ORGANISM="Emiliania-huxleyi-PLYM219" /LENGTH=361 /DNA_ID=CAMNT_0030854097 /DNA_START=19 /DNA_END=1104 /DNA_ORIENTATION=+
MGCAGSTLRSGASFEDSRLAAIESSRFHEVGHHAQFDEGGRFKQLPPADDDAKLLVANHPSLGVIRLDYDYPPALGD
VDHPGSFYYDVFYRVVPGLTFELCQSGELPDDVKQRFIDAITWLDEQGVAGITGDCGFFMYFQALARSVTSKPVFMS
SLCQLPAVVCAYAADEQIALFTANGESLKPMRELIKKECGVDPDDTRFVIVGCEDVPGFEAVANGDRVDVDSVVPHLV
RLAEDTVAKHAGTAKPIRAILFECTELPPYSDAVRAATRLPVFDSITCCNSMLASLMDNPRFGVNNWHLSWDGAHTTR
RFGDNVPPHLKGKLVNREHPENVARWNANLAERSSFSSAQQESIGRGSREL

>CAMPEP_0187787020 Alma 4/5 like/NCGR_PEP_ID=Emiliania-huxleyi-PLYM219-20130905|91030_1
/TAXON_ID=2903 /ORGANISM="Emiliania-huxleyi-PLYM219" /LENGTH=595 /DNA_ID=CAMNT_0030895381
/DNA_START=1 /DNA_END=1785 /DNA_ORIENTATION=+
XPETRRGDTDPTWAARTPPQRPRARRSSPGERSNPEPSAPASLSSGSYAATEHDGSAAQAIVSAADRAEYHYRIVY
AVVEGLPYQHVRAGKPLSASQEEALRVAVEELDLAGCVCITGDCGSFVHYQGAVRSLTTRAVVLSPLVQASLIATMYQ
RTEKVIVVTNDTNDYSQADLEQNLVAIGLEQDDAKRFIVVGLQHIRGFATADVADQEVSEWGLVDESVTTVERKAIITT
VERKAKERNAKAIILESTLLPSFSDALRARLGLPVFDALTLVDYFVHASTDNPRFGLDFDERQAGRCNAYDGLDPAKLP
AIGILRIDYEYPPALGDIACEKSYGYRTTHEVAEGLTFEVAQEGGPLCSEKRKNVAAALQRLQATPGVVGIAGDCGFLM
NYLADARSVSKLPCFISALIQCDLMATAYGDEAHFLVLTANGAALEPKFAEMLALANVTEAEVQARFHVLGLEDVDGF
DAVAKGEAVDTARVEPGIVALAKAAVARFPDVRAILLECTELPPYADALRSALGVQVFDAITLVDYVHSSTADNPHFGV
KFQTTVHAASTARFGRDSARNSERNSVPETPSTRPSSTPWRKSPPTL

>CAMPEP_0200904798 Similar to T. striata A/B (splice differences) /NCGR_PEP_ID=Tetraselmis-striata-LANL1001-20140214|2880_1 /ASSEMBLY_ACC=CAM_ASM_001234 /TAXON_ID=3165 /ORGANISM="Tetraselmis striata, Strain LANL1001" /LENGTH=503 /DNA_ID=CAMNT_0047091997 /DNA_START=225 /DNA_END=1736
/DNA_ORIENTATION=+
MAGKHFDRQFQLKQAASTQSAGSRESGLDDLDVIARRLMGEVEIKDRTWLFKTYKRCFLGTDLVKAMVKLQIAPDVK
GAVAVGNQLMERRVFHHVWEAGFEFKNSTLFYRFSFHEELLGGTDLTNISQTYVLDTCKRAKGNVEMVEYKSGKFAF
QGKALVDWMLKEGTVLTQEEAMKLANLFVACKLITKVVSDKGPMQAFSLSCLYVLTIKPSEGDKAKANSALGVLRLEY
GYPPIPGDIDHPNSFPYKVVYRQVPGLTFELAQSGALPDEVRESFVTAIRDLERLGVFGITGDCGFMANYQRFVSETT
SVKPVFMSSLCLIPSIMAGLKADAKLVVVTANSDSLAAIFPNRYDLPLPEEEFHKCSVTNEFMMAQMGCSIARPERIIN
MGLQDVTGFEVVAEATSLLSPDVDRVLVGNEIAKRVVDFLEIDSTVQAILLECTELPHFSAPLLRHYTQLPVFDALSVC
DFFQDASDSGEGQAGRRAACVGQHCQRGLPDTLIN

>CAMPEP_0200915634 Similar to T. striata A/B (splice differences) /NCGR_PEP_ID=Tetraselmis-striata-LANL1001-20140214|11490_1 /ASSEMBLY_ACC=CAM_ASM_001234 /TAXON_ID=3165 /ORGANISM="Tetraselmis striata, Strain LANL1001" /LENGTH=549 /DNA_ID=CAMNT_0047105949 /DNA_START=222 /DNA_END=1870
/DNA_ORIENTATION=+
MAGKHFDRQFQLKQAASTQSAGSRESGLDDLDVIARRLMGEVEIKDRTWLFKTYKRCFLGTDLVKAMVKLQIAPDVK
GAVAVGNQLMERRVFHHVWEAGFEFKNSTLFYRFSFHEELLGECGTPPHMCPPHQHGRLLSGAESRLMLLPAIVGG
GSXGWDGDMRIGGTDLTNISQTYVLDTCKRAKGNVEMVEYKSGKFAFQGKALVDWMLKEGTVLTQEEAMKLANLFV
ACKLITKVVSDKGPMQAFSLSCLYVLTIKPSEGDKAKANSALGVLRLEYGYPPIPGDIDHPNSFPYKVVYRQVPGLTFE
LAQSGALPDEVRESFVTAIRDLERLGVFGITGDCGFMANYQRFVSETTSVKPVFMSSLCLIPSIMAGLKADAKLVVVTA
NSDSLAAIFPNRYDLPLPEEEFHKCSVTNEFMMAQMGCSIARPERIINMGLQDVTGFEVVAEATSLLSPDVDRVLVGN
EIAKRVVDFLEIDSTVQAILLECTELPHFSAPLLRHYTQLPVFDALSVCDFFQDASDSGEGQAGRRAACVGQHCQRGL
PDTLIN

>CAMPEP_0200924274 Similar to T. striata C/D (partial) /NCGR_PEP_ID=Tetraselmis-striata-LANL1001-20140214|18675_1 /ASSEMBLY_ACC=CAM_ASM_001234 /TAXON_ID=3165 /ORGANISM="Tetraselmis striata, Strain LANL1001" /LENGTH=161 /DNA_ID=CAMNT_0047117247 /DNA_START=1 /DNA_END=482
/DNA_ORIENTATION=-
XWVVPFPRLQGDCGFMANYQRFVAESTAAAPVFMSSLCLIPTIMAGMKADAKLLVLTANSALLAALLPNRYDAPLAPA
EWGKCEITNAFLRKQCGCAVERPERIVNMGLQDLPGFQVVAEATSLLSPEVDRVLVGGGIAERVVSYIARDATVQAIL
LECTE

>CAMPEP_0200935740 Similar to T. striata C/D (partial) /NCGR_PEP_ID=Tetraselmis-striata-LANL1001-20140214|95585_1 /ASSEMBLY_ACC=CAM_ASM_001234 /TAXON_ID=3165 /ORGANISM="Tetraselmis striata, Strain LANL1001" /LENGTH=144 /DNA_ID=CAMNT_0047133793 /DNA_START=1 /DNA_END=431
/DNA_ORIENTATION=+
XVFHAVWGGDVEVMSTALLYRFSFHEELLGGVDLTNISVTYVLKTAERMEKHIDMVGYDKAGRPMFQGQAMLEWLT
KEGTVLTEEEGMQLCNLFVACKLTSCVTDKDCRAPFSPAALYIMSRRSARASNKETEKAAPLGVLRLE

>CAMPEP_0191203590 Similar to P. parvum A /NCGR_PEP_ID=Prymnesium-parvum-Texoma1-20131001|4355_1
/TAXON_ID=97485 /ORGANISM="Prymnesium-parvum-Texoma1" /LENGTH=583 /DNA_ID=CAMNT_0035116129
/DNA_START=1 /DNA_END=1750 /DNA_ORIENTATION=-
XGTLGIVRLDYNYPPAPGDIDHPASFAYDVFYKVVPGLTFEMCQSGKLTPEVEERFLNTIKYFEAKGVSGITGDCGFM
MYFQALARQATNKPVFMSALAQLPAVTAAFGNEELIAIFTANGSTLKPMEPMICDECGVNVEEQRYVIVGCEDVPHFE
AVARGEKVDVDKVTPGMIAKARDVLATYPNIRAILLECTELPPYADALRKETGLPVYDAITACDFFMMGVQDNERFGL
QDWQVVWDGQQEEYTFAQNLTEEEKAALVNVPKEERAGWLSSIGKTYASMSHVASTAKTMTEYVTSQALNATKAKE
APVRAKDGTLGIVRLDYNYPPAPGDIDHPASFAYDVFYKVVPGLTFEMCQSGKLTPEVEERFLNTIKYFEAKGVSGITG
DCGFMMYFQALARQATNKPVFMSALAQLPAVTAAFGNEELIAIFTANGSTLKPMEPMICDECGVNVEEQRYVIVGCED

VPHFEAVARGEKVDVDKVTPGMIAKARDVLATYPNIRAILLECTELPPYADALRKETGLPVYDAITACDFFMMGVQDNE
RFGLQDWQVVWDGQQEEYTFAQNLTEEEKKHLVNKVS

>CAMPEP_0191252250 Similar to P. parvum A /NCGR_PEP_ID=Prymnesium-parvum-Texoma1-20131001|104153_1 /TAXON_ID=97485 /ORGANISM="Prymnesium-parvum-Texoma1" /LENGTH=329 /DNA_ID=CAMNT_0035172997 /DNA_START=1 /DNA_END=984 /DNA_ORIENTATION=+
XKEPQGMGCCAAKEAKFDNVQQPEPTKATASAAPTTSSSVKPDVGADSPPLATSSCIKPPSETALSVKNVNISADKP
ARAKFGTLGIVRLDYNYPPAPGDIDHPASFAYDVFYKVVPGLTFEMCQSGKLTPEVEERFLNTIKYFEAKGVSGITGDC
GFMMYFQALARQATNKPVFMSALAQLPAVTAAFGNEELIAIFTANGSTLKPMEPMICDECGVNVEEQRYVIVGCEDVP
HFEAVARGEKVDVDKVTPGMIAKARDVLATYPNIRAILLECTELPPYADALRKETGLPVYDAITACDFFMMGVQDNERF
GLQDWQVVWDGQQEEX

>CAMPEP_0191271526 Similar to P. parvum B /NCGR_PEP_ID=Prymnesium-parvum-Texoma1-20131001|200449_1 /TAXON_ID=97485 /ORGANISM="Prymnesium-parvum-Texoma1" /LENGTH=726 /DNA_ID=CAMNT_0035195459 /DNA_START=1 /DNA_END=2179 /DNA_ORIENTATION=+
XRIAYALLGAMARRQQAEDGAELGKQRFTGAEALGWIVGQPWCVDEAHAKAVGDALVSAQLLLAADNPTPAFLLSPT
TEYTANYTFSISDLDVLCQRACAGVQYEPGDDKVFSGRALVTWLLDNGVATDRQWGTAAGAQLQQLKVLHSSQHSA
FASPAKLEFRDEPGRLYRFSVHEPNMRKIEGTDQISHSYLMDTAARMEAQLHPKPVGTSLHEVVAASVVAPRFCGDD
LTSWLLREGTARTHAEAVMMVEYMIAAQVLYPVGMKRPQSRSLQFAYTAADLRTFSADFEFSFVEPAKLARAAAYER
MQRSVEQGKVARSQLLLRHTIQRRLAMEHHGDLGDDLYELLYAKPKPAEERRAEPRHAVALAGDRHDAKAVRPLGVI
RLDYGYPPIPGDIDHPSSFDYPVVYRKVPGLLFEVAQEGVLTPEIRSGIEQAVRELEAHQVFGITGDCGFMANYQHLV
RSLASCPVFLSSLVLLPAISPLIGSRDRILCCTANGNSLAKLLPALWGAPPPPAAAYGVTHTFTHKHLDLHLLRPEQLVV
CGFERVPGFDVVAEASSLMSHAVDRELVEQXDHVDGAHDPLARPHDQDDPARVHGAATLHGDSAAGHRAARVRLTI
DLQLRACGVRCKPIGGIAPLTLSWTPLPDQIPGYAHDWMIHDGQLMVVCKHSEPCCRPVLQPFAHEGIRRDQHKGHN
VGGGVASRVKHYQVCDWHACRLVFYQLYEAP

>CAMPEP_0191230814 Similar to P. parvum B /NCGR_PEP_ID=Prymnesium-parvum-Texoma1-20131001|21691_1 /TAXON_ID=97485 /ORGANISM="Prymnesium-parvum-Texoma1" /LENGTH=784 /DNA_ID=CAMNT_0035146203 /DNA_START=1 /DNA_END=2353 /DNA_ORIENTATION=+
RGLGGLACMHRRQPRAKEDAPAHSAWNVTKLPLLHAAIADLLRAAQREAPDDPIAFASHYFLALQEEEGSPSEAREK
VAAAFAPSPLPLSPRERAPPLDRIAYALLGAMARRQQAEDGAELGKQRFTGAEALGWIVGQPWCVDEAHAKAVGDA
LVSAQLLLAADNPTPAFLLSPTTEYTANYTFSISDLDVLCQRACAGVQYEPGDDKVFSGRALVTWLLDNGVATDRQW
GTAAGAQLQQLKVLHSSQHSAFASPAKLEFRDEPGRLYRFSVHEPNMRKIEGTDQISHSYLMDTAARMEAQLHPKPV
GTSLHEVVAASVVAPRFCGDDLTSWLLREGTARTHAEAVMMVEYMIAAQVLYPVGMKRPQSRSLQFAYTAADLRTF
SADFEFSFVEPAKLARAAAYERMQRSVEQGKVARSQLLLRHTIQRRLAMEHHGDLGDDLYELLYAKPKPAEERRAEP
RHAVALAGDRHDXEGGASAGGDPPIPGDIDHPSSFDYPVVYRKVPGLLFEVAQEGVLTPEIRSGIEQAVRELEAHQVF
GITGDCGFMANYQHLVRSLASCPVFLSSLVLLPAISPLIGSRDRILCCTANGNSLAKLLPALWGAPPPPAAAYGVTHTF
THKHLDLHLLRPEQLVVCGFERVPGFDVVAEASSLMSHAVDRELVEQXDHVDGAHDPLARPHDQDDPARVHGAATL
HGDSAAGHRAARVRLTIDLQLRACGVRCKPIGGIAPLTLSWTPLPDQIPGYAHDWMIHDGQLMVVCKHSEPCCRPVL
QPFAHEGIRRDQHK
