## Supplemental dataset 3 for "Phylogeny and biogeography of the algal DMS-releasing enzyme"

**Dataset S3. Species with no DL homologs identified in available databases**

Search in available genomes:

Sacoglottis gabonensis

Dialium pachyphyllum

Porterandia cladantha

Zea mays (Corn)

Medicago sativa (Alfalfa)

Triticum aestivum (wheat)

Oryza sativa (rice)

Larrea tridentata (checked in SRA)

Alpinia zerumbet (Zingiberaceae) (checked in SRA)

Canna indica L. (Cannaceae) (checked in SRA)

Cissus sicyodes (L.) (Vitaceae) (checked in SRA)

Hibiscus rosa-sinensis L. (Malvaceae) (checked in SRA)

Inga vera W. (Fabaceae) (checked in SRA)

Mangifera indica L. (Anacardiaceae)

Pterocarpus indicus Wild. (Fabaceae) (checked in SRA)

Skeletonema menzellii (* in SRA there was one very partial read, though we wouldn't have counted it because it didn't have both cysteines - it was very short).

JGI Rhodophyta: TblastN against genome

Pyropia yezoensis U-51

Gracilariopsis chorda isolate SKKU-2015

Galdieria sulphuraria Azora

Galdieria sulphuraria MtSh

Galdieria sulphuraria SAG 21.92

Galdieria sulphuraria MS1

Galdieria sulphuraria YNP5578.1

Galdieria sulphuraria 5572

Galdieria sulphuraria 002

Cyanidioschyzon merolae Soos

Galdieria sulphuraria RT22

Galdieria phelgrea Soos

Chondrus crispus Stackhouse

Cyanidioschyzon merolae strain 10D

Galdieria sulphuraria 074W

Porphyra umbilicalis isolate 4086291

JGI blastp against predicted proteins (and tblastn vs genomes):

Chrysochromulina tobin CCMP291

Coccomyxa subellipsoidea C-169

Cryptophyceae sp. CCMP2293 v1.0

Cyanidioschyzon merolae Soos

Chlorella variabilis NC64A

Chlorokybus atmophyticus CCAC 0220

Chloropicon primus CCMP1205

Chondrus crispus Stackhouse

Chromochloris zofingiensis SAG 211-14

Chrysochromulina parva Lackey

Chlamydomonas reinhardtii v5.6

Chlamydomonas schloesseri CCAP 11/173

Chlorella sp. A99

Chlorella sorokiniana DOE1412

Chlorella sorokiniana UTEX 1230

Chlorella sorokiniana UTEX 1602

Chlorella sorokiniana str. 1228

Blastocystis hominis Singapore isolate B (sub-type 7)

Botryococcus braunii Showa v2.1

Caulerpa lentillifera

Chara braunii S276

Chlamydomonas eustigma NIES-2499

Chlamydomonas incerta SAG 7.73

Aurantiochytrium limacinum ATCC MYA-1381

Aureococcus anophagefferens clone 1984

Auxenochlorella protothecoides 0710

Auxenochlorella protothecoides UTEX 25

Bathycoccus prasinos RCC1105

Bigelowiella natans CCMP2755

Aplanochytrium kerguelense PBS07 v1.0

Arabidopsis lyrata

Asterochloris glomerata Cgr/DA1pho v2.0

Ochromonadaceae sp. CCMP2298 v1.0

Ochromonas sp. CCMP1393 v1.4

Ostreococcus lucimarinus

Ostreococcus sp. RCC809

Monoraphidium neglectum SAG 48.87

Naegleria gruberi v1.0

Nannochloropsis gaditana B-31

Nannochloropsis oceanica CCMP1779 v2.0

Nannochloropsis salina CCMP1776

Nemacystus decipiens Onna-1

Micractinium conductrix SAG 241.80

Micromonas commoda NOUM17 (RCC 299)

Micromonas pusilla CCMP1545

Minidiscus variabilis CCMP495 v1.0

Guillardia theta CCMP2712

Hyaloperonospora arabidopsidis Emoy2 v2.0

Klebsormidium nitens NIES-2285

Mesostigma viride CCAC 1140

Mesostigma viride NIES-296

Mesotaenium endlicherianum SAG 12.97

Galdieria sulphuraria MS1

Galdieria sulphuraria MtSh

Galdieria sulphuraria RT22

Galdieria sulphuraria SAG 21.92

Galdieria sulphuraria YNP5578.1

Giardia intestinalis ATCC 50803

Gonium pectorale NIES-2863

Gracilariopsis chorda isolate SKKU-2015

Fragilariopsis cylindrus CCMP 1102

Galdieria phelgrea Soos
Galdieria sulphuraria 002
Galdieria sulphuraria 074W
Galdieria sulphuraria 5572
Galdieria sulphuraria Azora
Enallax costatus CCAP 276/31 v1.0
Fistulifera solaris JPCC DA0580
Flechtneria rotunda SEV3-VF49 v1.0
Cyanidioschyzon merolae strain 10D
Cyanophora paradoxa CCMP329
Cyclotella cryptica CCMP322
Dunaliella salina CCAP19/18
Ectocarpus siliculosus Ec 32
Edaphochlamys debaryana CCAP 11/70
Scenedesmus obliquus var. UTEX2630 v1.0
Scenedesmus sp. NREL 46B-D3 v1.0
Schizochytrium aggregatum ATCC 28209
Scenedesmus obliquus UTEX B 3031
Scenedesmus obliquus var. DOE0013 v1.0
Scenedesmus obliquus var. UTEX 1450 v1.0
Pyropia yezoensis U-51
Raphidocelis subcapitata NIES-35
Reticulomyxa filosa
Saprolegnia parasitica CBS 223.65
Scenedesmus obliquus EN0004 v1.0
Scenedesmus obliquus UTEX 393
Picocystis sp. ML
Plasmodium falciparum 3D7
Populus trichocarpa v1.1
Porphyra umbilicalis isolate 4086291
Prasinoderma coloniale CCMP1413
Phytophthora capsici LT1534 v11.0
Phytophthora cinnamomi var cinnamomi v1.0

Phytophthora infestans T30-4

Phytophthora sojae v3.0

Picochlorum renovo

Picochlorum soloecismus DOE101

Phaeodactylum tricornutum CCAP 1055/1 v2.0

Physcomitrella patens subsp patens v1.1

Pavlovales sp. CCMP2436 v1.0

Ostreococcus tauri RCC1115 v1.0

Ostreococcus tauri RCC4221 v3.0

Oxytricha trifallax JRB310

Paramecium tetraurelia d4_2

Paraphysomonas imperforata CCMP1604 v1.4

Vitrella brassicaformis CCMP3155

Volvox carteri v2.1

Toxoplasma gondii ME49

Trebouxia sp. A1-2

Tribonema minus UTEX B ZZ1240 v1.0

Trichomonas vaginalis G3

Trypanosoma brucei brucei TREU927

Undaria pinnatifida M23

Tetradesmus obliquus UTEX B 72 v1.0

Tetrahymena thermophila SB210

Thalassiosira oceanica CCMP1005

Thalassiosira pseudonana CCMP 1335

Symbiochloris reticulata Switzerland extracted metagenome v1.0

Tetrabaena socialis NIES-571

Tetradesmus deserticola SNI-2 v1.0

Symbiochloris reticulata Africa extracted metagenome v1.0

Symbiochloris reticulata Scotland extracted metagenome v1.0

Symbiochloris reticulata Spain extracted metagenome v1.0

Symbiochloris reticulata Spain reference genome v1.0

Selaginella moellendorffii v1.0

Sorghum bicolor

all of phytozome (tblastn)

Zostera marina v3.1

Zostera marina v2.2

Zea mays RefGen_V4

Zea mays PHJ40 v1.2

Zea mays PHJ40 v1.1

Zea mays PHB47 v1.2

Zea mays PHB47 v1.1

Zea mays PH207 v1.1

Zea mays NKH8431 v1.2

Zea mays LH145 v1.2

Zea mays B84 v1.2

Volvox carteri v2.1

Vitis vinifera v2.1

Vigna unguiculata ZN016 v1.2

Vigna unguiculata v1.2

Vigna unguiculata v1.1

Vigna unguiculata UCR779 v1.1

Vigna unguiculata TZ30 v1.2

Vigna unguiculata Suvita2 v1.1

Vigna unguiculata Sanzi v1.1

Vigna unguiculata CB5-2 v1.1

Urochloa fusca v1.1

Triticum aestivum v2.2

Trifolium pratense v2

Thuja plicata v3.1

Thlaspi arvense v1.1

Thinopyrum intermedium v2.1

Theobroma cacao v2.1

Theobroma cacao v1.1

Stanleya pinnata v1.1

Spirodela polyrhiza v2

Spinacia oleracea Spov3

Sphagnum magellanicum v1.1

Sphagnum fallax v1.1

Sphagnum fallax v0.5

Sorghum bicolor v3.1.1

Sorghum bicolor SC187 v1.1

Sorghum bicolor RTx430 v2.1

Sorghum bicolor Rio v2.1

Sorghum bicolor BTx642 v1.1

Solanum tuberosum v6.1

Solanum tuberosum v4.03

Solanum lycopersicum ITAG4.0

Solanum lycopersicum ITAG3.2

Solanum lycopersicum ITAG2.4

Sinapis alba v3.1

Sinapis alba v1.1

Setaria viridis v2.1

Setaria viridis v1.1

Setaria italica v2.2

Selaginella moellendorffii v1.0

Schrenkiella parvula v2.2

Salix purpurea v5.1

Salix purpurea v1.0

Salix purpurea Fish Creek v3.1

Rorippa islandica v1.1

Ricinus communis v0.1

Quercus rubra v2.1

Prunus persica v2.1

Portulaca amilis v1.0

Porphyra umbilicalis v1.5

Populus trichocarpa v4.1

Populus trichocarpa v3.1

Populus trichocarpa v3.0

Populus trichocarpa Stettler14 v1.1

Populus maximowiczii x nigra NM6 v1.1

Populus deltoides WV94 v2.1

Poncirus trifoliata v1.3.1

Physcomitrium patens v3.3

Phaseolus vulgaris v2.1

Phaseolus vulgaris UI111 v1.1

Phaseolus vulgaris Labor Ovalle v1.1

Phaseolus vulgaris 5-593 v1.1

Phaseolus lunatus V1

Phaseolus acutifolius WLD v2.0

Phaseolus acutifolius v1.0

Pharus latifolius v1.1

Paspalum vaginatum v3.1

Panicum virgatum v5.1

Panicum virgatum v4.1

Panicum hallii v3.2

Panicum hallii v3.1

Panicum hallii v2.0

Panicum hallii HAL v2.2

Panicum hallii HAL v2.1

Ostreococcus lucimarinus v2.0

Oryza sativa v7.0

Oryza sativa Kitaake v3.1

Oropetium thomaeum v1.0

Olea europaea v1.0

Nymphaea colorata v1.2

Myagrum perfoliatum v2.1

Myagrum perfoliatum v1.1

Musa acuminata v1

Miscanthus sinensis v7.1

Mimulus guttatus v2.0

Mimulus guttatus TOL v5.0
Mimulus guttatus TOL v3.1
Mimulus guttatus NONTOL v4.0
Mimulus guttatus NONTOL v3.1
Micromonas sp RCC299 v3.0
Micromonas pusilla CCMP1545 v3.0
Medicago truncatula Mt4.0v1
Marchantia polymorpha v3.1
Manihot esculenta v8.1
Manihot esculenta v7.1
Manihot esculenta v6.1
Malus domestica v1.1
Malcolmia maritima v1.1
Lupinus albus v1
Lunaria annua v1.1
Lotus japonicus Lj1.0v1
Linum usitatissimum v1.0
Lindenbergia philippensis v1.1
Lepidium sativum v1.1
Lactuca sativa V8
Kalanchoe laxiflora v1.1
Kalanchoe fedtschenkoi v1.1
Joinvillea ascendens v1.1
Isatis tinctoria v1.1
Iberis amara v1.1
Hydrangea quercifolia v1.1
Hordeum vulgare r1
Helianthus annuus r1.2
Gossypium tomentosum v1.1
Gossypium raimondii v2.1
Gossypium mustelinum v1.1
Gossypium hirsutum v2.1
Gossypium hirsutum v1.1

Gossypium darwinii v1.1

Gossypium barbadense v1.1

Glycine soja v1.1

Glycine max Wm82.a4.v1

Glycine max Wm82.a2.v1

Glycine max Lee v1.1

Glycine max Fiskeby v1.1

Fragaria x ananassa v1.0.a1

Fragaria vesca v4.0.a2

Fragaria vesca v2.0.a2

Eutrema salsugineum v1.0

Euclidium syriacum v1.1

Eucalyptus grandis v2.0

Eruca vesicaria v1.1

Eleusine coracana v1.1

Dunaliella salina v1.0

Diptychocarpus strictus v2.1

Diptychocarpus strictus v1.1

Dioscorea alata v2.1

Dioscorea alata v1.1

Descurainia sophioides v1.1

Daucus carota v2.0

Cucumis sativus v1.0

Crambe hispanica v1.1

Corymbia citriodora v2.1

Coffea arabica v0.5

Coccomyxa subellipsoidea C-169 v2.0

Cleome violacea v2.1

Cleome violacea v1.1

Citrus sinensis v1.1

Citrus clementina v1.0

Cinnamomum kanehirae v3

Cicer arietinum v1.0

Chromochloris zofingiensis v5.2.3.2

Chlamydomonas reinhardtii v5.6

Chenopodium quinoa v1.0

Chasmanthium laxum v1.1

Ceratopteris richardii v2.1

Ceratodon purpureus R40 v1.1

Ceratodon purpureus GG1 v1.1

Caulanthus amplexicaulis v1.1

Castanea dentata v1.1

Carya illinoinensis v1.1

Carya illinoinensis Pawnee v1.1

Carya illinoinensis Lakota v1.1

Carya illinoinensis Elliott v1.1

Carica papaya ASGPBv0.4

Capsella rubella v1.1

Capsella grandiflora v1.1

Cakile maritima v1.1

Brassica rapa FPsc v1.3

Brassica oleracea capitata v1.0

Brachypodium sylvaticum v1.1

Brachypodium stacei v1.1

Brachypodium mexicanum v1.1

Brachypodium hybridum v1.1

Brachypodium hybridum Bhyb26 v2.1

Brachypodium distachyon v3.2

Brachypodium distachyon v3.1

Brachypodium distachyon v2.1

Brachypodium distachyon Uni2 v1

Brachypodium distachyon Tek-4 v1

Brachypodium distachyon Tek-2 v1

Brachypodium distachyon Sig2 v1

Brachypodium distachyon S8iic v1

Brachypodium distachyon Ron2 v1

Brachypodium distachyon Per1 v1

Brachypodium distachyon Pangenome v1

Brachypodium distachyon Mur1 v1

Brachypodium distachyon Mon3 v1

Brachypodium distachyon Mig3 v1

Brachypodium distachyon Luc1 v1

Brachypodium distachyon Koz-3 v1

Brachypodium distachyon Koz-1 v1

Brachypodium distachyon Kah-5 v1

Brachypodium distachyon Kah-1 v1

Brachypodium distachyon Jer1 v1

Brachypodium distachyon Gaz-8 v1

Brachypodium distachyon Foz1 v1

Brachypodium distachyon Bis-1 v1

Brachypodium distachyon BdTR13c v1

Brachypodium distachyon BdTR13a v1

Brachypodium distachyon BdTR12c v1

Brachypodium distachyon BdTR11i v1

Brachypodium distachyon BdTR11g v1

Brachypodium distachyon BdTR11a v1

Brachypodium distachyon BdTR10c v1

Brachypodium distachyon BdTR9k v1

Brachypodium distachyon BdTR8i v1

Brachypodium distachyon BdTR7a v1

Brachypodium distachyon BdTR5i v1

Brachypodium distachyon BdTR3c v1

Brachypodium distachyon BdTR2g v1

Brachypodium distachyon BdTR2b v1

Brachypodium distachyon BdTR1i v1

Brachypodium distachyon Bd30-1 v1.1

Brachypodium distachyon Bd30-1 v1

Brachypodium distachyon Bd29-1 v1

Brachypodium distachyon Bd21-3 v1.2

Brachypodium distachyon Bd21-3 v1.1

Brachypodium distachyon Bd21-3 v1

Brachypodium distachyon Bd21 AsmbCtrl v1

Brachypodium distachyon Bd21 AnntCtrl v1

Brachypodium distachyon Bd18-1 v1

Brachypodium distachyon Bd3-1 v1

Brachypodium distachyon Bd2-3 v1

Brachypodium distachyon Bd1-1 v1.1

Brachypodium distachyon Bd1-1 v1

Brachypodium distachyon Arn1 v1

Brachypodium distachyon Adi-12 v1

Brachypodium distachyon Adi-10 v1

Brachypodium distachyon Adi-2 v1

Brachypodium distachyon ABR9 v1

Brachypodium distachyon ABR8 v1

Brachypodium distachyon ABR7 v1

Brachypodium distachyon ABR6 v1

Brachypodium distachyon ABR5 v1

Brachypodium distachyon ABR4 v1

Brachypodium distachyon ABR3 v1

Brachypodium distachyon ABR2 v1

Botryococcus braunii v2.1

Boechera stricta v1.2

Betula platyphylla v1.1

Beta vulgaris EL10_1.0

Asparagus officinalis V1.1

Arachis hypogaea v1.0

Arabidopsis thaliana TAIR10

Arabidopsis thaliana Araport11

Arabidopsis lyrata v2.1

Arabidopsis halleri v1.1

Aquilegia coerulea v3.1

Ananas comosus v3

Anacardium occidentale v0.9

Amborella trichopoda v1.0

Amaranthus hypochondriacus v2.1

Alyssum linifolium v1.1

Acorus americanus v1.1

rutgers rhodophyta (tblastn)

Compsopogon_coeruleusSAG36.94MMETSP0312.RNAseqAssem.fa;

Cmer_Soos.gmc.fa; Cmer_Soos.est.fa;

Erythrolobus_australicusCCMP3124MMETSP1353.RNAseqAssem.fa;

ErythrolobusmadagascarensisCCMP3276MMETSP1354.RNAseqAssem.fa;

eukaryoteOtherV1.4.fa; Gphleg_Soos.gmc.fa; Gphleg_Soos.est.fa;

Gsulp_002.gmc.fa; Gsulp_002.est.fa; Gsulp_5572.gmc.fa;

Gsulp_5572.est.fa; Gsulp_Azora.gmc.fa; Gsulp_Azora.est.fa;

Gsulp_MS1.gmc.fa; Gsulp_MS1.est.fa; Gsulp_MtSh.gmc.fa;

Gsulp_MtSh.est.fa; Gsulp_RT22.gmc.fa; Gsulp_RT22.est.fa;

Gsulp_SAG21.gmc.fa; Gsulp_SAG21.est.fa; Gsulp_YNP5587_1.gmc.fa;

Gsulp_YNP5587_1.est.fa; Gracilariopsis_chorda.fa;

Madagascaria_erythrocladiodesCCMP3234MMETSP1450.RNAseqAssem.fa;

Porphyra.454ESTs.CAP3_Dec2010.fa;

Porphyra.all.V1.4.fixedIndels.fa;

Porphyra.edgeRNASeq.Trinity.NOV2012.fa;

Porphyra_haitanensis_v1_CDS.fasta;

Porphyra_haitanensis_v1_genome.fasta;

Porphyra_purpurea_chloroplast.GB.fa;

Porphyra_umbilicalis_mitochondrion.GB.fa;

porphyraV1.3.1ff.eukContam.bin.fa; porphyraV1.3.1ff.mainGenome.bin.fa;

porphyraV1.3.1ff.prokContam.bin.fa; porphyraV1.3.1ff.unknown.bin.fa;

porphyraV1.4.fa;

porphyraV1.5.fa;

Porphyra_yezoensis_v1_CDS.fasta;

Porphyra_yezoensis_v1_genome.fasta;

Porphyridium_aerugineumSAG1380-2MMETSP0313.RNAseqAssem.fa;

Pumbilicalis_plastid_cds.KeelingLab.fa;

Pyezoensis_CLC_genomic_scaffolds_v1.fa;

Rhodella_maculataCCMP736MMETSP0167.RNAseqAssem.fa;

Rhodella_maculataCCMP736MMETSP0314.RNAseqAssem.fa;

Rhodosorus_marinusMMETSP0011.RNAseqAssem.fa;

Rhodosorus_marinusUTEXLB2760MMETSP0315.RNAseqAssem.fa;

Timspurckia_oligopyrenoidesCCMP3278MMETSP1172.RNAseqAssem.fa

Genome in NCBI, TBLASTN:

Fragilaria radians (Synedra acus subsp. Radians)

Isochrysis galbana
