## Supplemental information for "Phylogeny and biogeography of the algal DMS-releasing enzyme"

Supplementary Information

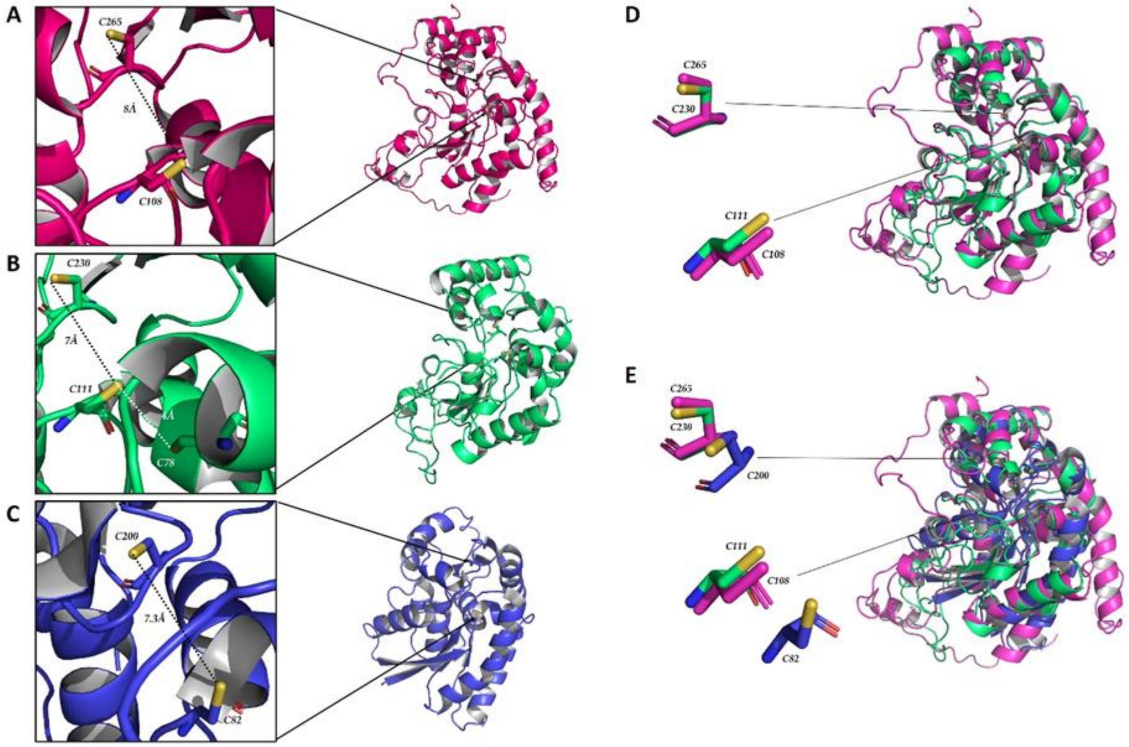


**Fig. S1.** Ribbon representation of the Alma1, Sym-Alma and bacterial Maleate Isomerase protein structures. (A-C) Predicted structure of two eukaryotic DMSP lyase proteins, Alma1 from *E. huxleyi* (A), and Sym-Alma from *Symbiodinium* A1 (B), as well as the known structure of bacterial Maleate Isomerase Iso (PDB:4FQ7) from *Pseudomonas putida* S16 (C). Insets show zoom-in view of the active site area in the DMSP lyases, where the canonical cysteines are shown as sticks: Cys108 and Cys265 for Alma1 (A) and Cys111 and Cys230 for Sym-Alma (B). (D-E) Structural alignment of Alma1 (pink) and Sym-Alma (green) (D) and Alma1, Sym-Alma and 4FQ7 (blue, E). Note that the cysteines residues of the eukaryotic homologs align perfectly, while the bacterial cysteines are not placed in the same position.

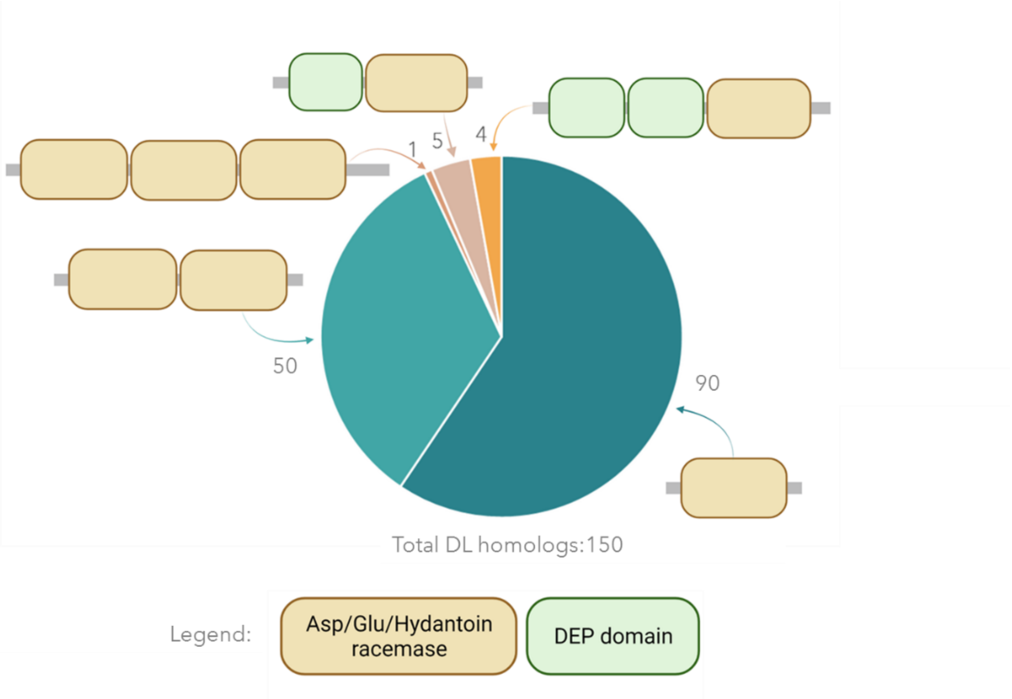


**Fig. S2. Conserved domain organization of eukaryotic DL homologs.** A schematic description of the different conserved domains (according to the CDD database) identified in a total of 150 DL homologs (see also dataset S1).

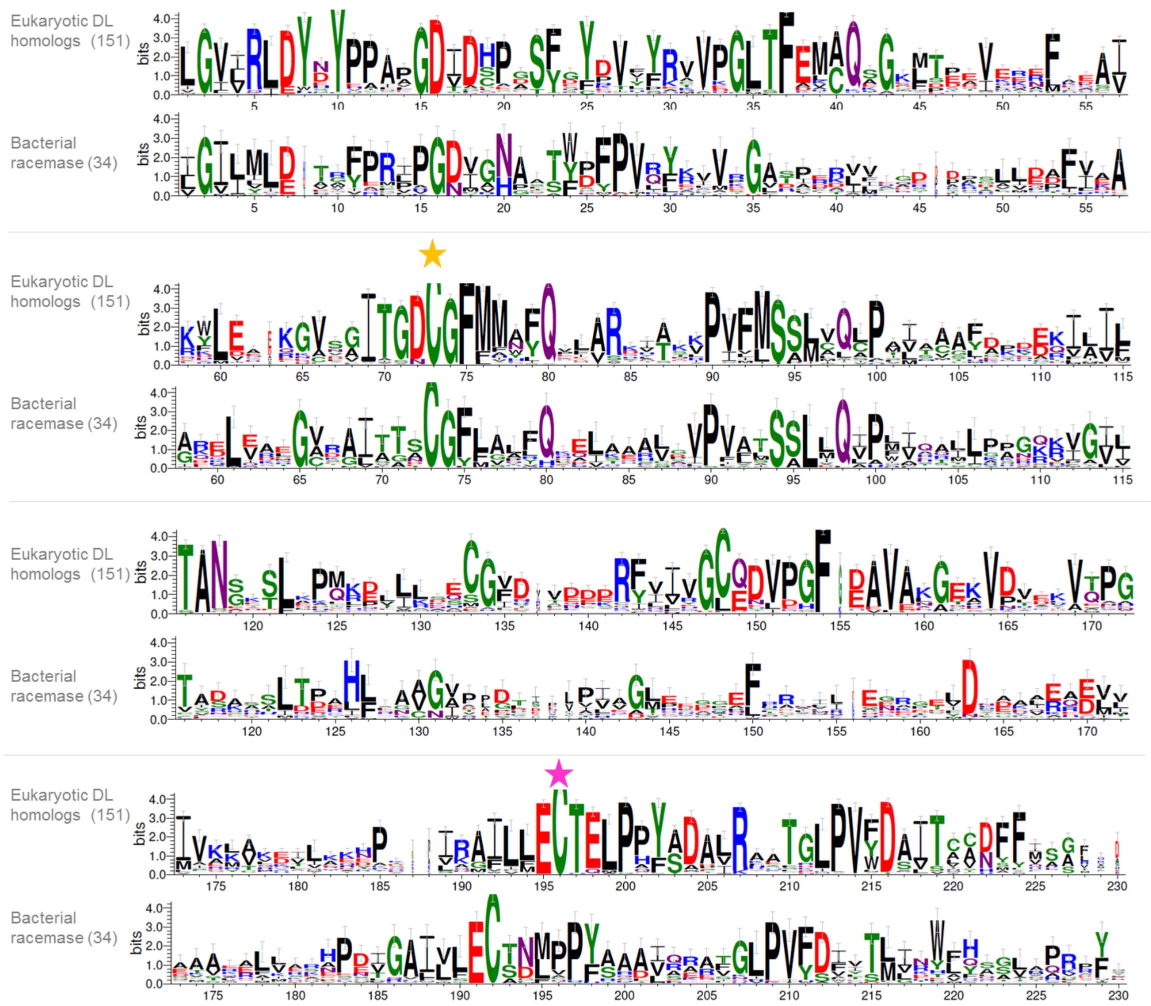


**Fig. S3. Sequence logos of multiple sequence alignment of the racemase domain from bacteria and eukaryotic DL homologs.** A total of 151 eukaryotic DL homologs were aligned (see dataset S1), and 34 bacterial sequences belong the CDD subfamily PRK0747 (which is the closest to the Asp/Glu/Hydantoin racemase domain). Stars designate the first (yellow) and second (pink) canonical cysteine residues located at the active site of the DL enzyme.

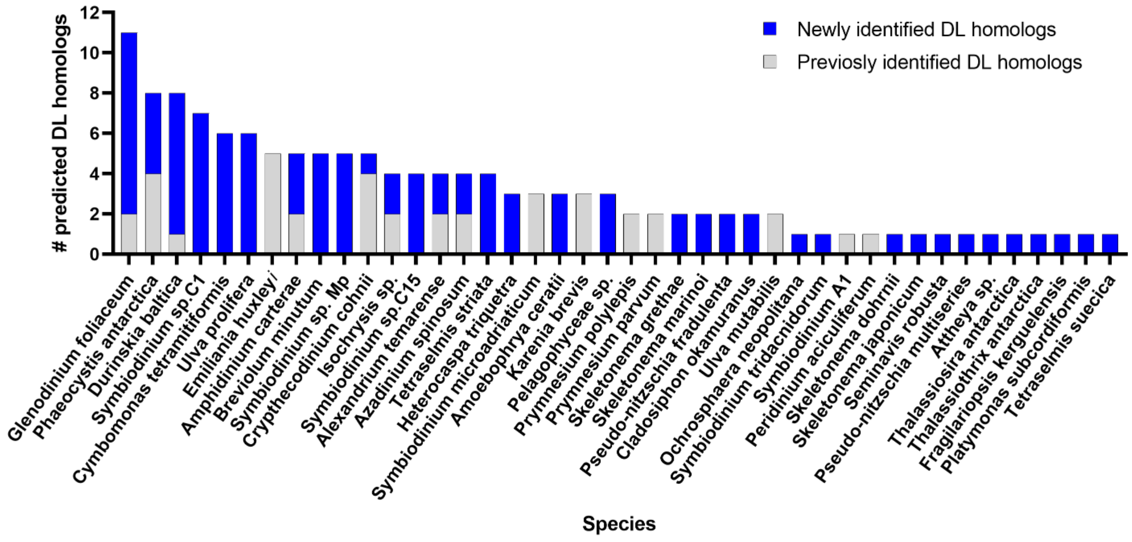


**Fig. S4. Predicted DL homologs in different phytoplankton species.** The number of predicted DL homologs is plotted per species.

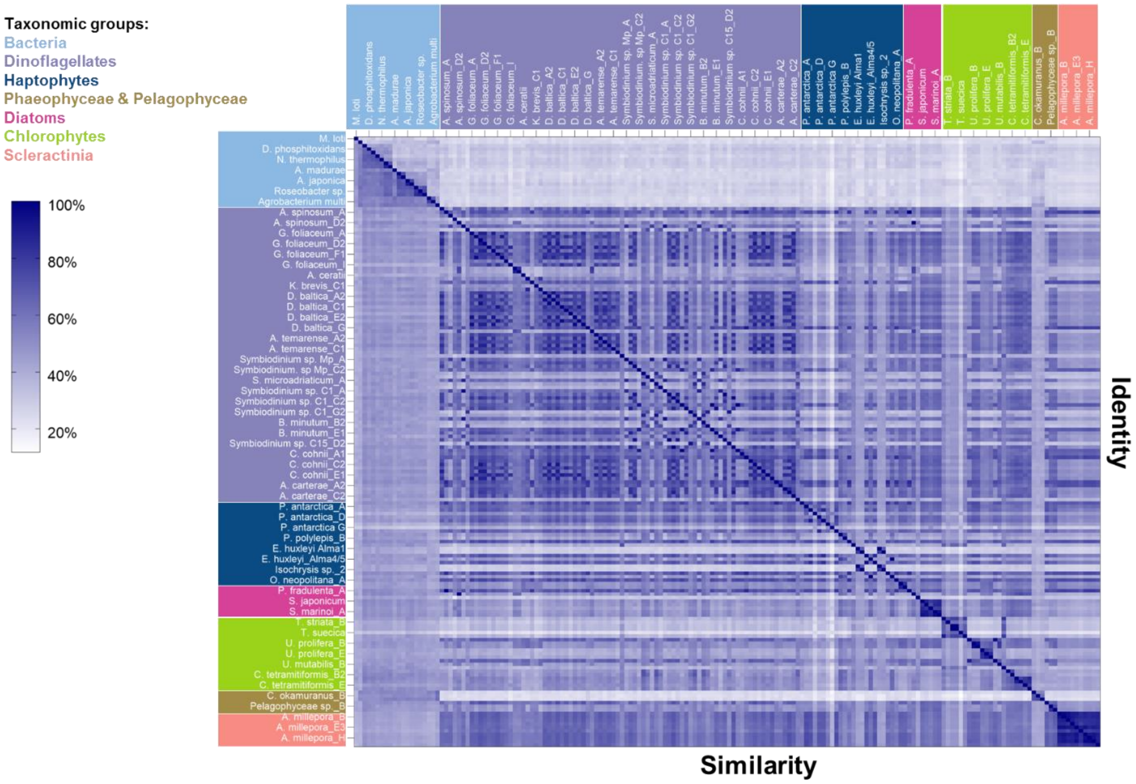


**Fig. S5. Sequence similarity and identity of bacterial and eukaryotic DL homologs.** The plot shows the sequence similarity scores below the diagonal and sequence identity scores above, for multiple sequence alignment of racemase domain of 174 DL homologs. Each row and column represent a DL homolog (see dataset S1). Due to space limitation, only third of the names could be presented. The sequences are orders according to their taxonomy (see legend).

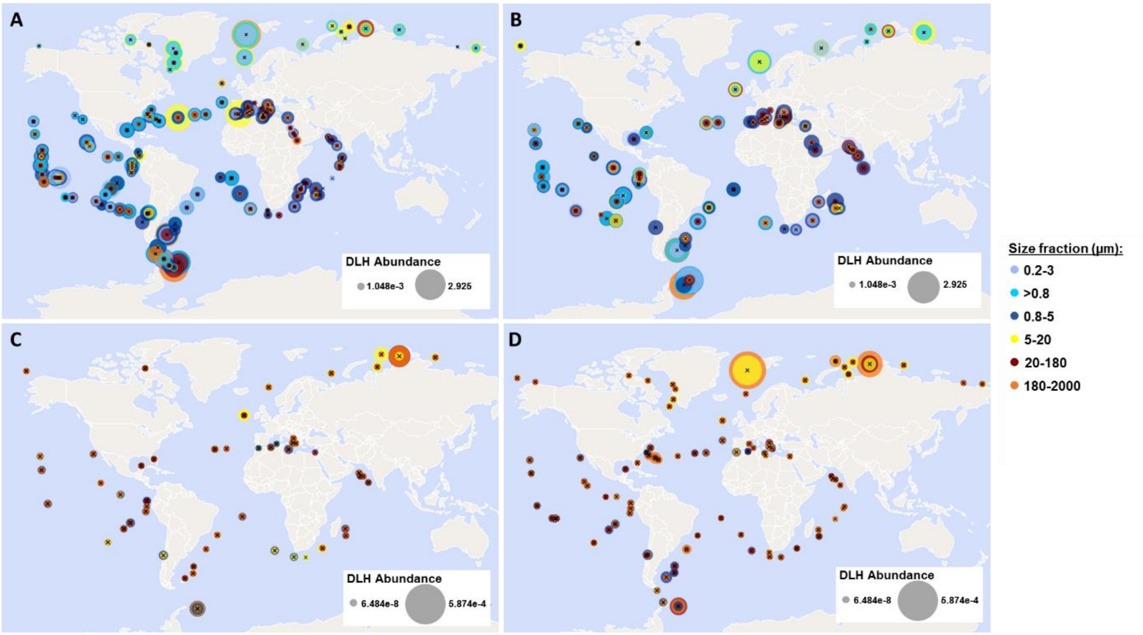


**Fig. S6. Abundance of DMSP lyase homologs (DLHs) in the oceans.** Geographic distribution of DLHs in phytoplankton found in *Tara* Oceans metagenomes (A,B) and metatranscriptomes (C,D) datasets. Color depicts the size fraction of each sample. The circle size is proportional to the relative abundance of DLH genes or transcripts, which was normalized as percent of mapped reads.

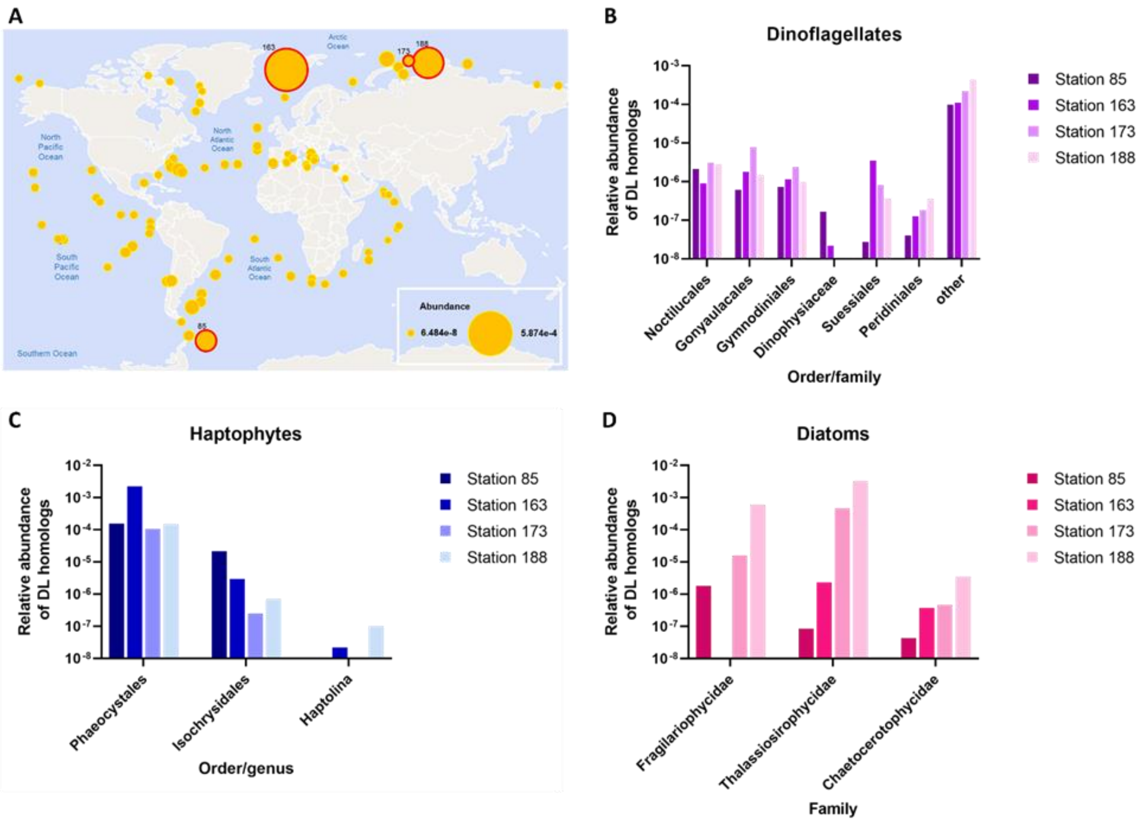


**Fig. S7. Taxonomy of phytoplankton in selected *Tara* station with high DL-homologs expression.** (A). Location of stations *Tara* 85, 163, 173 and 188. The circle size is proportional to the relative abundance of DLH genes or transcripts, which was normalized as percent of mapped reads. (B) Identified dinoflagellates orders/families in each station. (C) Identified haptophytes orders/genus in each station. (D) Identified diatoms families in each station.

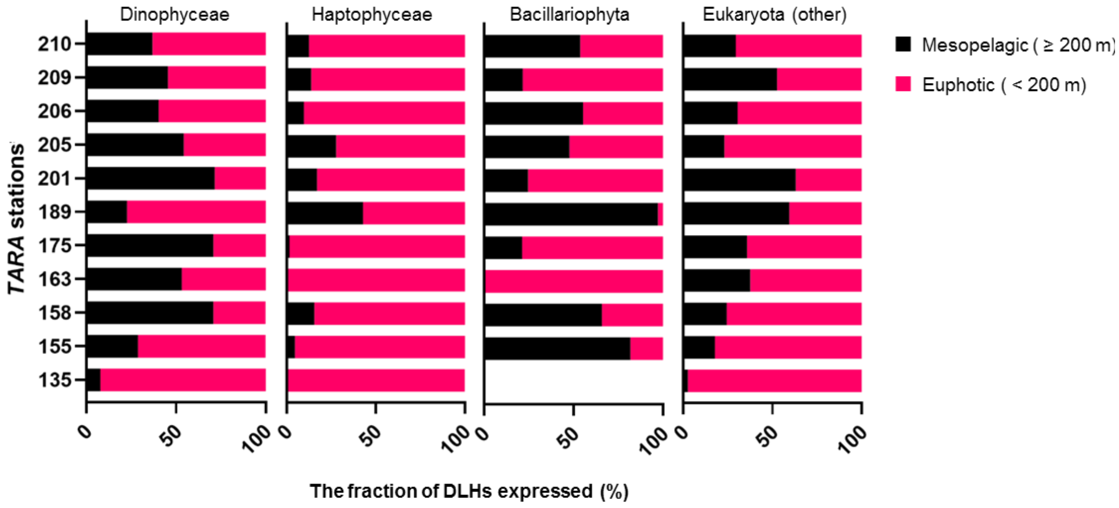


**Fig. S8. Expression of DLHs in the euphotic versus mesopelagic zones in the *Tara* Ocean dataset.** DLHs (DMSP lyase homologs) were detected in mesopelagic depths (200-751 m) in 11 stations (Y axis). The fraction of DLH transcripts expressed in the mesopelagic (black) and euphotic (pink) zone is presented for the main DLHs-expressing taxa.

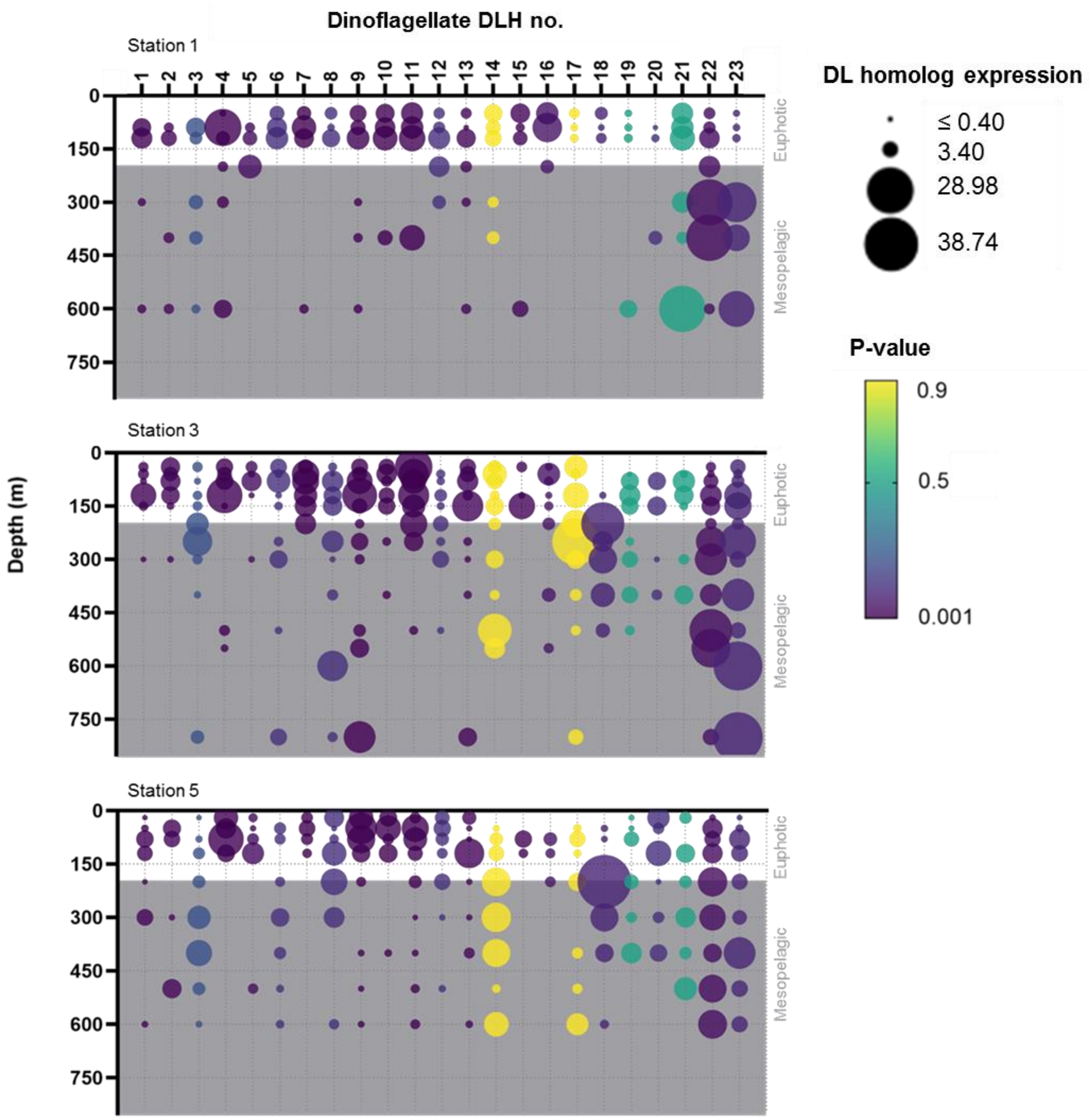


**Fig. S9. Dinoflagellate DLHs are differentially expressed in euphotic and mesopelagic depths in the central Pacific Ocean.** The expression level of 23 DLHs from natural dinoflagellates population is presented for three different locations (stations 1, 3, 5) in the central Pacific Ocean. Each station contains a depth profile. The circle size represents the normalized expression level. The color depicts the P - value, showing how significantly different is the expression between the euphotic (white background) and the mesopelagic (grey background) zones. The expression data was taken from Cohen NR, et al. (2021).

### Table S1. Conserved motifs in the DL protein sequence.

| Motif | P-value | E-value | Sites | Training set-positives | Training set-negatives | Score | Testing set-positives | Testing set-negatives | P-value | Match Threshold |
|---|---|---|---|---|---|---|---|---|---|---|
| GCZDVPGFD | 1.40E-16 | 4.10E-15 | 145 (96.0%) | 131 / 136 (96.3%) | 0 / 826 (0.0%) | 5.20E-157 | 14 / 15 (93.3%) | 0 / 91 (0.0%) | 1.40E-16 | 7.54392 |
| GDCGFMMAFQ | 6.60E-15 | 1.90E-13 | 149 (98.7%) | 136 / 136 (100.0%) | 0 / 826 (0.0%) | 1.60E-169 | 13 / 15 (86.7%) | 0 / 91 (0.0%) | 6.60E-15 | 25.5612 |
| IRAILLECTEL | 6.60E-15 | 1.90E-13 | 148 (98.0%) | 135 / 136 (99.3%) | 0 / 826 (0.0%) | 1.30E-166 | 13 / 15 (86.7%) | 0 / 91 (0.0%) | 6.60E-15 | 25.6013 |
| EAKGVSGIT | 6.60E-15 | 1.90E-13 | 126 (83.4%) | 113 / 136 (83.1%) | 3 / 826 (0.4%) | 2.50E-119 | 13 / 15 (86.7%) | 0 / 91 (0.0%) | 6.60E-15 | 11.3761 |
| RVVPGLTFEMAQSG | 2.10E-13 | 6.00E-12 | 144 (95.4%) | 132 / 136 (97.1%) | 0 / 826 (0.0%) | 3.20E-159 | 12 / 15 (80.0%) | 0 / 91 (0.0%) | 2.10E-13 | 16.3706 |
| GVIRLDYBY | 2.10E-13 | 6.00E-12 | 143 (94.7%) | 131 / 136 (96.3%) | 0 / 826 (0.0%) | 5.20E-157 | 12 / 15 (80.0%) | 0 / 91 (0.0%) | 2.10E-13 | 15.5877 |
| RAATGLPVFDAI | 2.10E-13 | 6.00E-12 | 143 (94.7%) | 131 / 136 (96.3%) | 0 / 826 (0.0%) | 5.20E-157 | 12 / 15 (80.0%) | 0 / 91 (0.0%) | 2.10E-13 | 20.2413 |
| GDIDHPGSFGY | 2.10E-13 | 6.00E-12 | 144 (95.4%) | 132 / 136 (97.1%) | 2 / 826 (0.2%) | 2.80E-155 | 12 / 15 (80.0%) | 0 / 91 (0.0%) | 2.10E-13 | 21.5522 |
| ILTANSKSLKP | 6.70E-13 | 1.90E-11 | 147 (97.4%) | 134 / 136 (98.5%) | 0 / 826 (0.0%) | 5.50E-164 | 13 / 15 (86.7%) | 2 / 91 (2.2%) | 6.70E-13 | 16.4622 |
| PPYADAL | 6.70E-13 | 1.90E-11 | 132 (87.4%) | 119 / 136 (87.5%) | 10 / 826 (1.2%) | 4.20E-120 | 13 / 15 (86.7%) | 2 / 91 (2.2%) | 6.70E-13 | 13.6881 |

**Table S2. DL homologs identified by Alcolombri et al., (2015).** The WP accessions are from NCBI and the CAMPEP accessions are from the MMETSP databases.

| Species | Domain | Phylum | Name in tree- Fig. 4a from Alcolombri et al., 2015 | Accession number |
|---|---|---|---|---|
| Paraburkholderia ferrariae | Bacteria | Proteobacteria | P. ferrariae | WP_028225891.1 |
| Streptosporangium roseum | Bacteria | Actinobacteria | S. roseum | WP_051865922.1 |
| Robbsia andropogonis | Bacteria | Proteobacteria | R. andropongonis | WP_024904647.1 |
| Geopsychrobacter electrodiphilus | Bacteria | Proteobacteria | G. electrodiphilus | WP_020674609.1 |
| Azospirillum lipoferum | Bacteria | Proteobacteria | A. lipoferum | WP_042444357.1 |
| Pseudovibrio sp. FO-BEG1 | Bacteria | Proteobacteria | Pseudovibrio sp. | WP_014285483.1 |
| Micromonospora sp. ATCC 39149 | Bacteria | Actinomycetia | Micromonospora sp. | WP_007071402.1 |
| Agrobacterium (Multispecies) | Bacteria | Proteobacteria | Agrobacterium multi | WP_007689264.1 |
| Mesorhizobium loti | Bacteria | Proteobacteria | M. loti | WP_019863393.1 |
| Actinomadura madurae | Bacteria | Actinobacteria | A. madurae | WP_063747721.1 |
| Amphritea japonica | Bacteria | Proteobacteria | A. japonica | WP_019621467.1 |
| Desulfotignum phosphitoxidans | Bacteria | Proteobacteria | D. phosphitoxidans | WP_006965703.1 |
| Roseobacter sp. SK209-2-6 | Bacteria | Proteobacteria | Roseobacter sp. | WP_008209391.1 |
| unclassified Leisingera (Multispecies) | Bacteria | Proteobacteria | Leisingera multi | WP_019296982.1 |
| Alkalihalobacillus wakoensis | Bacteria | Firmicutes | A. wakoensis | WP_034745099.1 |
| Natranaerobius thermophilus | Bacteria | Firmicutes | N. thermophilus | WP_012448288.1 |
| Desulfatirhabdium butyrativorans | Bacteria | Proteobacteria | D. butyrativorans | WP_028324949.1 |
| Acetomicrobium hydrogeniformans | Bacteria | Synergistetes | A. hydrogeniformans | WP_009200906.1 |
| Neptuniibacter caesariensis | Bacteria | Proteobacteria | N. caesariensis | WP_007022653.1 |
| Agrobacterium rhizogenes | Bacteria | Proteobacteria | A. rhizogenes | WP_047469519.1 |
| Candidatus Puniceispirillum marinum | Bacteria | Proteobacteria | C.P. marinum | WP_013045355.1 |
| Crypthecodinium cohnii (Seligo) | Eukaryota | Dinoflagellata | C. cohnii A | CAMPEP_0193854358 |
| Crypthecodinium cohnii (Seligo) | Eukaryota | Dinoflagellata | C. cohnii B | CAMPEP_0193926808 |
| Crypthecodinium cohnii (Seligo) | Eukaryota | Dinoflagellata | C. cohnii C | CAMPEP_0193865556 |
| Crypthecodinium cohnii (Seligo) | Eukaryota | Dinoflagellata | C. cohnii D | CAMPEP_0193933962 |
| Azadinium spinosum 3D9 | Eukaryota | Dinoflagellata | A. spinosum A | CAMPEP_0186837182 |
| Azadinium spinosum 3D9 | Eukaryota | Dinoflagellata | A. spinosum B | CAMPEP_0186768496 |
| Karenia brevis CCMP2299 | Eukaryota | Dinoflagellata | K. brevis A | CAMPEP_0188921804 |
| Karenia brevis CCMP2299 | Eukaryota | Dinoflagellata | K. brevis B | CAMPEP_0188855392 |
| Karenia brevis CCMP2299 | Eukaryota | Dinoflagellata | K. brevis C | CAMPEP_0188922230 |
| Alexandrium temarense CCMP1771 | Eukaryota | Dinoflagellata | A. temarense A | CAMPEP_0186397762 |
| Alexandrium temarense CCMP1771 | Eukaryota | Dinoflagellata | A. temarense B | CAMPEP_0186178466 |
| Peridinium aciculiferum PAER_2 | Eukaryota | Dinoflagellata | P. aciculiferum | CAMPEP_0190608078 |
| Durinskia baltica CSIRO_CS-38 | Eukaryota | Dinoflagellata | D. baltica | CAMPEP_0200062214 |
| Amphidinium carterae CCMP1314 | Eukaryota | Dinoflagellata | A. carterae A | CAMPEP_0186439658 |
| Amphidinium carterae CCMP1314 | Eukaryota | Dinoflagellata | A. carterae B | CAMPEP_0186460378 |
| Glenodinium foliaceum CCAP1116 | Eukaryota | Dinoflagellata | G. foliaceum A | CAMPEP_0188240084 |
| Glenodinium foliaceum CCAP1116 | Eukaryota | Dinoflagellata | G. foliaceum B | CAMPEP_0188434408 |
| Isochrysis sp. | Eukaryota | Haptophyta | Isochrysis sp. A | CAMPEP_0188750792 |
| Isochrysis sp. | Eukaryota | Haptophyta | Isochrysis sp. B | CAMPEP_0188739558 |
| Prymnesium parvum Texoma | Eukaryota | Haptophyta | P. parvum A | CAMPEP_0191203590 |
| Prymnesium parvum Texoma | Eukaryota | Haptophyta | P. parvum B | CAMPEP_0191252250 |

### Table S3. Confirmation of DL homologs expression in the MMETSP database.

| Species | MMETSP Sample IDs | Contigs | Expression ? |
| --- | --- | --- | --- |
| *P. aciculiferum* | MMETSP0370, MMETSP0371 | Peridinium-aciculiferum-PAER_2-20130926\|2150_0 | yes |
| *G. foliaceum* | MMETSP0118_2, MMETSP0119_2 | Glenodinium-foliaceum-CCAP1116_3-20130913\|6335_1 | yes |
| | | Glenodinium-foliaceum-CCAP1116_3-20130913\|269394_1 | yes |
| | | Glenodinium-foliaceum-CCAP1116_3-20130913\|39119_1 | yes |
| | | Glenodinium-foliaceum-CCAP1116_3-20130913\|264918_1 | yes |
| | | Glenodinium-foliaceum-CCAP1116_3-20130913\|265187_1 | yes |
| | | Glenodinium-foliaceum-CCAP1116_3-20130913\|162165_1 | yes |
| | | Glenodinium-foliaceum-CCAP1116_3-20130913\|11173_1 | yes |
| | | Glenodinium-foliaceum-CCAP1116_3-20130913\|144842_1 | yes |
| *S. kawagutii* | MMETSP0132_2C, MMETSP0133_2, MMETSP0134_2, MMETSP0135_2 | Symbiodinium-kawagutii-CCMP2468-20131203\|3329_1 | no |
| *Symbiodinium-sp-C15* | MMETSP1370, MMETSP1371 | Symbiodinium-sp-C15-20130923\|6074_1 | yes |
| | | Symbiodinium-sp-C15-20130923\|119450_1 | yes |
| | | Symbiodinium-sp-C15-20130923\|60114_1 | yes |
| | | Symbiodinium-sp-C15-20130923\|18175_1 | yes |
| | | Symbiodinium-sp-C15-20130923\|25623_1 | yes |
| | | Symbiodinium-sp-C15-20130923\|18047_1 | yes |
| *Symbiodinium-sp-Mp* | MMETSP1122, MMETSP1123, MMETSP1124, MMETSP1125 | Symbiodinium-sp-Mp-20130822\|28546_1 | yes |
| | | Symbiodinium-sp-Mp-20130822\|90026_1 | yes |
| | | Symbiodinium-sp-Mp-20130822\|7847_1 | yes |
| | | Symbiodinium-sp-Mp-20130822\|75336_1 | yes |
| | | Symbiodinium-sp-Mp-20130822\|189624_1 | yes |
| *Symbiodinium-sp-C1* | MMETSP1367, MMETSP1369 | Symbiodinium-sp-C1-20140214\|19746_1 | yes |
| | | Symbiodinium-sp-C1-20140214\|27762_1 | yes |
| | | Symbiodinium-sp-C1-20140214\|794_1 | yes |
| | | Symbiodinium-sp-C1-20140214\|25073_1 | yes |
| | | Symbiodinium-sp-C1-20140214\|27039_1 | yes |
| | | Symbiodinium-sp-C1-20140214\|22040_1 | yes |
| *K. brevis* | MMETSP0027, MMETSP0029, MMETSP0030, | Karenia-brevis-CCMP2229-20130916\|56488_1 | yes |
| | | Karenia-brevis-CCMP2229-20130916\|7473_1 | yes |
| | | Karenia-brevis-CCMP2229-20130916\|56810_1 | yes |
| *A. carterae* | MMETSP0398C, MMETSP0399, MMETSP0258, MMETSP0259 | Amphidinium-carterae-CCMP1314-20130924\|22079_1 | yes |
| | | Amphidinium-carterae-CCMP1314-20130924\|63059_1 | yes, except MMETSP0399 |
| | | Amphidinium-carterae-CCMP1314-20130924\|57390_1 | only in MMETSP0258, MMETSP0259 |
| | | Amphidinium-carterae-CCMP1314-20130924\|23864_1 | yes |
| | | Amphidinium-carterae-CCMP1314-20130924\|16687_1 | only in MMETSP0258, MMETSP0259 |
| *C. cohnii* | MMETSP0323_2, MMETSP0324_2, MMETSP0325_2, MMETSP0326_2 | Crypthecodinium-cohnii-Seligo-20130904\|11054_1 | yes |
| | | Crypthecodinium-cohnii-Seligo-20130904\|174461_1 | yes |
| | | Crypthecodinium-cohnii-Seligo-20130904\|19770_1 | yes |
| | | Crypthecodinium-cohnii-Seligo-20130904\|195906_1 | yes, except MMETSP0323_2 |
| | | Crypthecodinium-cohnii-Seligo-20130904\|20414_1 | yes |
| *A. spinosum* | MMETSP1036_2, MMETSP1037_2, MMETSP1038_2 | Azadinium-spinosum-3D9-20130829\|183522_1 | yes |
| | | Azadinium-spinosum-3D9-20130829\|29318_1 | yes |
| | | Azadinium-spinosum-3D9-20130829\|17069_1 | yes |
| | | Azadinium-spinosum-3D9-20130829\|21942_1 | yes |
| *D. baltica* | MMETSP0116_2, MMETSP0117_2 | Durinskia-baltica-CSIRO_CS-38-20140214\|203657_1 | no |
| | | Durinskia-baltica-CSIRO_CS-38-20140214\|157896_1 | yes |
| | | Durinskia-baltica-CSIRO_CS-38-20140214\|5581_1 | yes |
| | | Durinskia-baltica-CSIRO_CS-38-20140214\|164224_1 | yes |
| | | Durinskia-baltica-CSIRO_CS-38-20140214\|160103_1 | yes |
| | | Durinskia-baltica-CSIRO_CS-38-20140214\|157677_1 | yes |
| | | Durinskia-baltica-CSIRO_CS-38-20140214\|238428_1 | yes |
| | | Durinskia-baltica-CSIRO_CS-38-20140214\|230672_1 | yes |
| | | Durinskia-baltica-CSIRO_CS-38-20140214\|6229_1 | yes |
| | | Durinskia-baltica-CSIRO_CS-38-20140214\|209311_1 | only in MMETSP0117_2 |
| | | Durinskia-baltica-CSIRO_CS-38-20140214\|151706_1 | yes |
| | | Durinskia-baltica-CSIRO_CS-38-20140214\|238089_1 | yes |
| | | Durinskia-baltica-CSIRO_CS-38-20140214\|14482_1 | yes |
| *A. temarense* | MMETSP0378, MMETSP0380, MMETSP0382, MMETSP0384 | Alexandrium-temarense-CCMP1771-20130823\|410328_1 | yes |
| | | Alexandrium-temarense-CCMP1771-20130823\|2738_1 | yes |
| | | Alexandrium-temarense-CCMP1771-20130823\|37484_1 | yes |
| | | Alexandrium-temarense-CCMP1771-20130823\|165856_1 | yes |

### Table S3- continue. Confirmation of DL homologs expression in the MMETSP database.

| | | | |
|---|---|---|---|
| *Isochrysis sp.* | MMETSP1388, MMETSP1090 | Isochrysis-sp-CCMP1244-20130912\|8451_1 | yes |
| | | Isochrysis-sp-CCMP1244-20130912\|1392_1 | yes |
| | | Isochrysis-sp-CCMP1244-20130912\|2246_1 | yes |
| | | Isochrysis-sp-CCMP1244-20130912\|23305_1 | yes |
| *E. huxleyi* CCMP370 | MMETSP1154, MMETSP1155, MMETSP1156, MMETSP1157 | Emiliania-huxleyi-CCMP370-20130905\|29607_1 | yes |
| *E. huxleyi* PLYM219 | MMETSP1150, MMETSP1151, MMETSP1152, MMETSP1153 | Emiliania-huxleyi-PLYM219-20130905\|8612_1 | yes |
| | | Emiliania-huxleyi-PLYM219-20130905\|91030_1 | yes |
| *E. huxleyi* CCMP379 | MMETSP0994, MMETSP0995, MMETSP0996, MMETSP0997 | Emiliania-huxleyi-379-20130905\|8113_1 | yes |
| | | Emiliania-huxleyi-379-20130905\|18426_1 | yes |
| *C. polylepis* | MMETSP0143, MMETSP0145, | Chrysochromulina-polylepis-CCMP1757-20130903\|18714_1 | yes |
| *P. parvum texoma* | MMETSP0006_2, MMETSP0007, MMETSP0008_2, MMETSP0814, | Prymnesium-parvum-Texoma1-20131001\|4355_1 | yes |
| | | Prymnesium-parvum-Texoma1-20131001\|104153_1 | yes |
| | | Prymnesium-parvum-Texoma1-20131001\|200449_1 | yes |
| | | Prymnesium-parvum-Texoma1-20131001\|21691_1 | yes |
| *S. marinoi* | MMETSP0918, MMETSP0920 | Skeletonema-marinoi-SkelA-20130924\|7776_1 | yes |
| *S. dohrnii* | MMETSP0562, MMETSP0563 | Skeletonema-dohrnii-SkelB-20130926\|18976_1 | yes |
| *P. fradulenta** | MMETSP0850, MMETSP0851, | Pseudo_nitzschia-fradulenta-WWA7-20140214\|209615_1 | only in MMETSP0851 |
| | | Pseudo_nitzschia-fradulenta-WWA7-20140214\|96008_1 | only in MMETSP0851 |
| *T.antarctica** | MMETSP0902, MMETSP0903, | Thalassiosira-antarctica-CCMP982-20140214\|20162_1 | yes |
| *Tx. antarctica** | MMETSP0152, MMETSP0154 | Thalassiothrix-antarctica-L6_D1-20140214\|17297_1 | yes |
| *F. kerguelensis** | MMETSP0906, MMETSP0907, MMETSP0908 MMETSP0909 | Fragilariopsis-kerguelensis-L2_C3-20140214\|35188_1 | yes |
| *T. striata* | MMETSP0817, MMETSP0818, MMETSP0819, MMETSP0820 | Tetraselmis-striata-LANL1001-20140214\|2880_1 | yes |
| | | Tetraselmis-striata-LANL1001-20140214\|11490_1 | yes |
| | | Tetraselmis-striata-LANL1001-20140214\|18675_1 | yes, except in MMETSP0817 |
| | | Tetraselmis-striata-LANL1001-20140214\|95585_1 | yes, except in MMETSP0817 |

*Note: No genome was available for this species, the putative homolog was found in the MMETSP transcriptome data only.

### Table S4. Expression levels of identified DLHs under environmental stress conditions.

| Species | DL homolog | Experiment (stress condition tested) | DL expression - control | DL expression - stress condition tested | DL expression- Fold change | Biological replicates | Source |
|---|---|---|---|---|---|---|---|
| *Azadinium spinosum* | Azadinium spinosum A | Cold stress | 0 | 2710 | - | 1 | MMETSP |
| | Azadinium spinosum B | Cold stress | 203 | 191 | 0.9 | 1 | MMETSP |
| | Azadinium spinosum C | Cold stress | 685 | 702 | 1.0 | 1 | MMETSP |
| | Azadinium spinosum D | Cold stress | 1048 | 1115 | 1.1 | 1 | MMETSP |
| *Amphidinium carterae* | Amphidinium carterae A | High light | 12 | 16 | 1.3 | 1 | MMETSP |
| | Amphidinium carterae B | High light | 0 | 0 | not expressed | 1 | MMETSP |
| | Amphidinium carterae C | High light | 0 | 7 | - | 1 | MMETSP |
| | Amphidinium carterae D | High light | 196 | 199 | 1.0 | 1 | MMETSP |
| | Amphidinium carterae E | High light | 0 | 0 | not expressed | 1 | MMETSP |
| *Isochrysis* | Isochrysis sp. A | High light | 334 | 316 | 0.95 | 1 | MMETSP |
| | Isochrysis sp. B | High light | 194 | 490 | 2.53 | 1 | MMETSP |
| | Isochrysis sp. C | High light | 89 | 33865 | 382.46 | 1 | MMETSP |
| | Isochrysis sp. D | High light | 1597 | 846 | 0.53 | 1 | MMETSP |
| *Thalassiosira antarctica* | T.antarctica A | Cold stress | 203 | 399 | 1.97 | control -2; cold stress-1 | MMETSP |
| | T. antarctica B | Cold stress | 30 | 18 | 0.59 | control -2; cold stress-1 | MMETSP |
| *Thalassiosira antarctica* | T.antarctica A | Si limitation | 440 | 19 | 0.04 | control -2; Si limitation-1 | MMETSP |
| | T. antarctica B | Si limitation | 31 | 0 | 0.00 | control -2; Si limitation-1 | MMETSP |
| *Pseudonitzschia fradulenta* | P, fradulenta A | Si limitation | 0 | 260 | - | 1 | MMETSP |
| | P. fradulenta B | Si limitation | 0 | 568 | - | 1 | MMETSP |
| *Skeletonema dohrnii* | S. dohrnii | N limitation | 73 | 11 | 0.15 | 1 | MMETSP |
| *Seminavis robusta* | S. robusta | N limitation (48 h) | 4 | 2 | 0.54 | 2 | Osuna-Cruz et al., (2020) |
| | S. robusta | N limitation (72 h) | 10 | 1 | 0.10 | 2 | Osuna-Cruz et al., (2020) |
| *Ulva mutabilis* | U. mutabilis A | Cold stress | Not available | Not available | no change | 4 | De Clerck et al., (2018) |
| | U. mutabilis B | Cold stress | Not available | Not available | 0.08 | 8 | De Clerck et al., (2018) |
| *Acropora millepora* | A. millepora A | High CO2 | 3453 | 2919 | 0.85 | 3 | Moya et al., (2012) |
| | A. millepora D | High CO2 | 5850 | 5102 | 0.87 | 3 | Moya et al., (2012) |
| | A. millepora E | High CO2 | 1999 | 1497 | 0.75 | 3 | Moya et al., (2012) |
| | A. millepora F | High CO2 | 6066 | 6202 | 1.02 | 3 | Moya et al., (2012) |
| | A. millepora I | High CO2 | 10976 | 10177 | 0.93 | 3 | Moya et al., (2012) |
| | A. millepora G | High CO2 | 4221 | 3400 | 0.81 | 3 | Moya et al., (2012) |

**Table S5. DL homolog expression level and DMS concentration in surface water.** This is the data used for correlation analysis presented in Fig. 3B in the main text.

| DL expression data | | | | | | | | DMS data | | | | |
|---|---|---|---|---|---|---|---|---|---|---|---|---|
| **Tara station** | **Location** | **Latitude** | **Longitude** | **Sampling date** | **Sample_IDs included** | **Total DL homologs abundance in station** | **Comments** | **DMS (nM)** | **Latitude** | **Longitude** | **Smapling date** | **Reference** |
| Tara_011 | Northwest Mediterranean Sea | 41°66'N | 2°79'E | Oct 2009 | TARA_X000001286<br>TARA_X000001288<br>TARA_X000001334<br>TARA_X000001338 | 7.98E-05 | | 0.9 ± 0.2 | 41°40'N | 2°48'E | winte 2003 | Vila-Costa et al., 2008 |
| Tara_051 | La Réunion island | 21°46'S | 54°30'E | May 2010 | TARA_N000000185<br>TARA_N000000231<br>TARA_N000000196<br>TARA_N000000189 | 1.26E-04 | | 1.4 | 20°S | 56° E | Dec 1997 | Sciare et al., 1998 |
| Tara_082 | Falkland Islands | 47°16'S | 58°01'E | Dec 2010 | TARA_N000001386<br>TARA_N000001390<br>TARA_N000001394<br>TARA_N000001398 | 2.42E-04 | | 3.5 | ~50°S | ~60°E | Dec monthly avraged | Shrivardhan et al., 2022 |
| Tara_084 | Palmer station | 60°39'S | 60°47'W | Jan 2011 | TARA_N000001442<br>TARA_N000001362<br>TARA_N000001438<br>TARA_N000001440 | 8.00E-05 | | 4.1 | 66°54'S | 68°56'W | Jan 1994 | Berresheim et al., 1998 |
| Tara_085 | West Antarctic Peninsula | 62°17'S | 49°50'W | Jan 2011 | TARA_N000001040<br>TARA_N000001017<br>TARA_N000001029 | 3.50E-04 | Fraction 5-20 µm is missing | 23.6 ± 35.3 | 67°57'S | 68°22'W | Jan 2013-17 | Webb et al., 2019 |
| Tara_093 | South Pacific Ocean (near Chile) | 33°76'S | 72°61'W | Mar 2011 | TARA_N000001292<br>TARA_N000001290<br>TARA_N000001288 | 1.79E-05 | Fraction 0.8-5 µm is missing | 0.8 | ~40°S | 80°W | Feb 2000 | Lee et al., 2010 |
| Tara_098 | South Pacific Ocean | 26°26'S | 110°99'W | Feb 2011 | TARA_N000001578<br>TARA_N000001582<br>TARA_N000001586 | 0.000100792 | Fraction 0.8-5 µm is missing | 3.1 | ~35°S | 128°W | Feb 2000 | Lee et al., 2010 |
| Tara_145 | Gulf of Maine | 39°16'N | 70°07'W | Feb 2012 | TARA_N000003219<br>TARA_N000003223<br>TARA_N000003231 | 1.11E-04 | Fraction 0.8-5 µm is missing | 7.5 | ~43°'N | 70°'W | Sept 1994 | Sharma et al., 1999 |
| Tara_148 | Sargasso Sea | 31°78'N | 64°14'W | Feb 2012 | TARA_N000002119<br>TARA_N000002117<br>TARA_N000002115 | 9.74911E-05 | Fraction 0.8-5 µm is missing | 1 | 31°40'N | 64°10'W | Feb 2008 | Levine et al., 2012 |
| Tara_150 | Gulf Stream | 35°80'N | 37°10'W | Mar 2012 | TARA_N000002697<br>TARA_N000002699<br>TARA_N000002701<br>TARA_N000002703 | 1.05E-04 | | 1 | 42°2'N | 45°0'W | May 2003 | Lizotte et al., 2013 |
| Tara_163 | Greenland Sea | 76°07'N | 1°68'E | Jun 2013 | TARA_N010000679<br>TARA_N010000467 | 1.58E-03 | Fractions 0.8-5 and 20-180 µm are missing | 18.3 | 77°23'N | 1°39'E | Jul 2007 | Gali et al., 2010 |
| Tara_194 | Chukchi Sea | 73°33'N | 168°51'W | Sept 2013 | TARA_N010000091<br>TARA_N010000106<br>TARA_N010000110 | 3.24E-05 | Fraction 0.8-5 µm is missing | 2 | ~78°0'N, | 176°97'W, | Aug 2016 | Park et al., 2019 |
| Tara_201 | Devon Island | 74°32'N | 85°72'W | Sept 2013 | TARA_N010000157<br>TARA_N010000166<br>TARA_N010000174 | 4.42E-05 | Fraction 0.8-5 µm is missing | 0.55 | 73°92'N | 86°65'W | Oct 2007 | Luce et al., 2011 |
| Tara_205 | Northern Baffin Bay | 72°42'N | 71°95'W | Oct 2013 | TARA_N010000218<br>TARA_N010000225<br>TARA_N010000229 | 5.43E-05 | Fraction 0.8-5 µm is missing | 0.513 | 72°75'N | 67°00'W | Jul-Aug 2015 | Jarníková et al., 2018 |
| Tara_209 | Southern Baffin Bay | 64°72'N | 53°46'W | Oct 2013 | TARA_N010000308<br>TARA_N010000943<br>TARA_N010000947 | 7.67E-05 | Fraction 0.8-5 µm is missing | 2.006 | 60°45'N | 56°55'W | Jul-Aug 2015 | Jarníková et al., 2018 |
| Tara_210 | Labrador Sea | 61°54'N | 55°98'W | Oct 2013 | TARA_N010000957<br>TARA_N010000962<br>TARA_N010000966 | 8.21E-05 | Fraction 0.8-5 µm is missing | 3.366 | 56°12'N | 53°57'W | Jul-Aug 2015 | Jarníková et al., 2018 |
| Tara_131 | North Pacific Ocean | 22°74'N | 158°05'E | Sept 2011 | TARA_N000002356<br>TARA_N000002364<br>TARA_N000002360<br>TARA_N000002352 | 1.31E-04 | | 4.49 | 23°0"N | 177°48'E | Dec 2011 | Cui et al., 2015 |

**Dataset S1 (separate file).** DMSP lyase homologs.

**Dataset S2 (separate file).** Amino acid sequences of predicted DL homologs.

**Dataset S3 (separate file).** Species with no DL homologs identified in available databases.
